# Multiple Molecular Pathways to Longevity: Opposing Gene Expression Programs Define Distinct Aging Strategies

**DOI:** 10.64898/2025.12.29.696944

**Authors:** Zenith D. Rudich, Jiaxi Guan, Aura A. Tamez González, Grant F. Booth, Sonja K. Soo, Ulrich Anglas, Meeta Mistry, Megan M. Senchuk, Jeremy M. Van Raamsdonk

## Abstract

While aging is the greatest risk factor for the development of neurodegenerative disease, the role of aging in these diseases is poorly understood. Our previous work has shown that targeting aging pathways can be neuroprotective in animal models of neurodegenerative disease. Based on these findings, we believe that by gaining insight into the aging process, that knowledge can be applied to identify novel therapeutic targets for neurodegenerative disease. To advance our understanding of aging, we used a genomics approach to identify genes regulated by multiple lifespan-extending pathways. We performed RNA sequencing on nine long-lived *C. elegans* mutants representing seven longevity pathways: insulin/IGF-1 signaling, dietary restriction, germline deficiency, impaired chemosensation, reduced translation, elevated mitochondrial ROS, and mild mitochondrial impairment. We found that most pairs of long-lived mutants exhibited a significant overlap in differentially expressed genes. Comparing gene expression across the entire panel of long-lived mutants revealed three distinct longevity groups that could be clearly distinguished by gene expression. Interestingly, two of these groups showed modulation of specific genetic pathways in opposite directions, suggesting that there are multiple alternative strategies to achieving long life. Filtering for genes similarly modulated in at least six mutants identified 196 upregulated and 62 downregulated aging genes. Upregulated genes were enriched in immunity, defense and metabolism, while many downregulated genes impacted translation and gene expression. To assess the ability of these genes to enhance longevity individually, we knocked down the commonly upregulated genes in long-lived mutants and evaluated the resulting effect on lifespan. Using this approach, we identified several genes that affect lifespan individually. Upregulation of at least some of these genes was sufficient to enhance stress resistance and extend lifespan in wild-type worms. Overall, the shared longevity genes identified in this work offer potential targets to promote healthy aging and decrease age-onset disease.

## Introduction

Aging is the progressive decline in an organism’s physiological functions that reduces its ability to maintain homeostasis, respond to stress and survive, thereby leading to increased probability of disease and death. Both stochastic and programmed events contribute to aging, and the lifespan of an organism is influenced by both its genetics and the environment. At present, the molecular mechanisms that contribute to longevity are incompletely understood.

To better understand the aging process, researchers have tried to identify genes that can increase lifespan and elucidate how these genes act to promote longevity. This research has greatly benefited from the use of genetic model organisms, primarily yeast, *C. elegans, Drosophila* and mice. Starting first with *C. elegans,* it has been shown in each of these organisms that mutations in a single gene can extend lifespan ^1, 2, 3, 4, 5^. These long-lived genetic mutants have been divided into different pathways of lifespan extension based on which cellular functions are affected by the mutation.

Among the established pathways of longevity are: decreased insulin/IGF-1 signaling (*daf-2* mutants) ^1^; dietary restriction (*eat-2* mutants) ^6^; decreased translation (*ife-2* mutants) ^7, 8^, germline ablation (*glp-1* mutants) ^9^; mild impairment of mitochondrial function (*clk-1, isp-1, nuo-6* mutants) ^10, 11, 12^; decreased chemosensation (*osm-5* mutants) ^13^; and increased mitochondrial superoxide (*sod-2* mutants) ^14^.

In order to gain insight into the molecular mechanisms by which these genetic mutations increase lifespan, we and others have examined gene expression using either microarrays or RNA sequencing (RNA-seq) ^15, 16, 17, 18, 19, 20, 21, 22, 23, 24, 25^. These previous studies have looked at gene expression in individual mutants or small groups of mutants and have used a variety of different experimental paradigms. As a result, it has not been possible to directly compare gene expression across a large panel of long-lived mutants, as it is unclear the extent to which differences observed would be due to differences in the experimental details or genuine differences between the strains.

In exploring the mechanisms underlying lifespan extension in long-lived mutants, it has become clear that at least some mechanisms of lifespan extension are shared across multiple long-lived mutants. For example, the p38-mediated innate immune signaling pathways is required for the extended longevity of *daf-2, clk-1, isp-1, nuo-6, ife-2, eat-2* and *osm-5* mutants ^25, 26, 27^. The mitochondrial unfolded protein response is needed for the long life of *clk-1, isp-1, nuo-6* and *glp-1* mutants ^17, 28^. An increase in reactive oxygen species contributes to the long lifespan of *sod-2, clk-1, isp-1, nuo-6, daf-2,* and *glp-1* mutants ^14, 29, 30, 31^. Disruption of the AMPK gene *aak-2* decreases the long lifespan of *daf-2, clk-1, isp-1* and *glp-1* mutants ^32, 33^. Decreasing the levels of the FOXO transcription factor DAF-16 reduces the long lifespan of several long-lived mutants ^1, 6, 8, 13, 16, 34, 35^. However, it should be noted that the interpretation of these results is dependent on whether pathway disruption also decreases lifespan in wild-type worms and whether there is evidence of pathway activation ^36^.

In this work, we sought to identify unique and shared genetic pathways contributing to lifespan extension in a panel of nine long-lived mutants. We found that there is a highly significant degree of overlap between differentially expressed genes in most pairs of long-lived mutants. Examining gene expression across all nine long-lived mutants reveals three distinct longevity groups. Interestingly, there are specific groups of genes that are significantly modulated in opposite directions in different longevity groups indicating that there are multiple effective strategies to achieve long life. In screening genes that are differentially expressed in multiple long-lived mutants, we find that individual differentially expressed genes can significantly affect both lifespan and resistance to stress.

## Results

### Magnitude of lifespan extension differs between long-lived mutants

In order to gain insight into shared and unique genetic pathways contributing to longevity in a panel of long-lived genetic mutants, we selected nine different genetic mutants to perform transcriptomic analysis using RNA-seq. Before proceeding with RNA-seq, we first wanted to confirm that all of these strains are long-lived and to compare the relative magnitudes of lifespan extension. The lifespan was performed in absence of FUdR as the RNA-seq samples would not be exposed to FUdR. Under these conditions, we confirmed that all nine strains lived significantly longer than wild-type worms (**Figure 1**). Further, we found that *daf-2* worms had the greatest magnitude of lifespan extension followed by *osm-5, nuo-6, isp-1, ife-2, eat-2, sod-2* and *clk-1* worms (**Figure 1**)(*glp-1* worms were not included in this list as the lifespan was performed under different conditions in which worms are developed at 25°C until adulthood). When we previously quantified the lifespan of these strains using 25 µM FUdR, we observed similar increases in lifespan where *daf-2* worms lived the longest and *osm-5, isp-1* and *nuo-6* worms lived longer than *eat-2, sod-2, clk-1* and *ife-2* worms ^27^.

**Figure 1.**
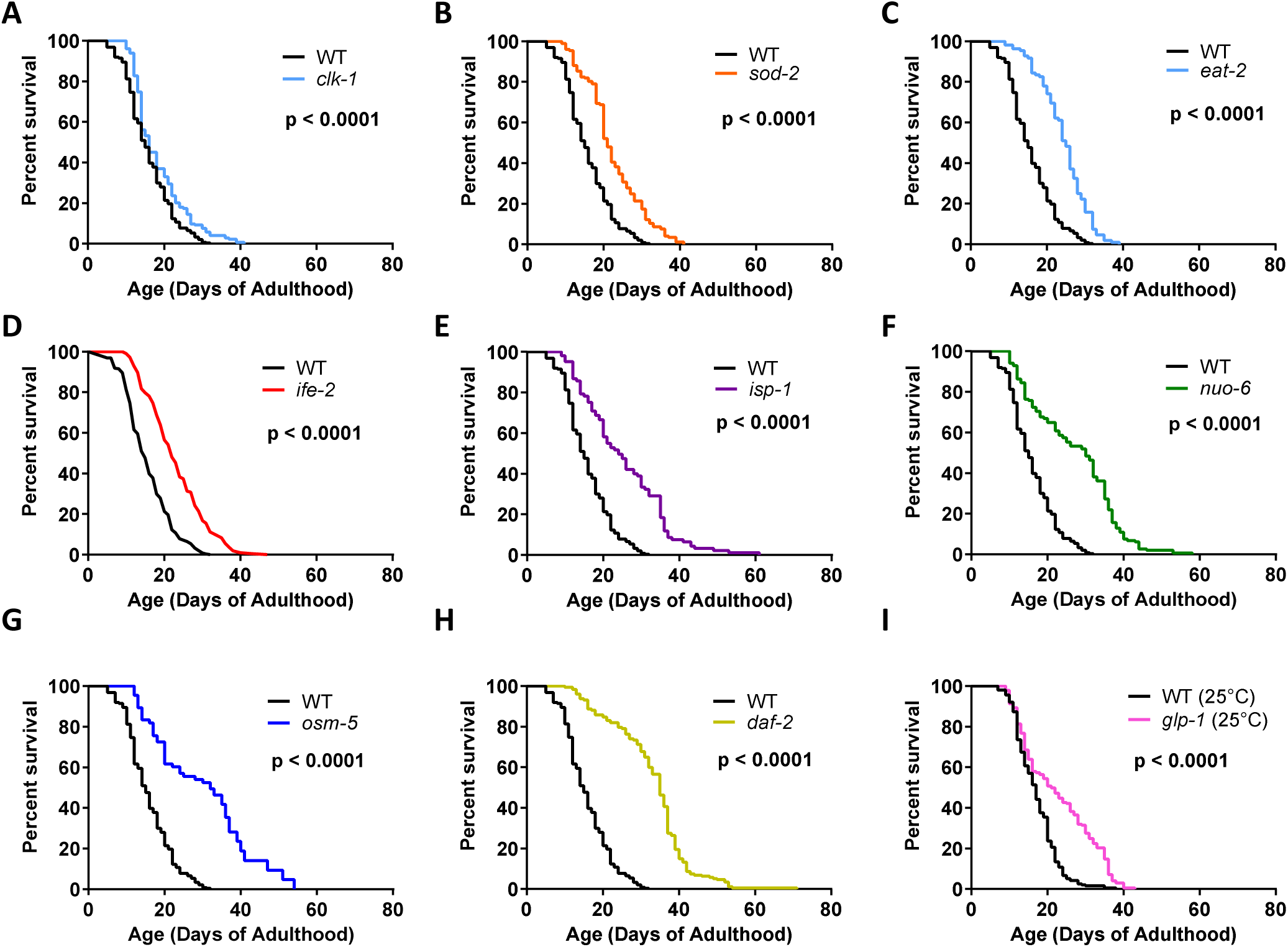
Long-lived mutants exhibit significantly extended lifespans of varying magnitudes. The lifespan of nine different long-lived mutants was measured on NGM plates with no FUdR. All of the long-lived mutants exhibited significantly increased lifespan of varying magnitudes. The long-lived mutants included *clk-1* (**A**)*, sod-2* (**B**)*, eat-2* (**C**)*, ife-2* (**D**)*, isp-1* (**E**)*, nuo-6* (**F**)*, osm-5* (**G**)*, daf-2* (**H**) and *glp-1* (**I**) mutants. *glp-1* worms were allowed to develop at 25°C then shifted to 20°C at adulthood. A minimum of three biological replicates were performed for each strain. Significance was assessed using the log-rank test. Three biological replicates were performed. Total n for each strain was as follows: N2 (244), *clk-1* (174), *sod-2* (174), *eat-2* (108), *ife-2* (214), *isp-1* (94), *nuo-6* (148), *osm-5* (48), *daf-2* (176), *glp-1* (197), *and* N2 - 25°C (195).

Having shown that all nine genetic mutants are long-lived, both with and without FUdR, we proceeded to analyze their gene expression by performing RNA-seq on pre-fertile young adult worms. We used a minimum of six biological replicates per strain. Principal component analysis (PCA) for each long-lived mutant demonstrated that all of the long-lived mutants could be clearly distinguished from wild-type worms based on gene expression (**Figure S1-S9**). We found that there are hundreds to thousands of differentially expressed genes in each of the long-lived mutant strains (**Figures S1-9**). In all strains, we observed both significantly upregulated and significantly downregulated genes (**Figures S1-S9**). Lists of differentially expressed genes from each long-lived mutant (**Table S1**) were used for enrichment analysis to identify pathways and functions that are overrepresented in each of the long-lived mutants individually (**Figure S10-18**).

We have compared the differentially expressed genes identified in this study to previous gene expression studies involving these long-lived mutant strains. In each case, we observed a highly significant degree of overlap (**Figure S19-S25**). We have provided a list of the overlapping genes between our study and previous studies in **Table S2**.

### Expression of specific single genes at young adulthood is highly correlated with longevity

Having identified thousands of genes that are differentially expressed in long-lived mutants, we next sought to determine the extent to which expression of individual differentially expressed genes is associated with longevity. To do this, we compared the expression levels of each individual gene to their lifespan with and without FUdR. Using this analysis, we found that there are 853 genes whose expression levels are positively correlated with longevity measured in absence of FUdR, and 1459 genes whose expression levels are positively correlated with longevity when FUdR is present (**Figure 2A; Table S3**). Of these, there were 753 genes in common. In contrast, there were very few genes whose expression levels are negatively correlated with longevity (45 in absence of FUdR, 149 with FUdR; **Figure 2B; Table S3**). Individual genes exhibited R-squared values as high as 0.901 (p=0.0004) for positively correlated genes (**Figure 2C**) or 0.833 (p=0.0028) for negatively correlated genes (**Figure 2D**). Combined, this suggests that the expression level of individual genes at young adulthood can be predictive of lifespan.

**Figure 2.**
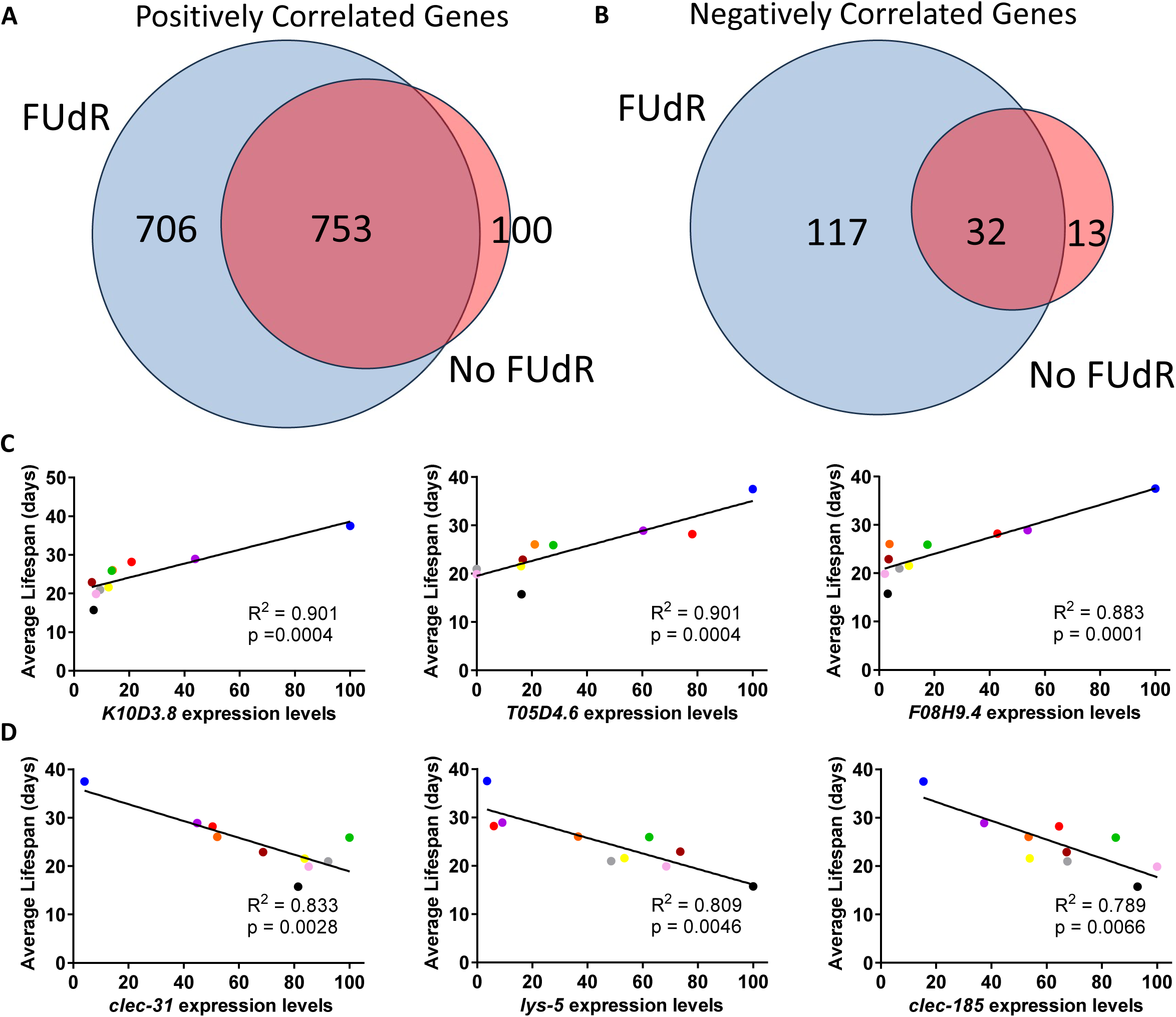
Gene expression levels during young adulthood are highly correlated with lifespan. (**A**) Throughout the genome there were 853 genes positively correlated with lifespan on plates containing no FUdR and 1459 genes positively correlated with lifespan on plates with 25 µM FUdR. Of these genes, there were 753 genes positively correlated with lifespan under both condition. (**B**) There were much fewer genes that were negatively correlated with lifespan: 45 genes were negatively correlated with lifespan on plates containing no FUdR and 149 genes were negatively correlated with lifespan on plates with 25 µM FUdR. There were 32 genes negatively correlated with lifespan under both conditions. (**C**) Examples of genes showing a high positive correlation with lifespan. (**D**) Examples of genes showing a high negative correlation with lifespan. Combined, these results suggest that gene expression at young adulthood can be predictive of lifespan. For C and D, blue = *daf-2,* purple = *isp-1,* red = *nuo-6*, orange = *osm-5,* green = *glp-1*, brown = *eat-2,* grey = *clk-1,* yellow = *sod-2* and pink = *ife-2*.

### Highly significant degree of overlap in differentially expressed genes between pairs of long-lived mutants

We next compared differentially expressed genes between pairs of long-lived mutants to determine the extent to which each mutant exhibited overlap with the other mutants and assess which long-lived mutants are most similar in gene expression. To determine the significance of each overlap, we compared the observed number of overlapping genes to the number of overlapping genes expected if two similarly sized gene sets were selected completely at random. We found that all of the long-lived mutants show a statistically significant overlap with multiple other long-lived mutants for both upregulated (**Figure 3; Figure S26**) and downregulated genes (**Figure S26,S27**). While genes differentially expressed in *daf-2* and *ife-2* worms exhibited a significant overlap with genes differentially expressed in all other long-lived mutants, genes differentially expressed in *eat-2* worms showed relatively little overlap with other long-lived mutants (**Figure 3; Figure S26,S27**). Combined, these results suggest that there are common genetic pathways contributing to longevity shared across multiple long-lived mutants, but that some long-lived mutants have relatively unique changes in gene expression. This may represent the existence of different strategies to achieve long life.

**Figure 3.**
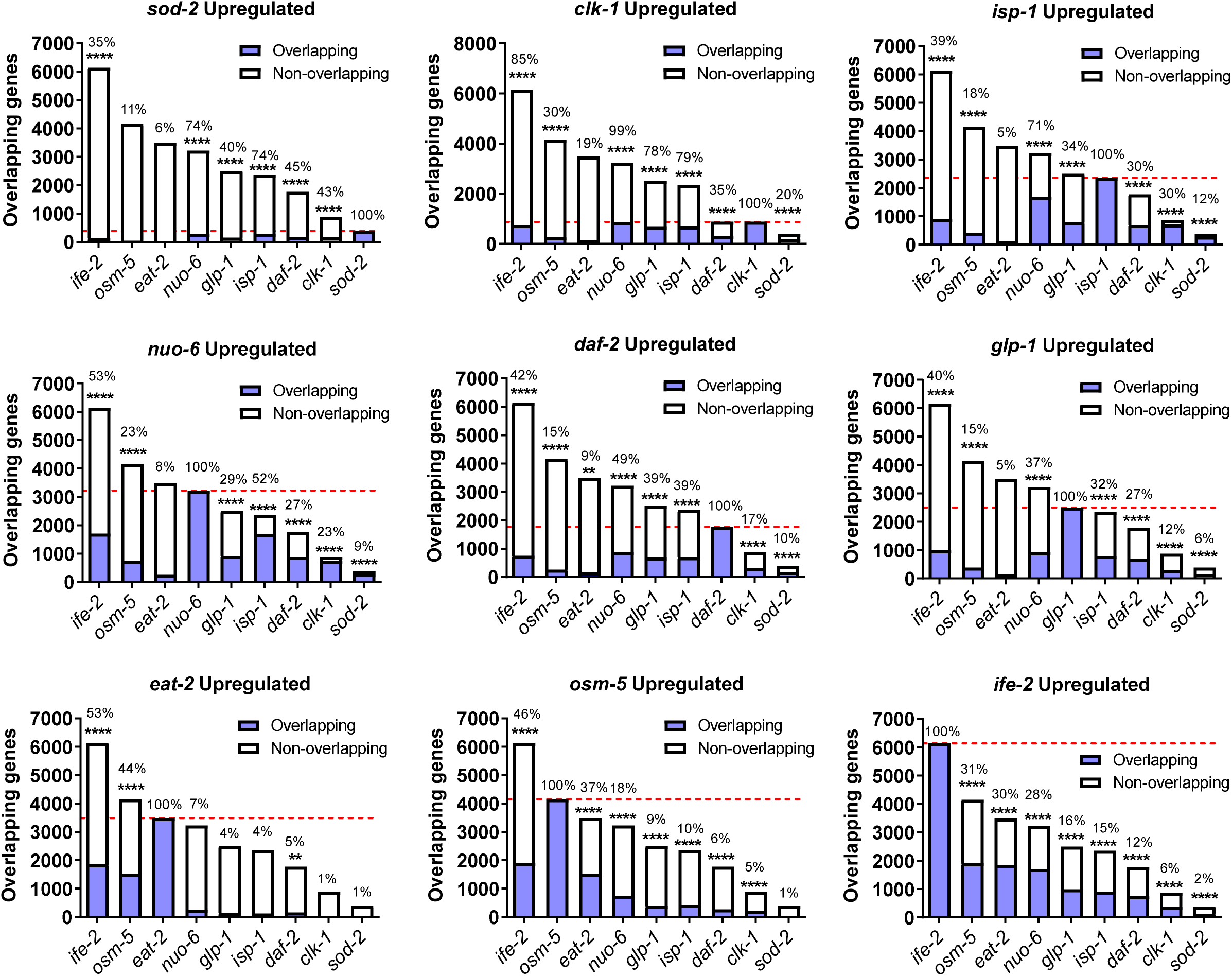
Degree of overlap of differentially expressed genes between pairs of long-lived mutants is highly significant. Genes upregulated in each of the nine different long-lived mutants were compared to genes upregulated in the other eight long-lived mutants in a pairwise manner. All of the long-lived mutants showed a significant degree of overlap with multiple other long-lived mutants suggesting that shared common pathways are promoting longevity. *eat-2* mutants showed the least overlap with other long-lived mutants suggesting that the pathways promoting longevity in this mutant are relatively unique compared to other long-lived mutants. The percentage shown above each bar indicates the number of overlapping genes as a percentage of the total number of upregulated genes for the strain indicated in the graph title on top of each graph. Statistical significance was assessed using a Fisher’s exact test. **p<0.01, ***p<0.001, ****p<0.0001.

### Long-lived genetic mutants segregate into three distinct longevity groups based on gene expression

Having observed a significant degree of overlap between differentially expressed genes between pairs of long-lived mutants, we next looked for overlaps across differentially expressed genes from multiple long-lived mutants. Heatmaps comparing gene expression across all of the long-lived mutants reveal that the long-lived mutants cluster into three distinct longevity groups (**Figure 4A,B; Figure S28**). The largest group, which we will refer to as group 1, contains *daf-2, nuo-6, isp-1, clk-1, glp-1* and *sod-2* worms. The second group, which we will refer to as group 2, contains *osm-5* and *eat-2* mutants. Finally, *ife-2* worms exhibit large numbers of both upregulated and downregulated genes which show partial overlap with both group 1 and group 2 genes.

**Figure 4.**
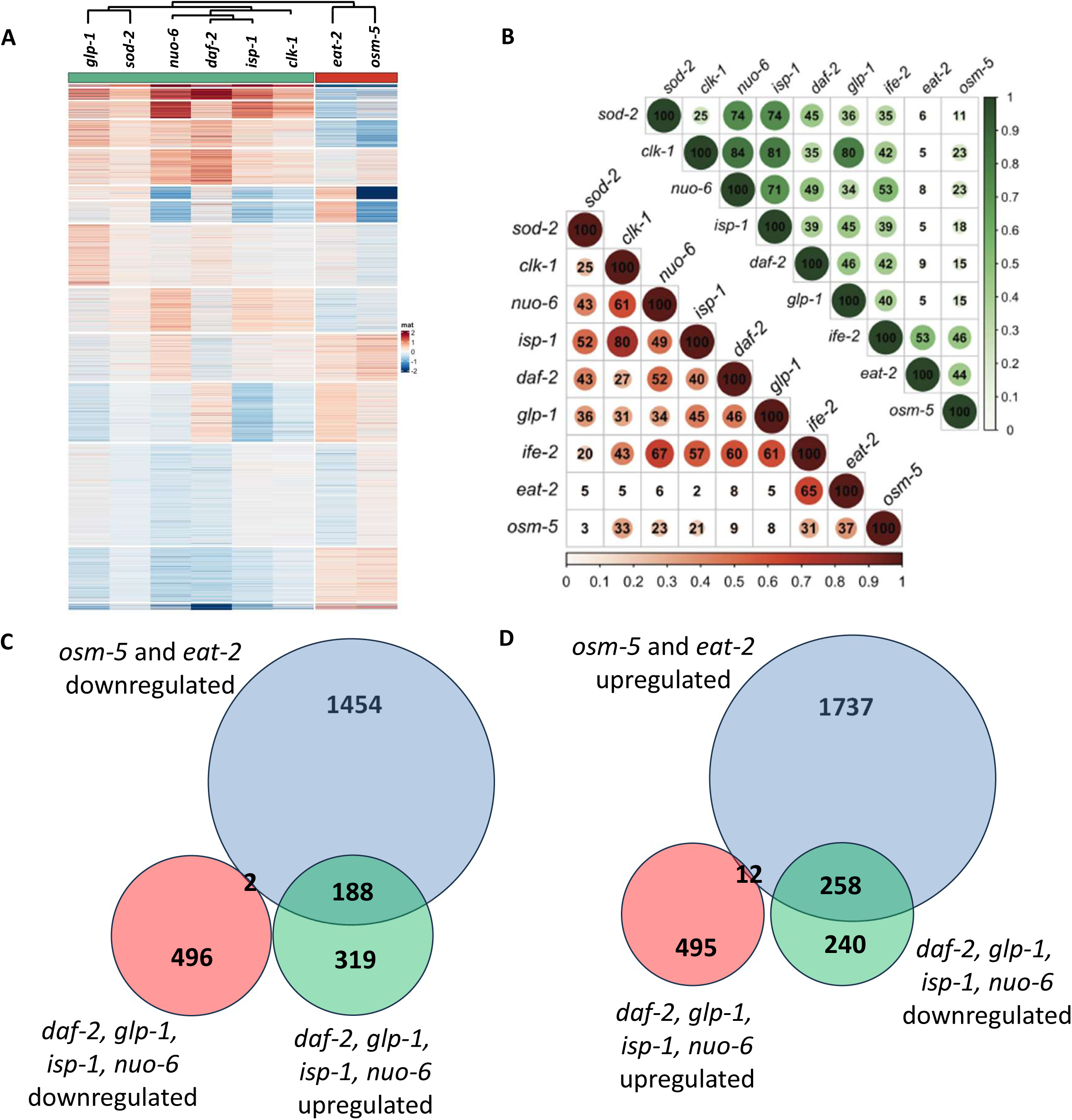
Long-lived mutants segregate into three distinct longevity groups based on gene expression. **A.** A heat map comparing eight of the nine-long lived mutants indicates that the mutants cluster into two groups based on gene expression. Group 1 contains *daf-2, nuo-6, isp-1, glp-1, clk-1* and *sod-2* mutants. Group 2 contains *osm-5* and *eat-2* mutants. **B.** Diagram showing the percentage overlap of upregulated and downregulated genes in the nine long-lived mutants. The group 2 strains *eat-2* and *osm-5* show little overlap with group 1 strains. *ife-2* worms exhibit overlap with both group 1 and group 2 strains. **C.** Genes that are downregulated in group 2 worms (*eat-2, osm-5*) show a much greater overlap with genes upregulated in group 1 worms (*daf-2, nuo-6, isp-1, glp-1*) than with genes downregulated in group 1 worms. **D.** Similarly, genes upregulated in group 2 worms show a much greater overlap with genes downregulated in group 1 worms than with genes upregulated in group 1 worms. Thus, for many genes group 1 and group 2 worms exhibit changes in gene expression in the opposite direction.

Interestingly, some groups of genes upregulated in group 1 were found to be downregulated in group 2, while some groups of genes downregulated in group 1 were found to be upregulated in group 2 (**Figure 4C,D**). For example, of the 507 genes commonly upregulated in the group 1 mutants *daf-2, isp-1, nuo-6* and *glp-1,* 188 of these genes are downregulated in group 2 mutants *osm-5* and *eat-2,* while only 12 are also upregulated in group 2. Of the 498 genes that are commonly downregulated in the group 1 mutants *daf-2, isp-1, nuo-6* and *glp-1,* 258 of these genes are upregulated in group 2 mutants *osm-5* and *eat-2*, while only 2 are also downregulated in group 2. This suggests that group 1 and group 2 longevity mutants may be utilizing different strategies to live long, and that genetic pathways that affect longevity can be modulated in different ways to achieve long life.

To gain further insight into the genetic pathways that are modulated in different directions between group 1 and group 2 mutants, we examined the expression of individual genes from specific pathways known to affect longevity. We and others have demonstrated that the FOXO transcription factor DAF-16 contributes to lifespan extension in *daf-2* worms, as well as other group 1 mutants including *nuo-6, isp-1, clk-1* and *glp-1* ^1, 16, 37^. In examining the expression of high confidence DAF-16 target genes ^38^, we found that genes that are activated by DAF-16 (Class I target genes) are upregulated in most of the group 1 longevity mutants, but either unchanged or decreased in group 2 mutants (**Figure 5**). Similarly, genes that are repressed by DAF-16 (Class II DAF-16 target genes) were found to be downregulated in group 1 longevity mutants but not in group 2 or group 3 mutants (**Figure S29**).

**Figure 5.**
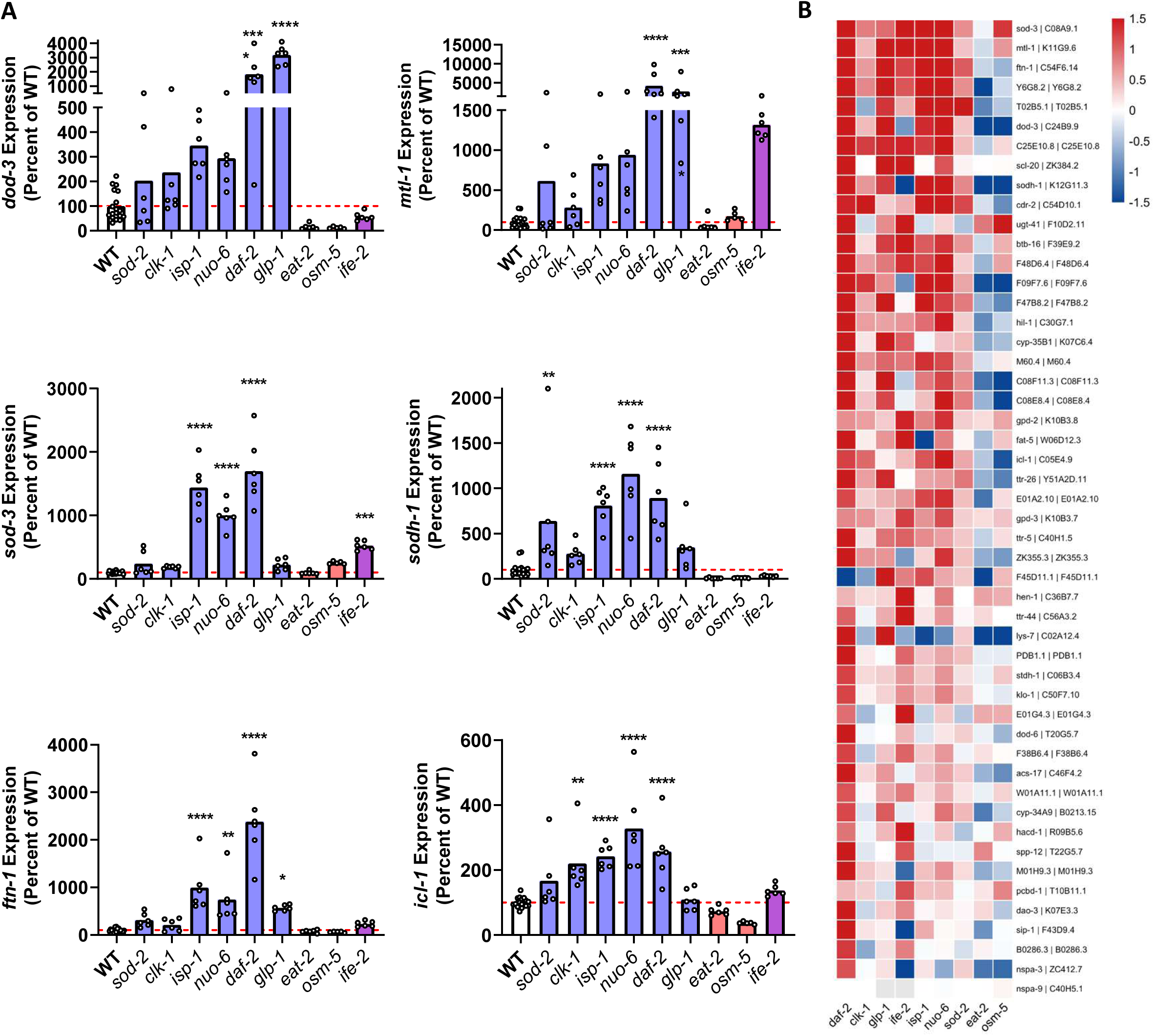
Group 1 longevity mutants exhibit upregulation of Class I DAF-16 target genes. **A.** Examination of expression of Class I DAF-16 target genes (genes activated by DAF-16) across group 1 (blue bars), group 2 (red bars) and group 3 (purple bars) longevity mutants reveals that DAF-16 target genes are upregulated in group 1 mutants but either unchanged or downregulated in group 2 mutants. **B.** A heat map of the top 50 high confidence DAF-16 target genes from Tepper et al. *Cell* 2014 shows activation of the DAF-16 stress response pathway in group 1 and group 3 mutants but not in group 2 mutants. These results suggest that group 1 longevity mutants rely on the DAF-16 stress response pathway to promote longevity while group 2 mutants utilize other pathways. Panel A includes RNA-seq data from six biological replicates per strain except for wild-type, which included 18 biological replicates. Statistical significance was assessed using a one-way ANOVA with Dunnet’s multiple comparisons test. **p<0.01, ***p<0.001, ****p<0.0001.

We and others have also shown that the mitochondrial unfolded protein response (mitoUPR) is required for the extended longevity of group 1 longevity mutants *nuo-6, isp-1, clk-1* and *glp-1* ^17, 28^. As with the DAF-16 target genes, we found that mitoUPR target genes are upregulated in most of the group 1 longevity mutants, but downregulated or unchanged in group 2 longevity mutants (**Figure 6**). At the same time, there were several genes that are significantly upregulated in group 2 longevity mutants but downregulated in group 1 longevity mutants (**Figure S30**). This included the PQM-1 target genes *F55G11.2* and *fat-7* ^38^. Combined, this indicates that genetic pathways that affect longevity can be modulated in opposite directions to achieve lifespan extension (**Table S4** provides a list of genes that are significantly upregulated in group 1 worms (*daf-2, glp-1, isp-1, nuo-6, clk-1,* and *sod-2* worms) but downregulated in group 2 worms (*osm-5* and *eat-2* worms), and vice versa).

**Figure 6.**
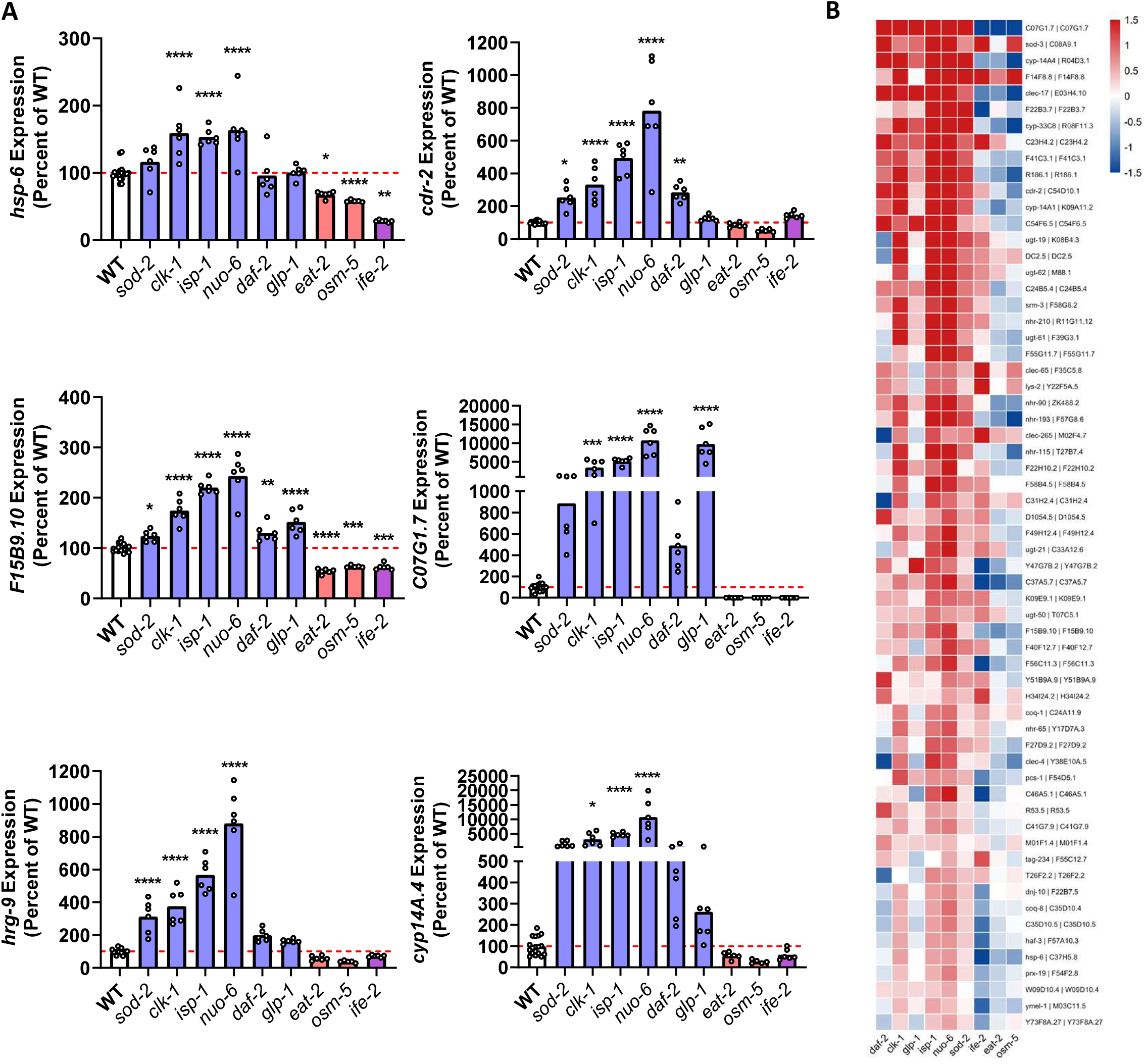
Group 1 longevity mutants exhibit upregulation of ATFS-1 target genes. **A.** Examination of expression of ATFS-1 target genes across group 1 (blue bars), group 2 (red bars) and group 3 (purple bars) longevity mutants reveals that ATFS-1 target genes are upregulated in group 1 mutants but either unchanged or downregulated in group 2 and group 3 mutants. **B.** A heat map of the top 61 high confidence ATFS-1 target genes from Soo et al. *Micropublication Biology* 2021 shows activation of the mitochondrial unfolded protein response (mitoUPR) pathway in group 1 mutants but not in group 2 or group 3 mutants. These results suggest that group 1 longevity mutants rely on the mitoUPR pathway to promote longevity while group 2 and group 3 mutants utilize other pathways. Panel A includes RNA-seq data from six biological replicates per strain except for wild-type, which included 18 biological replicates. Statistical significance was assessed using a one-way ANOVA with Dunnet’s multiple comparisons test. **p<0.01, ***p<0.001, ****p<0.0001.

To gain further insight into the pathways contributing to longevity in the long-lived mutants and how these differ between the three longevity groups, we performed transcription factor analysis on the RNA-seq data to determine which transcription factors might be driving the longevity-associated transcriptional changes. To do this we used two complementary approaches: (1) transcription factor inference, which is based on the coordinated expression changes of known transcription factors; and (2) motif enrichment analysis, which is based on identifying transcription factor binding motifs in the promoters of differentially expressed genes. After identifying which transcription factors were identified for each individual mutant (**Table S5**), we then compared the identified transcription factors across all nine mutants.

Interestingly, while 33 of the same transcription factors were implicated in group 1 and group 2 longevity mutants, 25 are modulated in different directions (activated in group 1, repressed in group 2 or vice versa) while only 5 are modulated in the same direction (**Figure S31**). This indicates that although group 1 and group 2 longevity mutants may modulate overlapping pathways to achieve long lifespan, in most cases these pathways are modulated in opposite directions.

Previous research has shown that aging is associated with widespread changes in gene expression^39^. Accordingly, we compared gene expression in the long-lived mutants to gene expression changes that take place during aging. We found that all of the long-lived mutants exhibit changes in gene expression that were both the same direction as changes that take place during aging (e.g. upregulated in long-lived mutant and upregulated during aging or downregulated in long-lived mutant and downregulated during aging) and genes that were modulated in opposite direction as changes that take place during aging (upregulated in long-lived mutant and downregulated during aging or downregulated in long-lived mutant and upregulated during aging (**Table S6**). Interestingly, group 1 longevity mutants exhibited a greater number of transcriptional changes that were in the same direction as genes that are modulated during aging, while group 2 longevity mutants exhibit a greater number of changes that are in the opposite direction as genes that are modulated during aging (**Figure S32**). *ife-2* mutants (group 3) exhibited a similar number of differentially expressed genes that were in the same direction and opposite direction as genes that are modulated during aging.

### Genes that are commonly upregulated in the majority of long-lived mutants are involved in innate immunity, defense and metabolism

We next sought to determine the extent to which there are genetic pathways contributing to longevity that are shared across all or many of the long-lived mutants. We found that there were little or no genes that were differentially expressed in the same directions across seven or more long-lived mutants (**Figure 7A, Table S7**). There were 196 genes that were found to be significantly upregulated in six or more long-lived mutants. There were 362 genes that were found to be downregulated in five or more long-lived mutants. We performed Gene Ontology (GO) term and KEGG pathway enrichment on these sets of overlapping genes. Among the genes upregulated in seven or more long-lived mutants, there was an overrepresentation of genes involved in innate immunity, defense and metabolism (**Figure 7B, Figure S33**). This is consistent with our previous work highlighting the contribution of innate immune signaling and pathways of cellular resilience to longevity ^16, 17, 26, 27, 36^. Genes downregulated in five or more long-lived mutants included genes involved in translation, gene expression, ribosome function and metabolism (**Figure 7C, Figure S34**). This is consistent with existing work showing that decreasing translation can extend lifespan ^7, 8, 40^.

**Figure 7.**
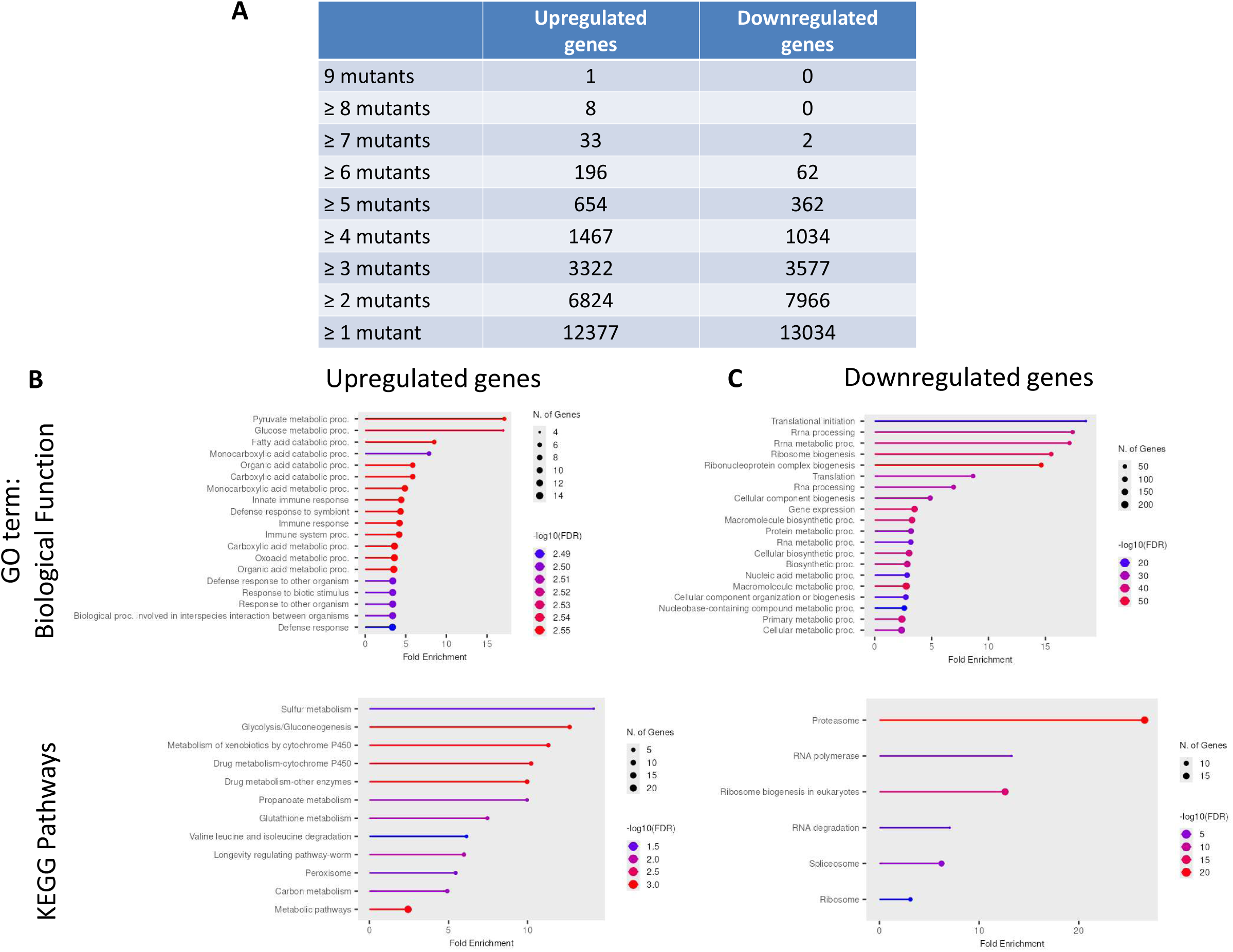
Genes that are commonly upregulated in the majority of long-lived mutants are involved in metabolism, defense and innate immunity. **A.** Comparing differentially expressed genes across the panel of long-lived mutants revealed that very few genes are commonly regulated across seven or more strains. **B.** Among the 196 genes that are upregulated in 6 or more long-lived mutants there was an enrichment for genes involved in various metabolic processes, defense response and innate immunity. **C.** Among the 362 genes that are downregulated in 5 or more long-lived mutants there was an enrichment for genes involved in translation, ribosome and gene expression.

### Individual genes that are modulated in the majority of long-lived mutants are required for lifespan extension and resistance to stress

Having successfully identified genetic pathways contributing to longevity across multiple long-lived mutants, we next determined the extent to which individual differentially expressed genes contributed to lifespan extension in the long-lived mutants. To do this, we focused on the 196 genes that are significantly upregulated in six or more long-lived mutants. We rationalized that the upregulation of these genes could either be contributing to longevity, associated with longevity or could be acting to limit longevity. To distinguish between these possibilities, we planned to decrease the expression of these upregulated genes using RNA interference (RNAi) and then measure lifespan. Of these 196 genes, we tested 116 RNAi clones which were (1) in the Ahringer RNAi library; and (2) grew after one or two attempts. We decided to knock down these genes in either *daf-2* or *nuo-6* mutants, depending on which strain exhibited the greatest magnitude of upregulation, as these two strains provide a large window to see the effects of knockdown. We included *daf-16* RNAi as a positive control in each experiment.

Among the genes tested in *daf-2* mutants, we found that the majority of RNAi clones did not affect *daf-2* lifespan but some RNAi clones were found that modestly increased or decreased longevity (**Figure S35**). For those RNAi clones that decreased lifespan, the magnitude of decrease was always much less than *daf-16* RNAi. Similarly, we found that most of the RNAi clones targeting commonly upregulated genes in *nuo-6* mutants had no effect on lifespan, but that there were clones that both increased and decreased *nuo-6* longevity (**Figure S36**). In order to compare clones across the different lifespan trials, we calculated a weighted mortality score for each gene. This score was calculated as the mortality for the RNAi clone minus the mortality of empty vector all divided by the mortality of *daf-16* RNAi minus the mortality of empty vector (**Figure S37**). Based on these mortality scores, we selected seven genes for confirmation in *nuo-6* mutants and testing in wild-type: C08F11.7, *ugt-62,* K05C4.9, DC2.5, C05B5.5, T07C4.5 and W03B1.7. We found that RNAi targeting all of these genes decreased lifespan in both *nuo-6* and wild-type worms (**Figure 8B,C; Figure S38**). This suggests that these genes are generally important for longevity and not specifically required for the extended longevity of *nuo-6* worms.

**Figure 8.**
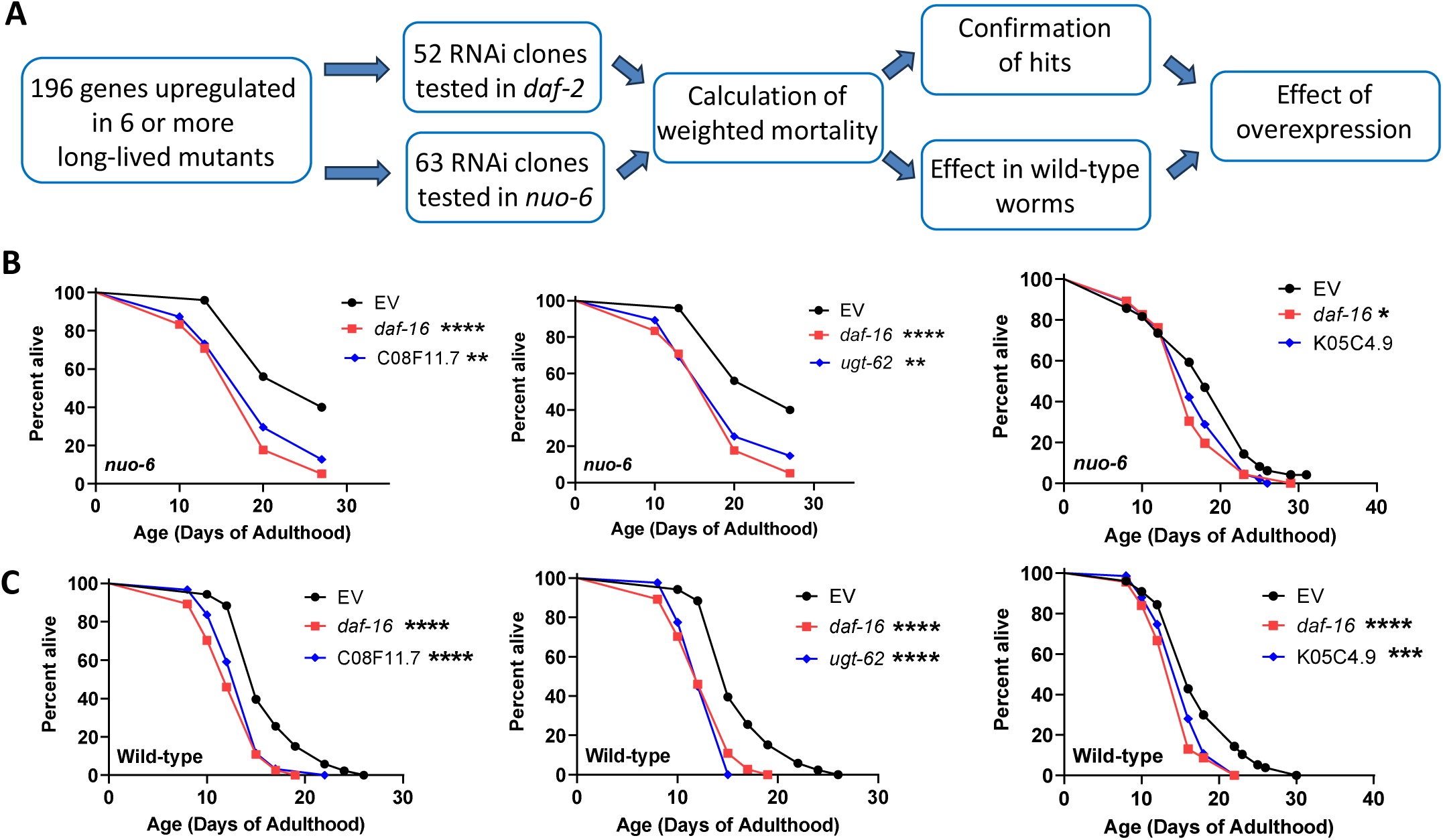
Targeted RNA interference screen identifies differentially expressed genes that can individually affect longevity. To identify differentially expressed genes that can affect lifespan individually, genes that were found to be upregulated in six or more long-lived mutants were knocked down in *daf-2* or *nuo-6* mutants and the resulting effect on lifespan was quantified. While the majority of these RNAi clones did not affect longevity, we found that there were RNAi clones that both increased and decreased lifespan. Three genes were selected to study further: C08F11.7, *ugt-62* and K05C4.9. **(B)** We confirmed that these genes decrease lifespan in *nuo-6* worms. **(C)** Knocking down these genes was also found to decreased wild-type lifespan. Total n per treatment was as follows: *nuo-6* EV (47), *nuo-6 daf-16* RNAi (91), *nuo-6* C08F11.7 RNAi (62), *nuo-6 ugt-62* RNAi (64), *nuo-6* K05C4.9 RNAi (45), WT EV (86), WT *daf-16* RNAi (69), WT C08F11.7 RNAi (61), WT *ugt-62* RNAi (40), WT K05C4.9 RNAi (75). Statistical significance was assessed using the log-rank test.

For further characterization, we decided to focus on three genes. These genes were selected based on the strength and reproducibility of the increase in weighted mortality in order to identify genes with a clear, consistent impact. C05B5.5 and T07C4.5 were ruled out because they had an inconsistent impact on weighted mortality. W03B1.7 was ruled out because it did not have a strong enough effect on weighted mortality. Finally, C08F11.7, *ugt-62,* and K05C4.9 were chosen as they had a greater impact on weighted mortality than DC2.5.

First, we examined the expression of these three genes across the panel of the long-lived mutants. We found that all three of these genes are upregulated in group 1 longevity mutants but not in group 2 or group 3 (**Figure 9A**). Based on this, we wondered if these genes are more important for the lifespan of group 1 longevity mutants than group 2 or group 3. Accordingly, we knocked down these genes in *daf-2* (group 1), *eat-2* (group 2) and *ife-2* (group 3) worms and measured lifespan. Surprisingly, we found that RNAi targeting each of these genes did not decrease *daf-2* lifespan (**Figure 9B**). In contrast, RNAi targeting C08F11.7 or *ugt-62* decreased the lifespan of both *eat-2* worms and *ife-2* worms, while RNAi targeting K05C4.9 significantly decreased *ife-2* lifespan and resulted in a trend towards decreased lifespan in *eat-2* worms (**Figure 9C,D**). Combined, this suggests that C08F11.7, *ugt-62,* and K05C4.9 are generally affecting longevity and not specifically important for group 1 mutants.

**Figure 9.**
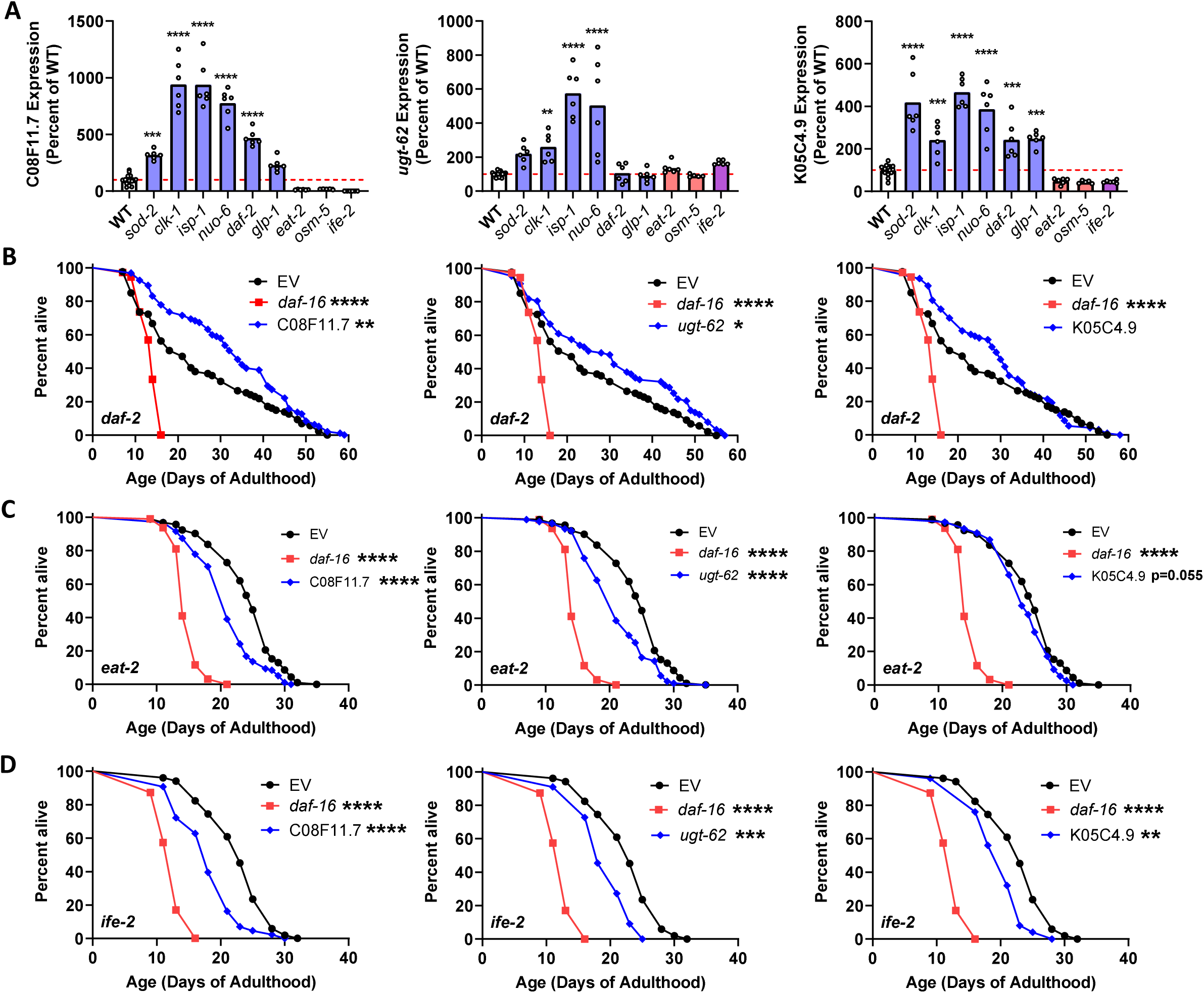
Knockdown of C08F11.7, *ugt-62* and K05C4.9 decreases lifespan in multiple long-lived mutants. **(A)** The expression of C08F11.7, *ugt-62* and K05C4.9 is increased in group 1 longevity mutants, but either unchanged or decreased in group 2 and group 3 mutants. **(B)** Knockdown of C08F11.7, *ugt-62* or K05C4.9 did not decrease the longevity of group 1 *daf-2* mutants. **(C)** Knockdown of C08F11.7 or *ugt-62* but not K05C4.9 decreased lifespan in group 2 *eat-2* mutants. **(D)** Knockdown of C08F11.7, *ugt-62* or K05C4.9 decreased the longevity of group 3 *ife-2* mutants. 25 µM FUdR was used in the lifespan studies. Two biological replicates were performed. The total n for each group was as follows: *daf-2* EV (87), *daf-2 daf-16* RNAi (72), *daf-2* C08F11.7 RNAi (95), *daf-2 ugt-62* RNAi (87), *daf-2* K05C4.9 RNAi (93), *eat-2* EV (92), *eat-2 daf-16* RNAi (95), *eat-2* C08F11.7 RNAi (95), *eat-2 ugt-62* RNAi (91), *eat-2* K05C4.9 RNAi (76), *ife-2* EV (92), *ife-2 daf-16* RNAi (106), *ife-2* C08F11.7 RNAi (81), *ife-2 ugt-62* RNAi (56), *ife-2* K05C4.9 RNAi (64). Statistical significance was assessed using a one-way ANOVA with Dunnett’s multiple comparisons test in panel A and a log-rank test in panels B-D.

### Overexpression of individual upregulated genes can increase stress resistance and lifespan

Having shown that knockdown of C08F11.7, *ugt-62,* and K05C4.9 can decrease lifespan, we next sought to determine the extent to which increasing expression of these genes could increase longevity. Accordingly, we generated novel strains that ubiquitously overexpress C08F11.7, *ugt-62,* and K05C4.9 under the *rpl-28, rpl-2* and *eft-3* promoters, respectively (these strains will be referred to as C08F11.7 OE, *ugt-62* OE, and K05C4.9 OE worms). We used different ubiquitous promoters for each strain so that it would be possible to combine the strains together without transcription factor dilution. In measuring the lifespan of these strains, we found that overexpression of C08F11.7 significantly increased lifespan (**Figure 10A**), while *ugt-62* overexpression caused a small decrease in lifespan (**Figure 10B**), and K05C4.9 overexpression decreased lifespan (**Figure 10C**). Interestingly, worms overexpressing both C08F11.7 and *ugt-62* together exhibited a larger increase in lifespan than C08F11.7 overexpression worms alone (**Figure 10D**).

**Figure 10.**
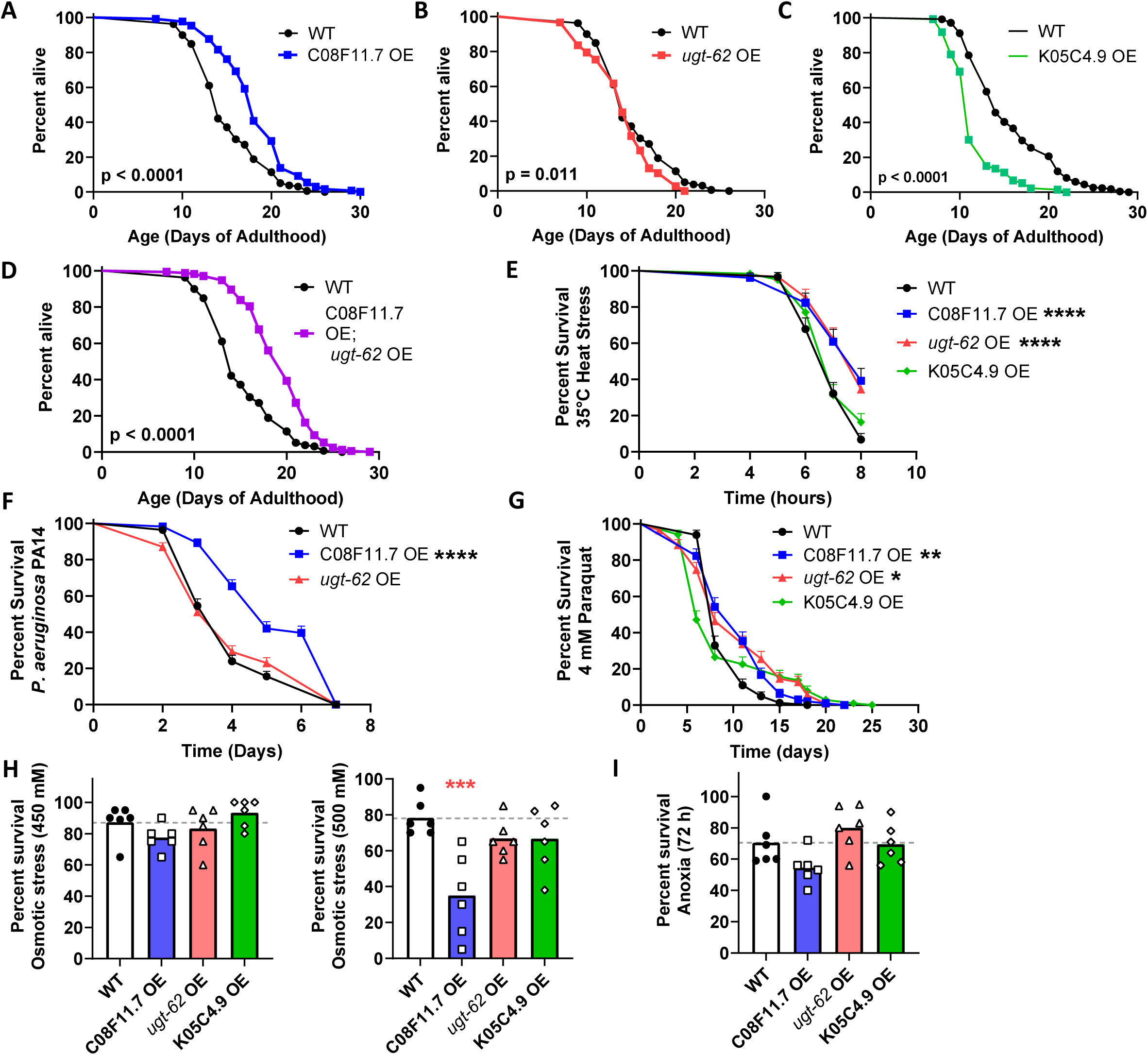
Overexpression of individual genes that are differentially expressed in long-lived mutant strains can increase lifespan and resistance to stress. (**A**) Overexpression (OE) of C08F11.7 significantly increased lifespan, while overexpression of *ugt-62* (**B**) or K05C4.9 (**C**) decreased longevity. (**D**) Overexpression of C08F11.7 and *ugt-62* combined increased lifespan to a greater extent than either gene alone. (**E**) Resistance to heat stress (35°C) was enhanced in worms overexpressing C08F11.7 or *ugt-62.* (**F**) C08F11.7 OE worms also showed enhanced resistance to bacterial pathogens (*P. aeruginosa* strain PA14). (**G**) Resistance to oxidative stress (4 mM paraquat) was increased in C07F11.7 OE and *ugt-62* OE worms. There was no increase in resistance to osmotic stress (450 mM, 500 mM; **H**) or anoxia (72 h; **I**) in any of the overexpression strains. Three biological replicates were performed for each lifespan study. The total number of animals per strain for lifespan studies was as follows: WT (159), C08F11.7 OE (130), *ugt-62* OE (146), K05C4.9 OE (133), C08F11.7 OE; *ugt-62* OE (173). Statistical significance was assessed with a log-rank test in panels A-G and a one-way ANOVA with Dunnett’s multiple comparison’s test in panels H and I. ***p<0.001. The full genotypes of the overexpression strains are: *sybIs6868[rpl-28p::C08F11.7, myo-2p::mCherry], sybIs6945[rpl-2p::ugt-62, myo-3p::mCherry]* and*, sybIs6925[eft-3p::K05C4.9, ges-1p::mCherry]*.

Next, we decided to determine the extent to which overexpression of these three genes could enhance resistance to stress ^41^. We found that both C08F11.7 OE and *ugt-62* OE worms have increased resistance to heat stress (35°C) compared to wild-type worms, while heat stress resistance was unaffected by overexpression of K05C4.9 (**Figure 10E**). In examining bacterial pathogen resistance, we found that C08F11.7 OE worms exhibit increased survival when exposed to *P. aeruginosa* strain PA14, while *ugt-62* OE worms survive as long as wild-type worms (**Figure 10F**). Overexpression of C08F11.7 or *ugt-62* both resulted in increased resistance to oxidative stress (4 mM paraquat; **Figure 10G**). In contrast, none of the overexpression strains exhibited increased survival when exposed to osmotic stress (450 mM NaCl, 500 mM NaCl; **Figure 10H**) or anoxia (72 hours; **Figure 10I**). Combined, this indicates that overexpression of C08F11.7 increases both stress resistance and lifespan while overexpression of *ugt-62* increases resistance to stress.

## Discussion

In this work, we compared gene expression changes across a panel of nine long-lived mutants. We found that these mutants exhibit significant overlap among their differentially expressed genes, but that there appear to be both common and opposing strategies to achieve long life. Genes involved in metabolism, immunity, translation and stress resilience were found to be central regulators of longevity. Individual genes were found to be predictive of lifespan and to be able to affect both lifespan and stress resistance individually.

This research extends previous single mutant transcriptomic studies by analyzing a panel of long-lived mutants under the same experimental conditions in order to facilitate direct comparison by reducing technical variation. We found that there are hundreds of genes whose expression at young adulthood exhibits a significant positive correlation with lifespan. The expression of these genes can be used as a predictor of longevity. Previous studies have reported genes that can act as predictors of lifespan ^42, 43, 44, 45, 46^. This is consistent with research showing that changes during development or early adulthood are sufficient to increase lifespan ^47, 48, 49, 50^ and suggest that an organism’s aging trajectory can be set early in life.

Comparing gene expression across pairs of long-lived mutants demonstrated that most long-lived mutants share a significant number of differentially expressed genes with multiple other long-lived mutants. The number of overlapping genes that we observed may be an underestimate of the actual number of overlapping genes. When we previously examined gene expression by RNA-seq in the same strain with the same experimental paradigm but at different times, we observed that there were significant differences in gene expression ^51^. Thus, at least some of the differences between the long-lived mutants may be due to normal variations in gene expression measurement. Nonetheless, the fact that there are large numbers of genes that are differentially expressed in the same direction in multiple long-lived mutants suggests that at least some of the genetic pathways contributing to lifespan extension in these mutants are shared.

At the same time, the long-lived mutants also exhibited significant differences in gene expression. An important finding of this work is that the long-lived mutants examined cluster into three distinct longevity groups based on gene expression. This clustering suggests that extended lifespan can arise from different molecular strategies. Moreover, the observation that many genes that are upregulated in group 1 longevity mutants are downregulated in group 2 longevity mutants, and vice versa, indicates that the same pathways can be modulated in opposite directions to increase lifespan. The group 1 longevity mutants (*daf-2, glp-1, clk-1, isp-1, nuo-6, sod-2*) exhibit activation of DAF-16 and ATFS-1 target genes and the loss of these transcription factors has been shown to decrease their longevity ^1, 16, 35^. The group 2 longevity mutants (*eat-2, osm-5*) exhibit upregulation of other genes, not upregulated in group 1 mutants, including PQM-1 target genes. *ife-2* (group 3) mutants exhibit the greatest number of transcriptional changes, which exhibit overlap with changes in both group 1 and group 2 mutants.

It is interesting to note that loss of *daf-16* decreases the lifespan of long-lived mutants in all three longevity groups (see our review ^36^). However, loss of *daf-16* also decreases wild-type lifespan. Thus, without further evidence it is hard to distinguish between the loss of *daf-16* non-specifically decreasing lifespan verses activation of DAF-16 actually contributing to lifespan extension. In *daf-2* mutants and the long-lived mitochondrial mutants (*clk-1, isp-1, nuo-6*), which are all part of longevity group 1, there is increased nuclear localization of DAF-16 and upregulation of DAF-16 target genes (e.g. see **Figure 5**). In addition, the differentially expressed genes in the long-lived mitochondrial mutants exhibit about a 50% overlap with the differentially expressed genes in *daf-2* mutants ^16^. In contrast, the group 2 longevity mutants *eat-2* and *osm-5* do not show consistent upregulation of DAF-16-activated genes and there is much less overlap between differentially expressed genes in *daf-2* mutants and differentially expressed genes in *eat-2* and *osm-5* mutants. We believe that these results are consistent with loss of DAF-16 causing a general decrease in lifespan and not specifically contributing to longevity in the group 2 longevity mutants *eat-2* and *osm-5*.

While our data identify several genes that are regulated in opposite directions in group 1 and group 2 longevity mutants, we do not yet know the extent to which each of these genes contribute to the longevity of group 1 and group 2 mutants. The three genes that we focused on for further characterization (C08F11.7, *ugt-62* and K05C4.9) are upregulated in group 1 longevity mutants but not group 2 mutants. Contrary to what might be expected, RNAi knockdown of these genes does not decrease the lifespan of the group 1 longevity mutant *daf-2* but does decrease the lifespan of the group 2 longevity mutant *eat-2*. We recently reviewed the effect of different resilience pathways on the lifespan of long-lived genetic mutants. Disruption of *daf-16, sek-1, skn-1, hsf-1, ire-1* and *trx-1* can decrease lifespan in both group 1 and group 2 longevity mutants, but also decreases lifespan in wild-type worms suggesting that at least in some mutants the effect on longevity may be non-specific ^36^. Disruption of *hif-1* does not affect the longevity of group 2 mutants, but does affect the lifespan of some group 1 mutants (*clk-1, isp-1, nuo-6*) but not others (*daf-2, glp-1*).

It is also unclear whether the longevity strategies utilized by group 1 and group 2 longevity mutants can operate simultaneously or whether they are mutually exclusive. To more definitively answer this question, it would be interesting to cross different combinations of group 1 and group 2 longevity mutants to see the extent to which different longevity groups synergize or potentially inhibit lifespan extension.

Among the genes commonly upregulated in multiple long-lived mutants, there was an enrichment for genes involved in innate immunity, defense and metabolism. Our group and others have previously demonstrated an important role for innate immune signaling in the extended longevity of long-lived mutants ^25, 26^. We and others have also shown that various pathways of cellular resilience promote longevity in long-lived mutants ^16, 17, 27, 36, 48, 52, 53^. Genes that were found to be downregulated in multiple long-lived mutants were involved in translation and gene expression, anabolic processes that promote growth. Previous work has demonstrated that decreasing translation is sufficient to extend lifespan ^7, 8, 40^. Together, these patterns support a model in which long-lived animals alter their metabolism to shift the balance from growth and biosynthesis toward defense and repair.

To move beyond correlation and test causation, we performed a targeted RNAi screen to examine the effect of knocking down genes that are upregulated in multiple long-lived mutants on lifespan. The majority of RNAi clones targeting commonly upregulated genes had no significant effect on longevity. This could be due to multiple reasons: (1) increased expression of the gene in long-lived mutants does not contribute to longevity; (2) increased expression of the gene contributes to longevity but the effect size is too small to detect when acting individually; (3) the level of knockdown achieved by the RNAi clone was not sufficient to inhibit gene function thereby resulting in no effect on longevity (we did not measure the level of knockdown achieved as we do not know what level of knockdown is need to affect function and it was unfeasible for this screen); or (4) the RNAi clone did not target the correct gene (the Ahringer RNAi library is known to have errors and we only confirmed the sequence of RNAi clones for positive hits).

Among the RNAi clones that significantly affected the longevity of *daf-2* or *nuo-6* worms, some clones increased lifespan and some decreased lifespan. We focused on the RNAi clones that decreased lifespan as this suggests that these genes are required for the longevity of these long-lived mutants. We selected seven genes to re-confirm in *nuo-6* mutants and test in wild-type worms. We found that RNAi clones targeting each of the seven genes (C08F11.7, *ugt-62*, K05C4.9, DC2.5, C05B5.5, T07C4.5 and W03B1.7) significantly decreased both *nuo-6* and wild-type lifespan. The fact that these genes also shorten wild-type lifespan suggests that these genes are acting as general pro-longevity factors and not specifically contributing to the extended lifespan of *nuo-6* worms. Based on this conclusion, we examined the effect of overexpressing three of these genes in a wild-type background. We found that overexpression of C08F11.7 results in increased lifespan, thermotolerance, and resistance to oxidative stress and bacterial pathogens. In contrast, overexpression of K05C4.9 significantly decreased lifespan. This indicates that not all genes that are upregulated in long-lived mutants are acting to promote longevity. In the case of K05C4.9, increasing or decreasing its levels reduces longevity suggesting that its levels need to be tightly regulated to achieve a normal lifespan. In future work it would be interesting to examine additional healthspan parameters in these strains ^54^.

Having quantified gene expression levels in nine different long-lived mutants, we developed a tool for the research community for examining expression of genes of interest across the panel of long-lived mutants in this study ^55^. This tool can be accessed at the website: https://vanraamsdonk.shinyapps.io/mutant_comparison_viewer/. This tool can be used to look up the expression of a gene or genes of interest in one or more of the long-lived mutants that we studied. The tool is online for everyone to use and can generate graphs comparing gene expression across long-lived mutants, indicate whether changes observed are statistically significant and provide access to the raw data. This tool can be used by researchers studying aging to test hypotheses about specific genes or genetic pathways that may be contributing to longevity. In addition to the online tool we generated for viewing our RNA-seq data, there are a number of excellent online tools that can be used to further analyse the data we generated in order to gain further insights into the aging process ^56^.

### Conclusions

In this work, we compared gene expression across a panel of long-lived genetic mutants. We found that these mutants share many gene expression changes suggesting that they employ a number of common genetic pathways to promote longevity. This includes pathways involved in cellular resilience and innate immunity. At the same time, the long-lived mutants also show unique patterns of gene expression that separates the mutants into three distinct longevity groups that in some cases modulate the same genetic pathways in opposite directions to achieve long life. While most individual differentially expressed genes do not exhibit a detectable effect on longevity, at least some of the differentially expressed genes can affect both lifespan and stress resistance individually. Overall, this work demonstrates that there are both shared and unique pathways contributing to lifespan extension in genetic long-lived mutants.

## Materials and Methods

### Strains

The strains used in this study included: N2 (wild-type), *ife-2 (ok306)*, *clk-1(qm30)*, *sod-2(ok1030)*, *eat-2(ad1116)*, *osm-5(p813)*, *nuo-6(qm200)*, *isp-1(qm150)*, *daf-2(e1370)*, *glp-1(e2141)*, PHX6925 sybIs6925[*eft-3p::K05C4.9, ges-1p::mCherry*], PHX6945 sybIs6945[*rpl-2p::ugt-62, myo-3p::mCherry*] and PHX6868 sybIs6868[*rpl-28p::C08F11.7, myo-2p::mCherry*]. PHX6925, PHX6945 and PHX 6868 were generated by SunyBiotech. All strains were grown and maintained in nematode grown medium (NGM) plates at 20°C except *glp-1(e2141)*, which was maintained at 20°C, but developed at 25°C for experiments. Plates were seeded with OP50 *E. coli* as a food source.

### Lifespan

Lifespan analyses were performed on NGM plates seeded with OP50 bacteria. For most lifespan experiments no 5-fluoro-2’-deoxyuridine (FUdR) was used. In cases where FUdR was utilized, we used a low concentration of 25 µM, which we have previously shown not to affect wild-type lifespan and have minimal effects on other genotypes ^57^. When FUdR was used, worms were transferred to the FUdR plates as prefertile young adults. Only adult lifespan was measured so that differences in development time would not affect the lifespan. During the lifespan study, worms were checked every 2-3 days until death. Worms with internal hatching or externalization of internal organs were removed from the study.

### RNA isolation

RNA was isolated using TRIZOL as described previously ^58^. Briefly, pre-fertile young adult worms were washed three times in M9 buffer and frozen in TRIZOL. After freeze-thawing three times, RNA was separated using chloroform, precipitated with isopropanol, and washed with 75% ethanol before being dissolved in RNAse free water. Isolated RNA was then frozen until library preparation and sequencing.

### RNA sequencing and Bioinformatic Analysis

Library preparation and RNA sequencing was performed as previously described ^15^. RNA-seq data is available on NCBI GEO: GSE179825 https://www.ncbi.nlm.nih.gov/geo/query/acc.cgi?acc=GSE179825 ^59, 60^, GSE93724 https://www.ncbi.nlm.nih.gov/geo/query/acc.cgi?acc=GSE93724 ^16^, GSE110984 https://www.ncbi.nlm.nih.gov/geo/query/acc.cgi?acc=GSE110984 ^17^ and was analyzed by the Harvard School of Public Health Bioinformatics core for this paper as previously described ^26, 59^. ***Read mapping and expression level estimation***. All samples were analyzed with an RNA-seq workflow provided by the bcbio-nextgen framework (https://bcbio-nextgen.readthedocs.org/en/latest/). Initial read quality was evaluated with FastQC (http://www.bioinformatics.babraham.ac.uk/projects/fastqc/) to verify that library preparation and sequencing produced data suitable for downstream processing. When required, adapter sequences, polyA contamination, and low-quality bases were removed using cutadapt (http://code.google.com/p/cutadapt/). Trimmed reads were aligned to the Ensembl build WBcel235 (release 90) of the *C.elegans* genome using STAR ^61^. Alignment quality was reviewed by examining metrics such as coverage uniformity, rRNA levels, genomic distribution of mapped reads (including matches to annotated transcripts and introns), and overall library complexity. Transcript abundances were quantified with Salmon ^62^ to identify transcript-level abundance estimates and then collapsed down to the gene-level using the R Bioconductor package tximport ^63^. Principal component analysis (PCA) and hierarchical clustering verified that samples grouped appropriately by batch and mutant background.

#### Differential gene expression and enrichment analysis

Gene-level differential expression was assessed with the DESeq2 package in R ^64^. For each comparison between mutant and wild-type samples, genes meeting a false discovery rate (FDR) cutoff of 0.01 were considered significant. When experiments spanned two sequencing batches, the model included a batch term to account for this source of variation.

#### Overlap analysis

Differentially expressed genes were divided according to whether they were up- or down-regulated and then compared across lists. The statistical significance of the overlaps was evaluated with a hypergeometric test.

#### Heatmap generation

For each dataset involving a long-lived mutant versus wild-type, raw counts were transformed using DESeq2’s regularized log (rlog) method ^64^, which stabilizes variance across the expression range and enhances unbiased clustering.

#### Weighted Venn diagrams

Weighted Venn diagrams were prepared using BioVenn https://www.biovenn.nl/.

#### UpSetR plots

UpsetR plots were prepared using R. https://pmc.ncbi.nlm.nih.gov/articles/PMC5870712/

#### Gene Ontology

Gene ontology enrichment plots were prepared using ShinyGO 0.85.1 https://bioinformatics.sdstate.edu/go/.

#### Overlap between gene sets

The significance of overlap between lists of differentially expressed genes was determined by calculating the Jaccard Index (number of overlapping genes/total number of genes) and significance was assessed using Fisher’s exact test.

### Targeted RNAi lifespan screen

For the targeted RNAi lifespan screen, there were RNAi clones for 154 of the 196 genes upregulated in 6 or more long-lived mutants. We found that several of these clones did not grow after two attempts leaving 116 clones. 52 of these clones were tested in *daf-2* mutants and 63 were tested in *nuo-6* mutants. For each lifespan trial, *daf-2* or *nuo-6* worms were synchronized by bleaching. The resulting eggs were placed on RNAi plates to start the lifespan study. Worms were transferred to new plates as necessary prior to offspring developing to adulthood. Survival of the worms was assessed approximately every 5 days.

### Heat stress assay

Prefertile young adult animals were transferred to a 35°C incubator on NGM plates seeded with OP50 bacteria. Survival was measured every hour beginning at 4 hours. Three biological replicates with 20 worms per replicate were performed.

### Bacterial pathogen stress assay

Resistance to bacterial pathogens was assessed by exposure to *P. aeruginosa* strain PA14 using a modified slow kill assay as we have done previously ^26, 60^. An overnight culture of *P. aeruginosa* strain PA14 was seeded onto NGM plates containing 20 mg/L FUdR. After growth at 37°C overnight and then room temperature overnight, day 3 adult worms, which had been grown on plates containing FUdR from adulthood, were transferred to the plates seeded with PA14. Survival was monitored daily with plates incubated at 20°C. Four biological replicates with 30-60 worms per replicate were performed.

### Osmotic stress assay

Worms at day 1 of adulthood were placed in 450 mM and 500 mM NaCl NGM plates seeded with 200 µL 5x OP50 bacteria^65^. Survival was scored after 48 hours of incubation at 20°C. Three biological replicates with 20 worms per replicate were performed.

### Paraquat assay

To assess chronic oxidative stress resistance, day 1 adult worms were placed in NGM plates with 4 mM Paraquat and 100 µL FUdR, prepared 3 days before the experiment^66^. The plates were seeded with 200 µl of 10X OP50 bacteria one day before the start of the experiment, protected from the light as they dried overnight. 20 worms were picked onto each plate and scoring was done every 2-3 days until death. Three biological replicates were performed.

### Anoxia assay

Day 1 adult animals were placed in NGM plates with 50 µM FUdR, seeded with 2x OP50. After picking the worms into the plates, the plates were placed in a BD GasPak™ EZ Anaerobe pouch system, which was tightly closed after the addition of the gas pack. Animals were placed on incubation at 20°C for 72 hours, and then the pouch was opened to allow reoxygenation.

Survival was scored after 24 hours of reoxygenation at 20°C. Three biological replicates with 20 worms each were performed.

### Statistical Analysis

Statistical analysis was performed using Graphpad PRISM Version 9.4.1. For lifespan and survival analyses, a log-rank test was used. For comparisons of a single variable across multiple groups, a one-way ANOVA with Dunnett’s multiple comparisons test was used. Error bars indicate standard error of the mean. *p<0.05, **p<0.01, ***p<0.001, ****p<0.0001.

## Acknowledgments

Some strains were provided by the CGC, which is funded by NIH Office of Research Infrastructure Programs (P30 OD010440). We would also like to acknowledge the *C. elegans* knockout consortium and the National Bioresource Project of Japan for providing strains used in this research.

## Competing interests

The authors have declared that no competing interests exist.

## Author Contributions

Conceptualization: JVR. Methodology: ZDR, JG, ATG, GB, SKS, UA, MM, MMS, JVR. Investigation: ZDR, JG, ATG, GB, SKS, UA, MM, MMS, JVR. Visualization: ZDR, JG, ATG, GB, SKS, MM, JVR. Writing – original draft: JVR. Writing – review and editing: ZDR, JG, ATG, GB, SKS, UA, MM, MMS, JVR. Supervision: JVR.

## Materials & Correspondence

Correspondence and material requests should be addressed to Jeremy Van Raamsdonk.

## Data availability

RNA-seq data has been deposited on GEO: GSE179825, GSE93724, GSE110984. All other data and strains generated in the current study are included with the manuscript or available from the corresponding author on request.

## Funding

This work was supported by the Canadian Institutes of Health Research (CIHR; http://www.cihr-irsc.gc.ca/; JVR) and the Natural Sciences and Engineering Research Council of Canada (NSERC; https://www.nserc-crsng.gc.ca/index_eng.asp; JVR). JVR was the recipient of a Research Scholar career award from the Fonds de Recherche du Québec Santé (FRQS) and Parkinson Quebec. SKS and ZDR were supported by training awards from the FRQS. The funders had no role in study design, data collection and analysis, decision to publish, or preparation of the manuscript.

## Supplemental Figures

**Figure S1.**
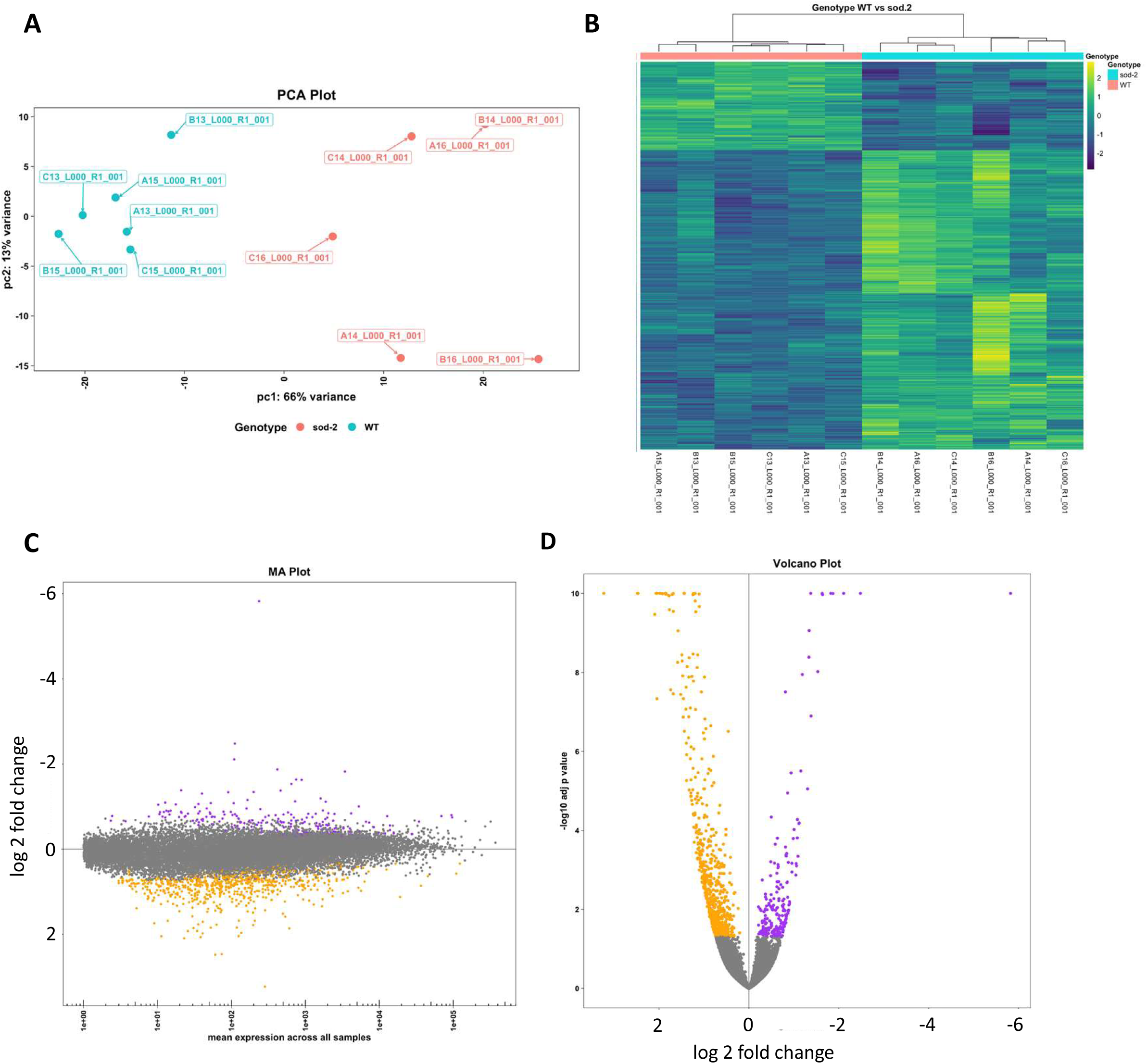
Transcriptomic analysis of long-lived *sod-2* mutants. (**A**) Principal component analysis plot. (**B**) Heatmap comparing gene expression to wild-type (WT) worms. (**C**) Mean average (MA) plot examining log fold change across all genes. (**D**) Volcano plot comparing adjusted p-value and log fold change across all genes.

**Figure S2.**
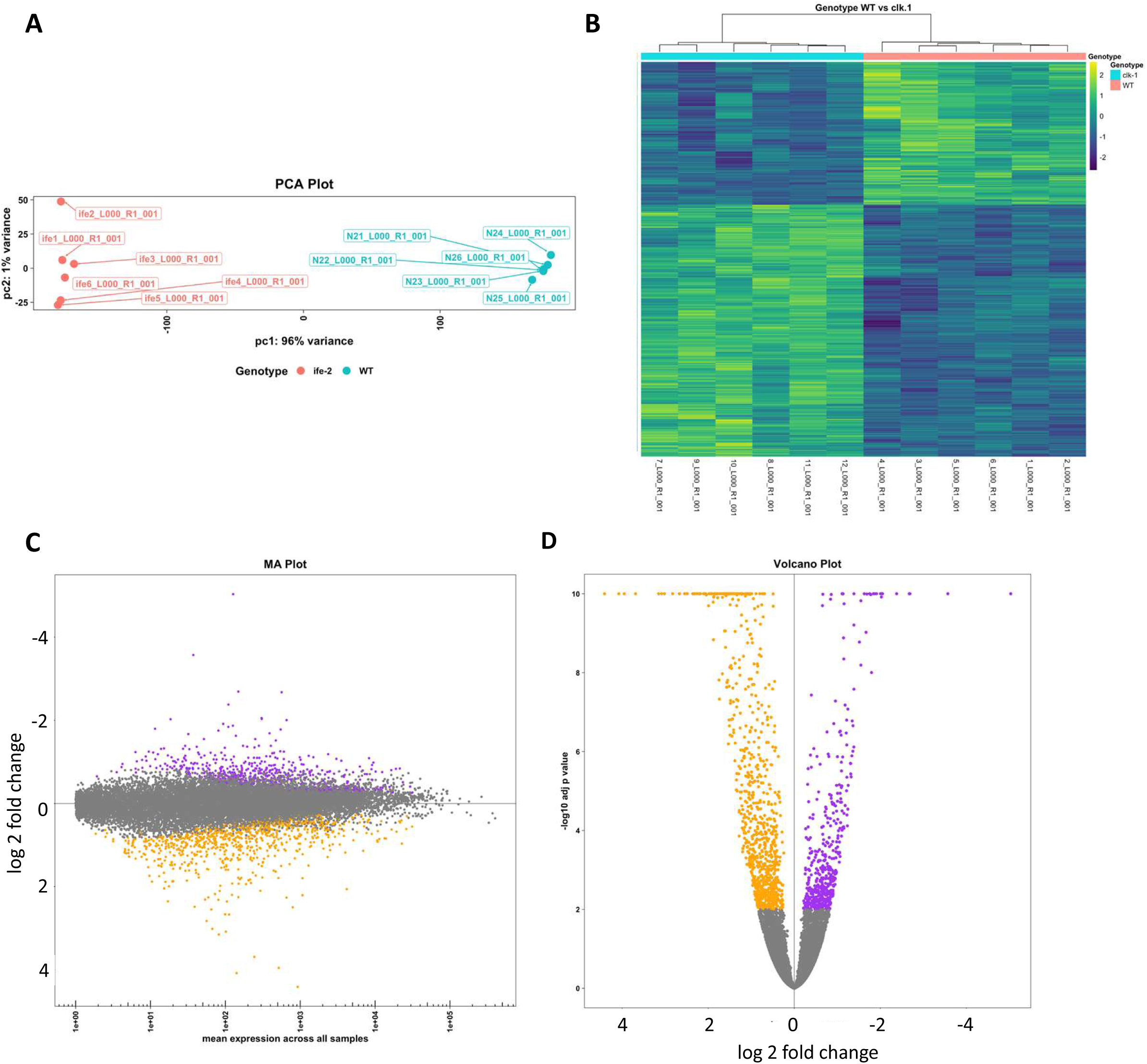
Transcriptomic analysis of long-lived *clk-1* mutants. (**A**) Principal component analysis plot. (**B**) Heatmap comparing gene expression to wild-type (WT) worms. (**C**) Mean average (MA) plot examining log fold change across all genes. (**D**) Volcano plot comparing adjusted p-value and log fold change across all genes.

**Figure S3.**
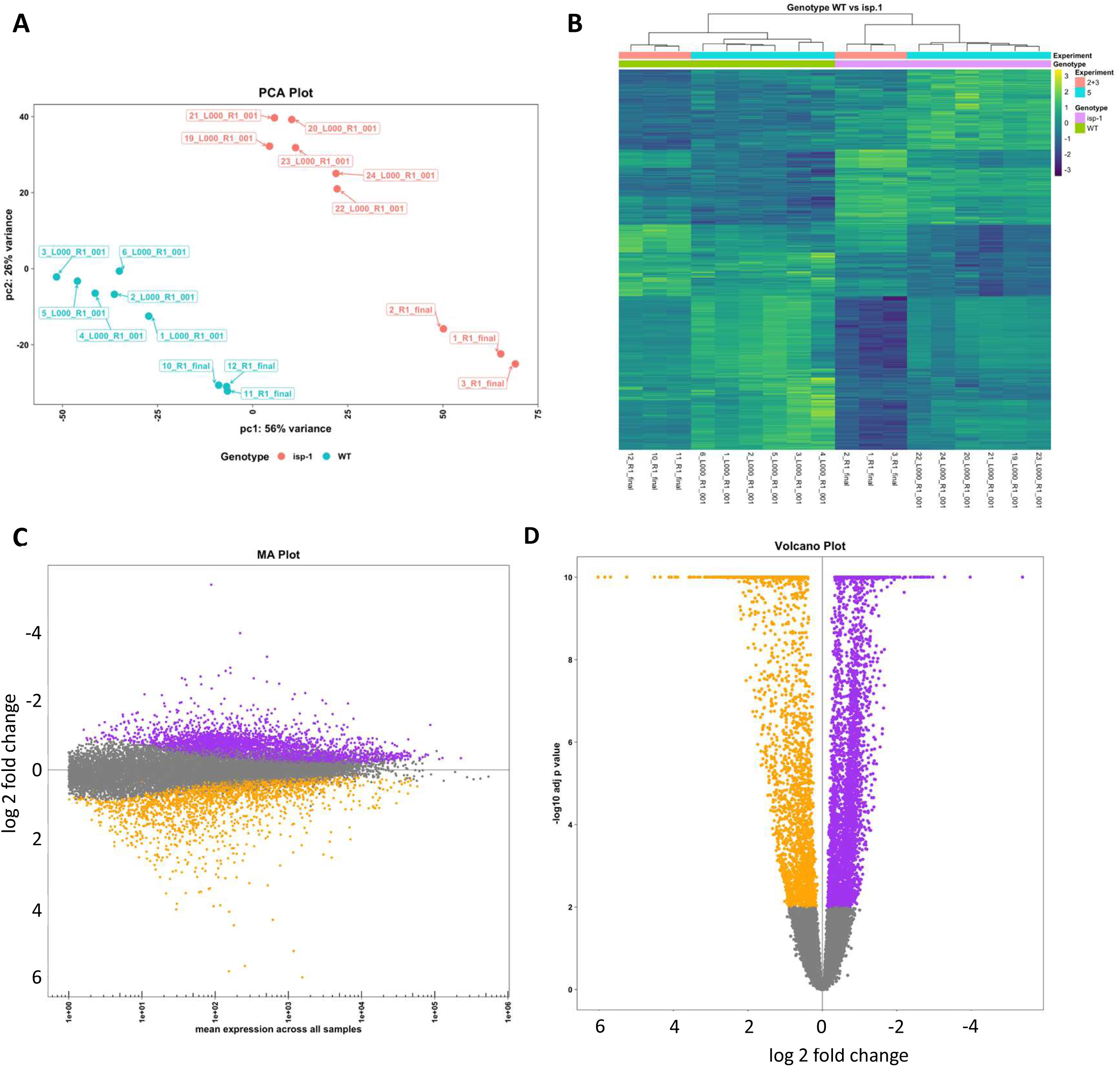
Transcriptomic analysis of long-lived *isp-1* mutants. (**A**) Principal component analysis plot. (**B**) Heatmap comparing gene expression to wild-type (WT) worms. (**C**) Mean average (MA) plot examining log fold change across all genes. (**D**) Volcano plot comparing adjusted p-value and log fold change across all genes.

**Figure S4.**
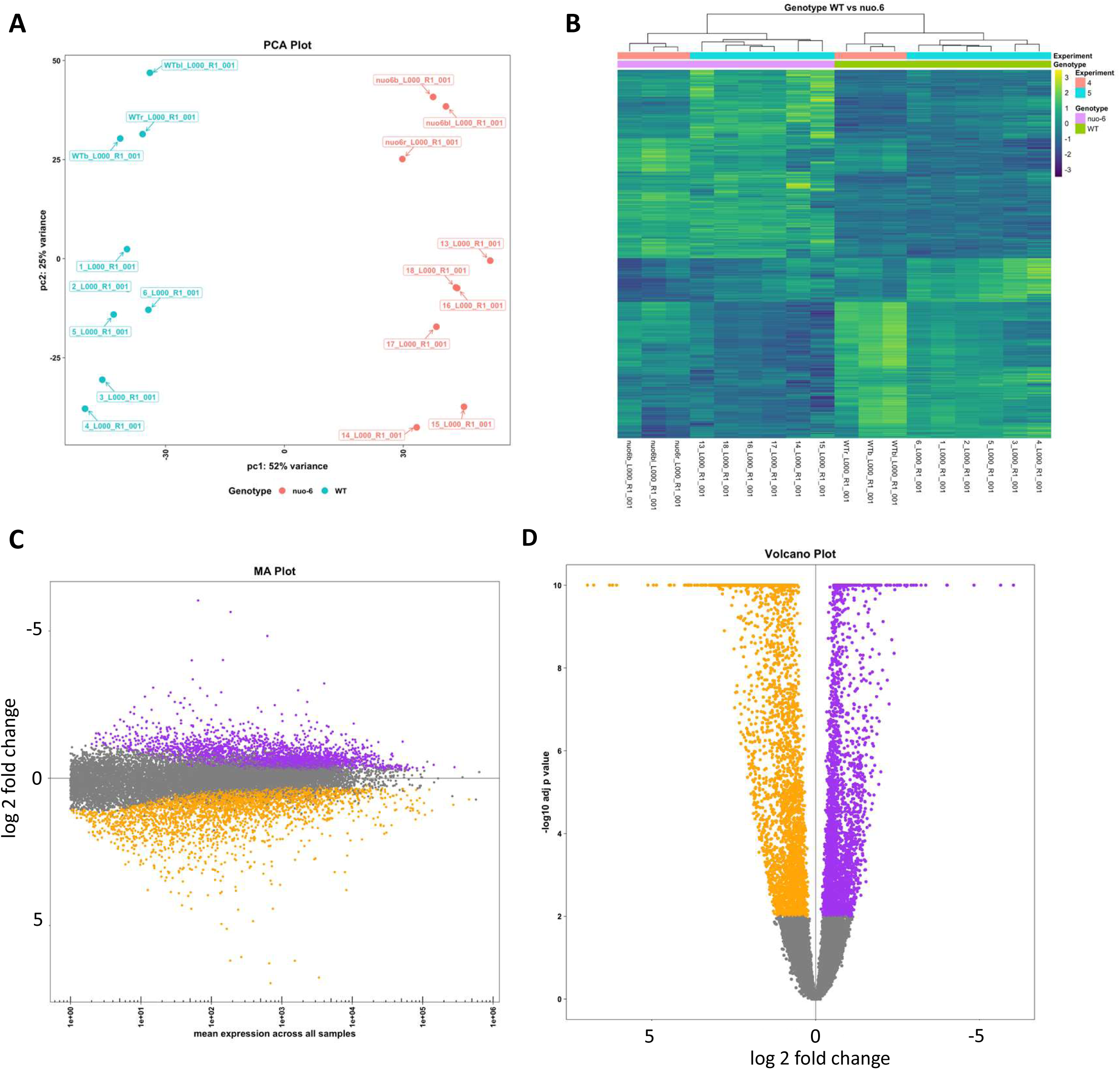
Transcriptomic analysis of long-lived *nuo-6* mutants. (**A**) Principal component analysis plot. (**B**) Heatmap comparing gene expression to wild-type (WT) worms. (**C**) Mean average (MA) plot examining log fold change across all genes. (**D**) Volcano plot comparing adjusted p-value and log fold change across all genes.

**Figure S5.**
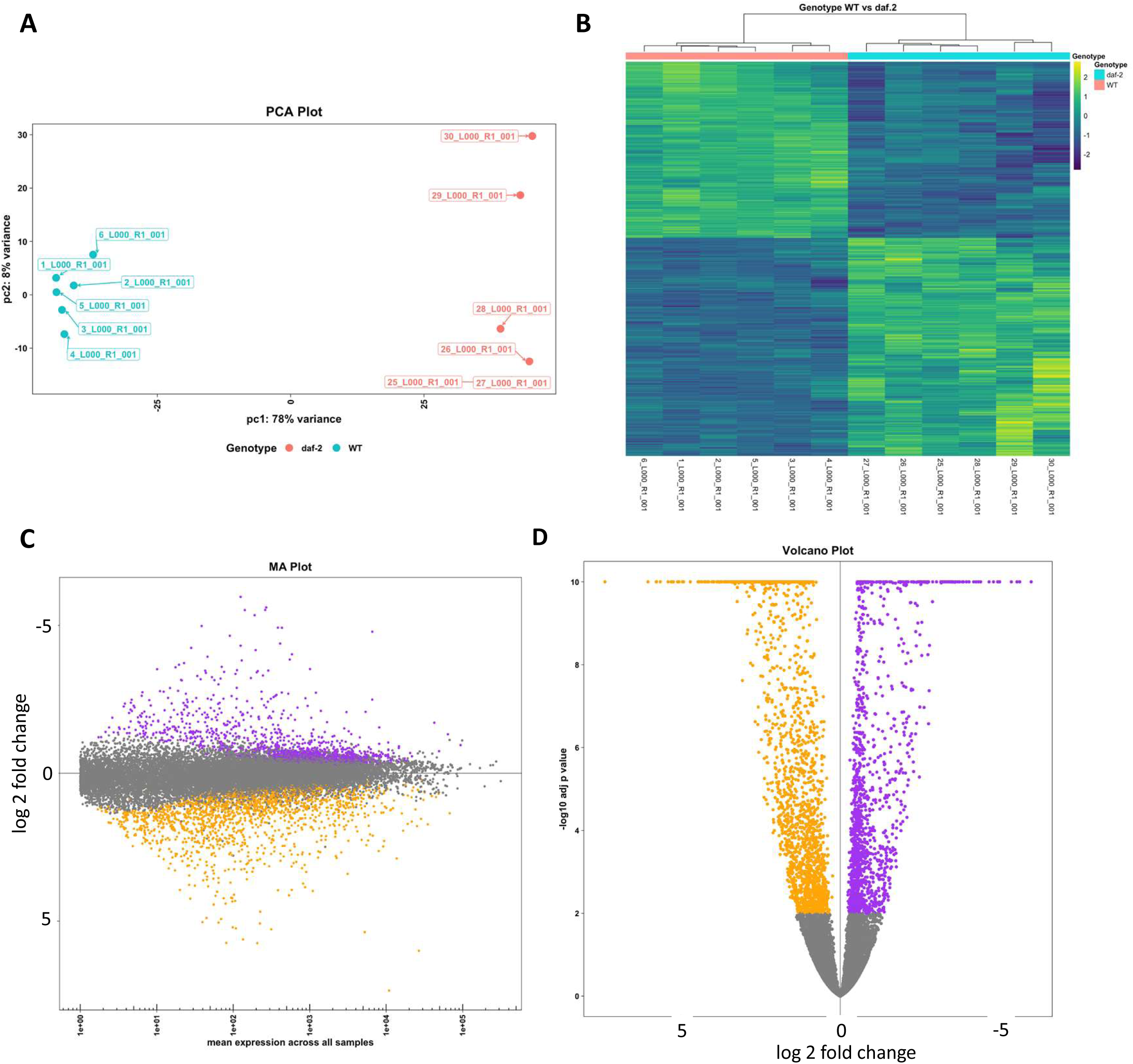
Transcriptomic analysis of long-lived *daf-2* mutants. (**A**) Principal component analysis plot. (**B**) Heatmap comparing gene expression to wild-type (WT) worms. (**C**) Mean average (MA) plot examining log fold change across all genes. (**D**) Volcano plot comparing adjusted p-value and log fold change across all genes.

**Figure S6.**
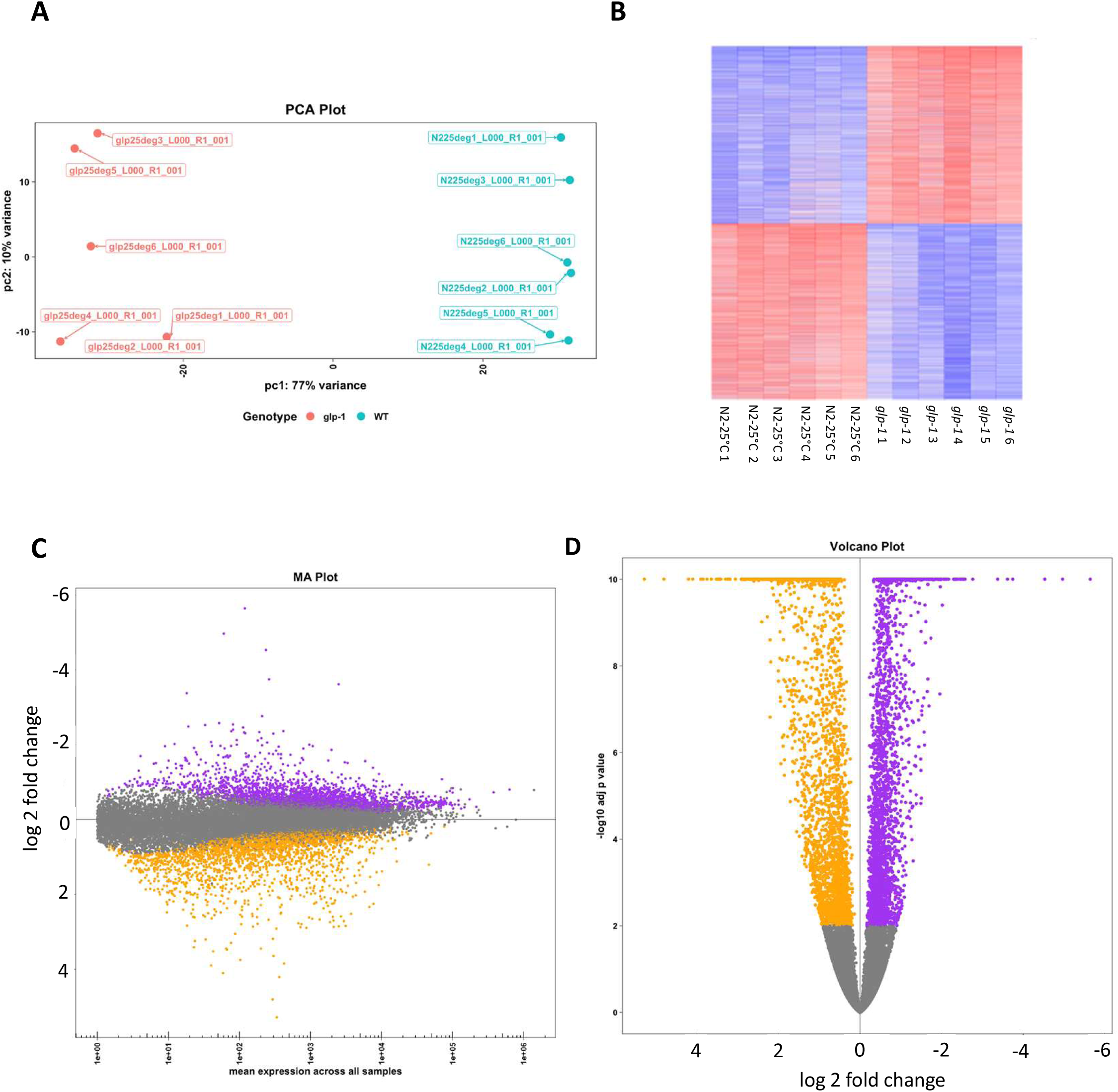
Transcriptomic analysis of long-lived *glp-1* mutants. (**A**) Principal component analysis plot. (**B**) Heatmap comparing gene expression to wild-type (WT) worms. (**C**) Mean average (MA) plot examining log fold change across all genes. (**D**) Volcano plot comparing adjusted p-value and log fold change across all genes.

**Figure S7.**
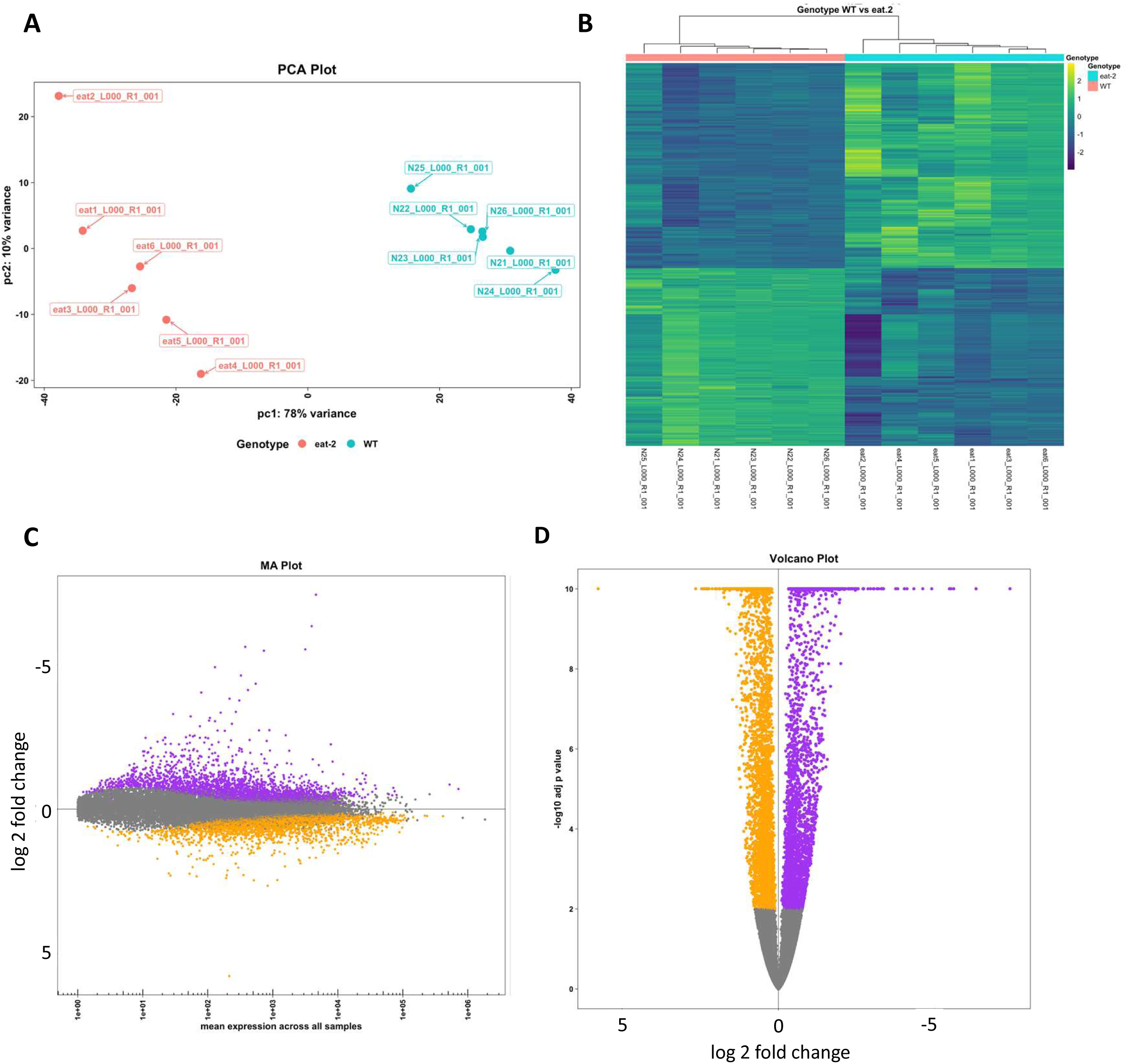
Transcriptomic analysis of long-lived *eat-2* mutants. (**A**) Principal component analysis plot. (**B**) Heatmap comparing gene expression to wild-type (WT) worms. (**C**) Mean average (MA) plot examining log fold change across all genes. (**D**) Volcano plot comparing adjusted p-value and log fold change across all genes.

**Figure S8.**
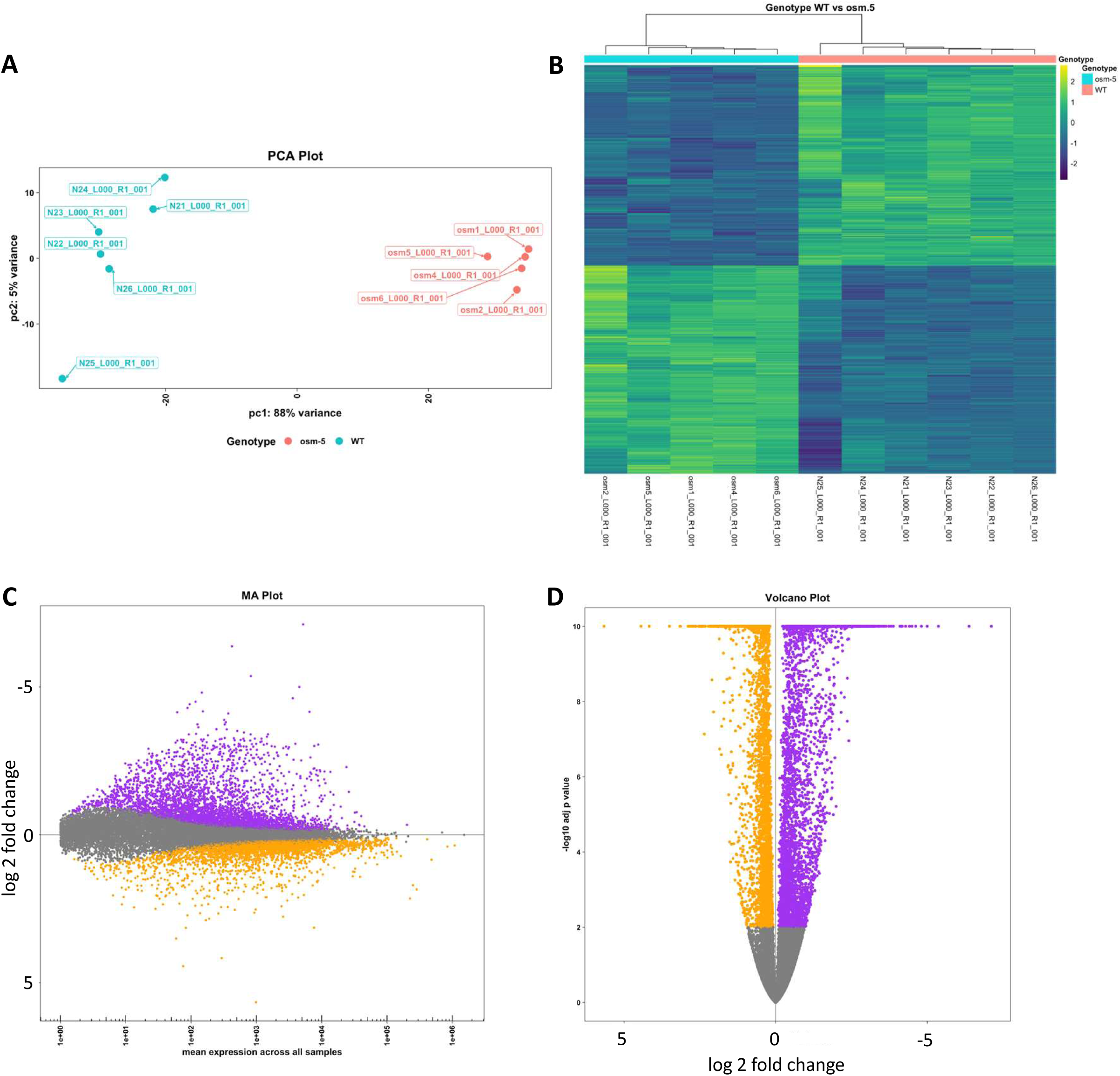
Transcriptomic analysis of long-lived *osm-5* mutants. (**A**) Principal component analysis plot. (**B**) Heatmap comparing gene expression to wild-type (WT) worms. (**C**) Mean average (MA) plot examining log fold change across all genes. (**D**) Volcano plot comparing adjusted p-value and log fold change across all genes.

**Figure S9.**
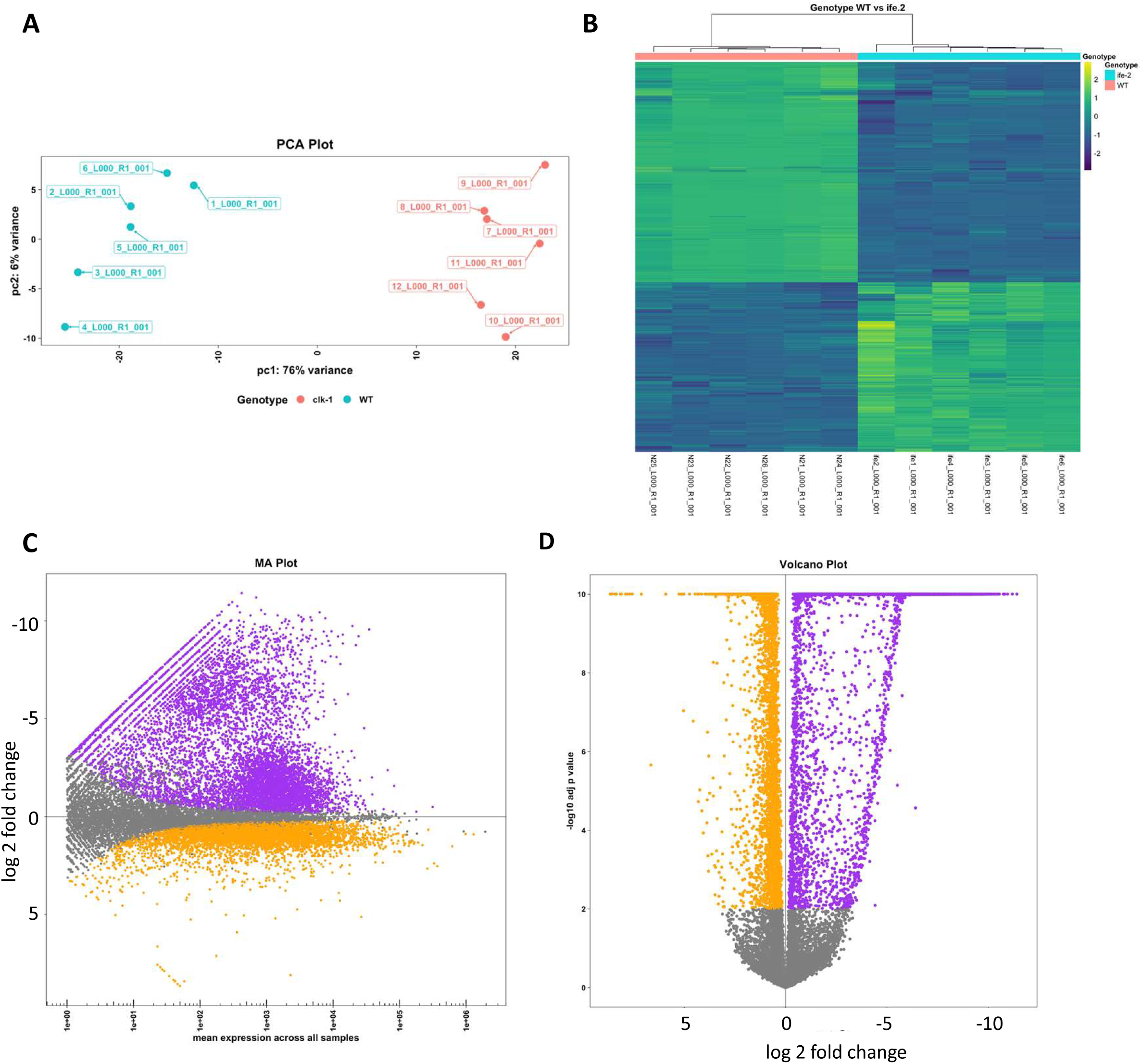
Transcriptomic analysis of long-lived *ife-2* mutants. (**A**) Principal component analysis plot. (**B**) Heatmap comparing gene expression to wild-type (WT) worms. (**C**) Mean average (MA) plot examining log fold change across all genes. (**D**) Volcano plot comparing adjusted p-value and log fold change across all genes.

**Figure S10.**
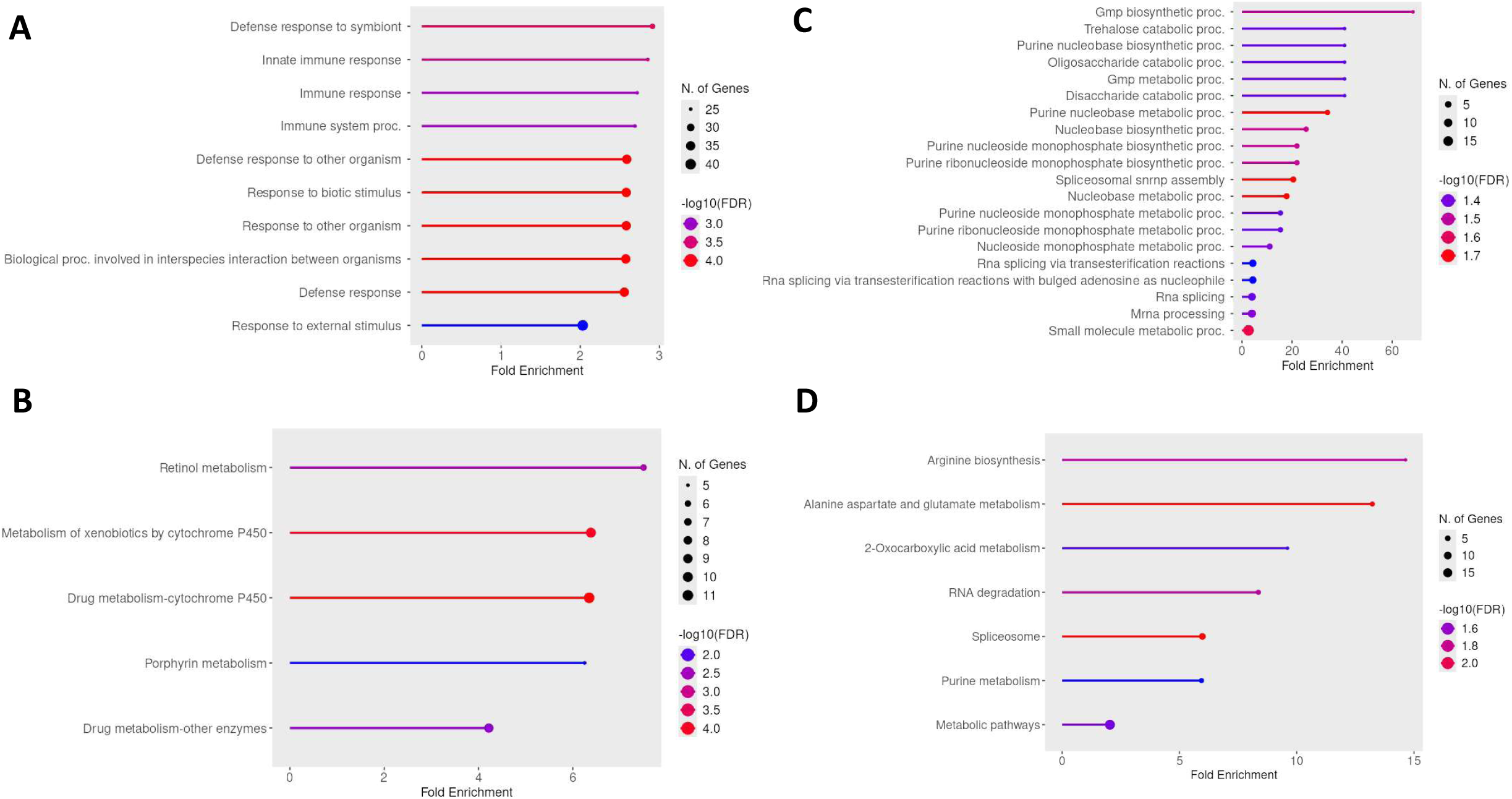
Enrichment analysis of differentially expressed genes in long-lived *sod-2* mutants. (**A**) Gene ontology (GO) term enrichment of upregulated genes. (**B**) Kegg pathway enrichment for upregulated genes. (**C**) GO term enrichment for downregulated genes. (**D**) Kegg pathway enrichment for downregulated genes. Enrichment analysis was performed using ShingGo 0.85.1 https://bioinformatics.sdstate.edu/go/.

**Figure S11.**
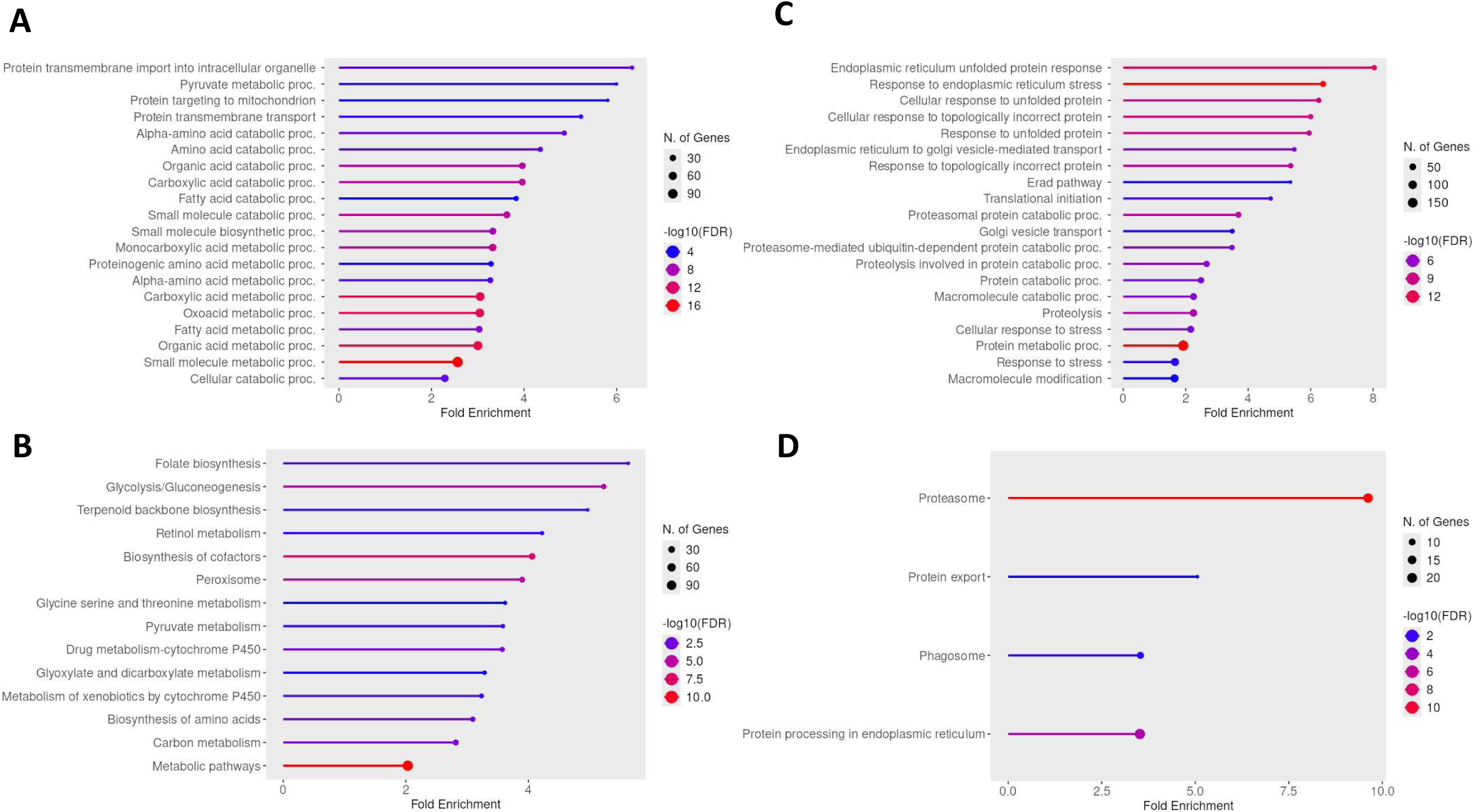
Enrichment analysis of differentially expressed genes in long-lived *clk-1* mutants. (**A**) Gene ontology (GO) term enrichment of upregulated genes. (**B**) Kegg pathway enrichment for upregulated genes. (**C**) GO term enrichment for downregulated genes. (**D**) Kegg pathway enrichment for downregulated genes. Enrichment analysis was performed using ShingGo 0.85.1 https://bioinformatics.sdstate.edu/go/.

**Figure S12.**
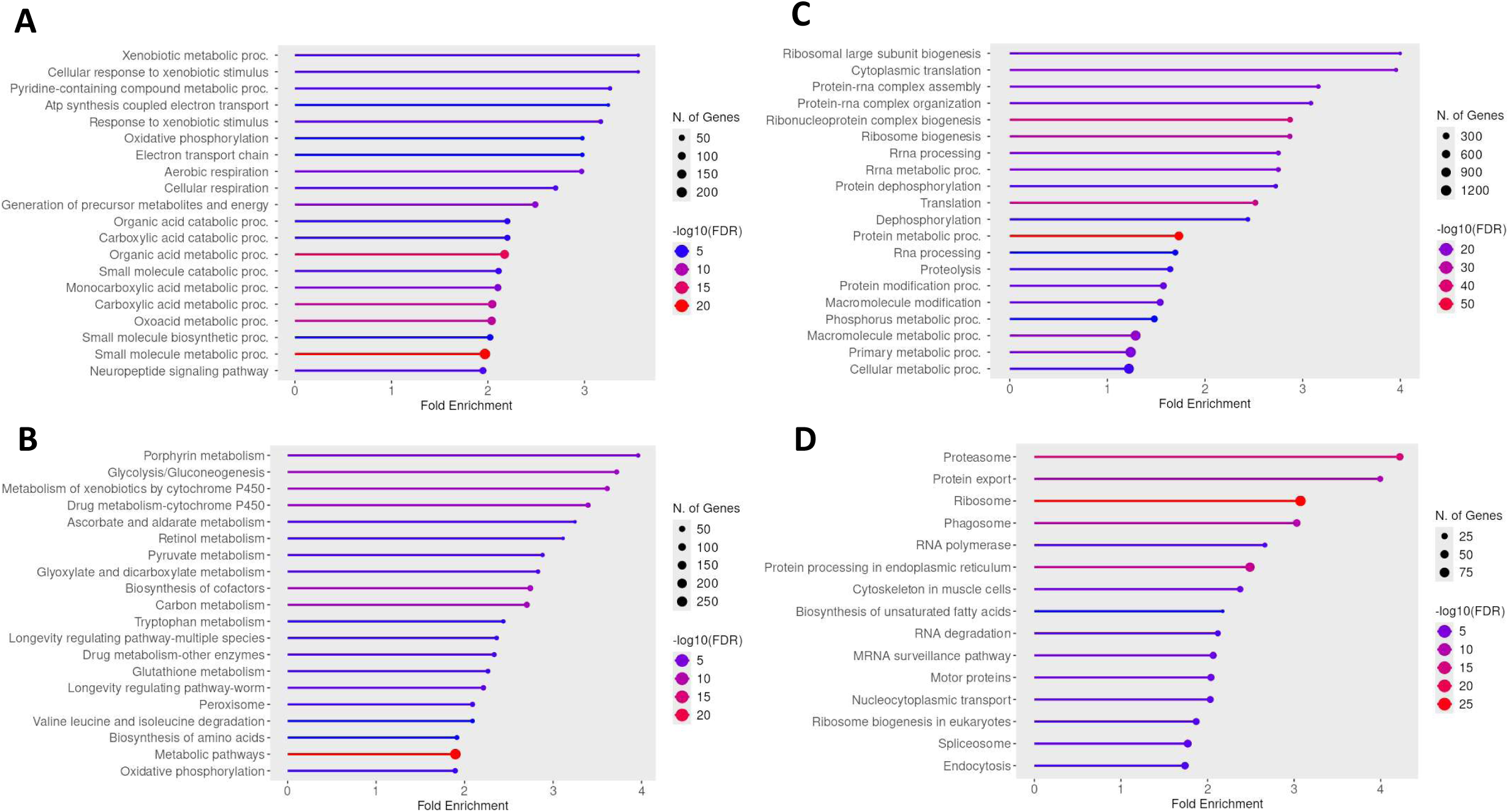
Enrichment analysis of differentially expressed genes in long-lived *isp-1* mutants. (**A**) Gene ontology (GO) term enrichment of upregulated genes. (**B**) Kegg pathway enrichment for upregulated genes. (**C**) GO term enrichment for downregulated genes. (**D**) Kegg pathway enrichment for downregulated genes. Enrichment analysis was performed using ShingGo 0.85.1 https://bioinformatics.sdstate.edu/go/.

**Figure S13.**
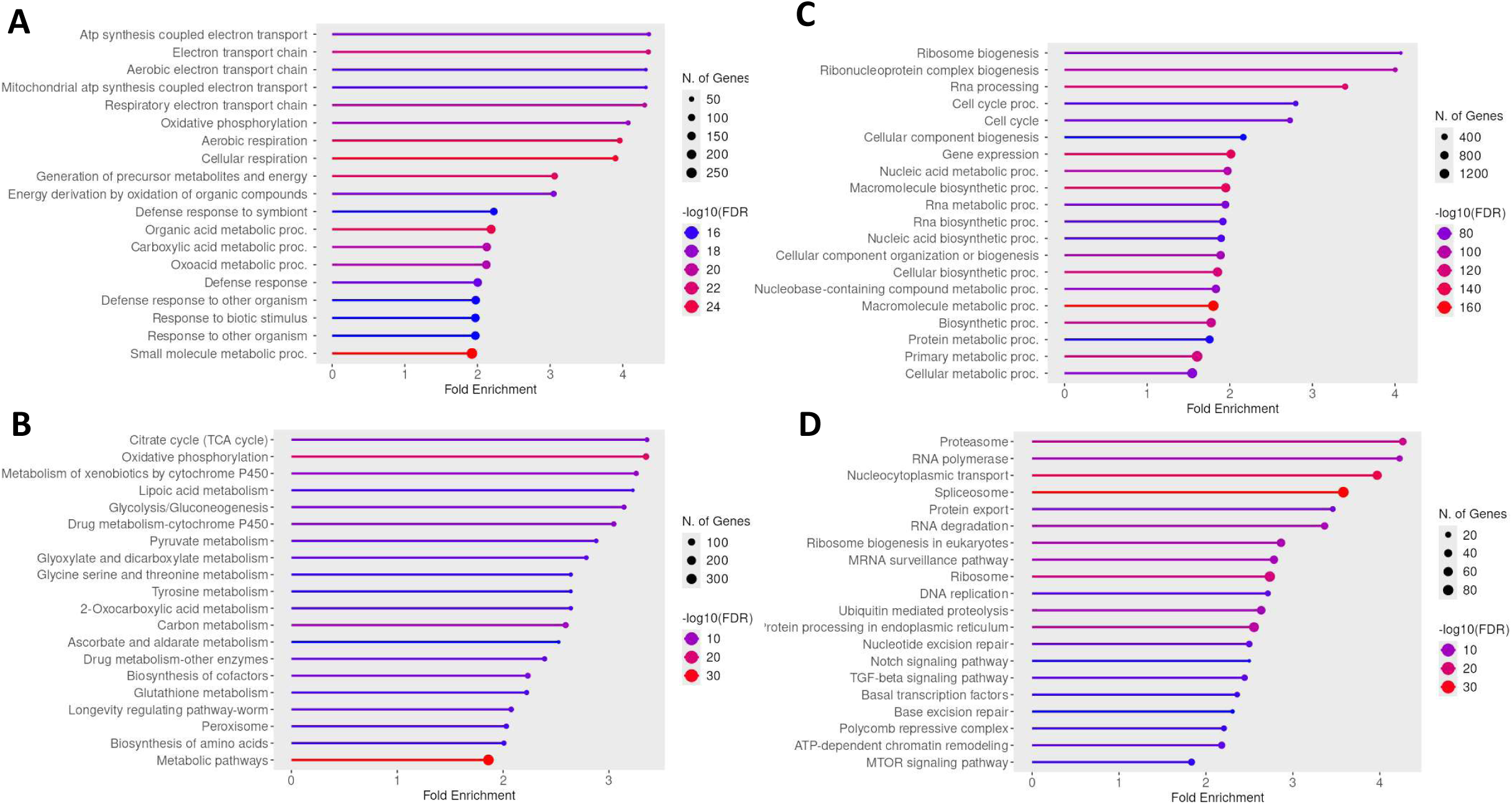
Enrichment analysis of differentially expressed genes in long-lived *nuo-6* mutants. (**A**) Gene ontology (GO) term enrichment of upregulated genes. (**B**) Kegg pathway enrichment for upregulated genes. (**C**) GO term enrichment for downregulated genes. (**D**) Kegg pathway enrichment for downregulated genes. Enrichment analysis was performed using ShingGo 0.85.1 https://bioinformatics.sdstate.edu/go/.

**Figure S14.**
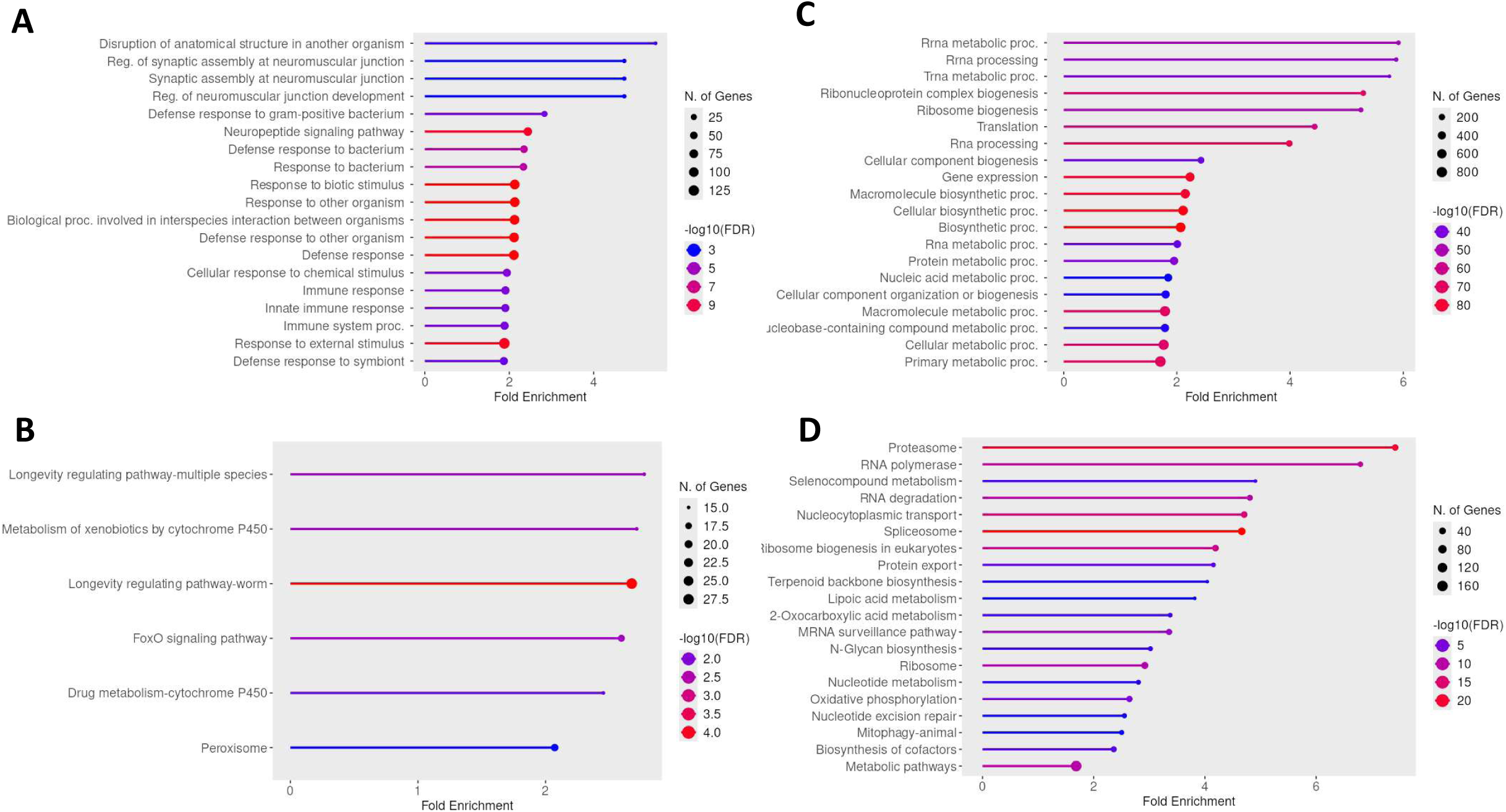
Enrichment analysis of differentially expressed genes in long-lived *daf-2* mutants. (**A**) Gene ontology (GO) term enrichment of upregulated genes. (**B**) Kegg pathway enrichment for upregulated genes. (**C**) GO term enrichment for downregulated genes. (**D**) Kegg pathway enrichment for downregulated genes. Enrichment analysis was performed using ShingGo 0.85.1 https://bioinformatics.sdstate.edu/go/.

**Figure S15.**
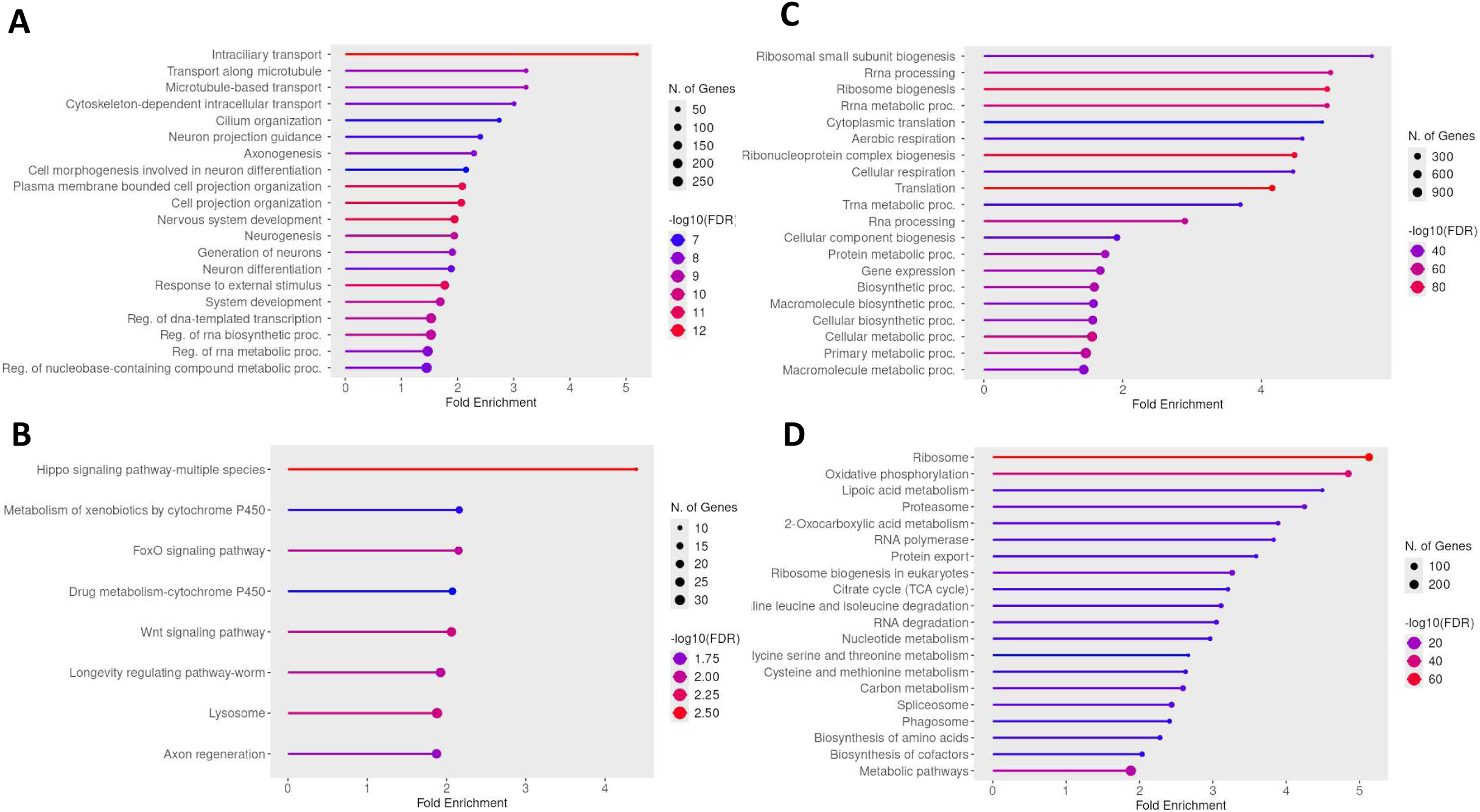
Enrichment analysis of differentially expressed genes in long-lived *glp-1* mutants. (**A**) Gene ontology (GO) term enrichment of upregulated genes. (**B**) Kegg pathway enrichment for upregulated genes. (**C**) GO term enrichment for downregulated genes. (**D**) Kegg pathway enrichment for downregulated genes. Enrichment analysis was performed using ShingGo 0.85.1 https://bioinformatics.sdstate.edu/go/.

**Figure S16.**
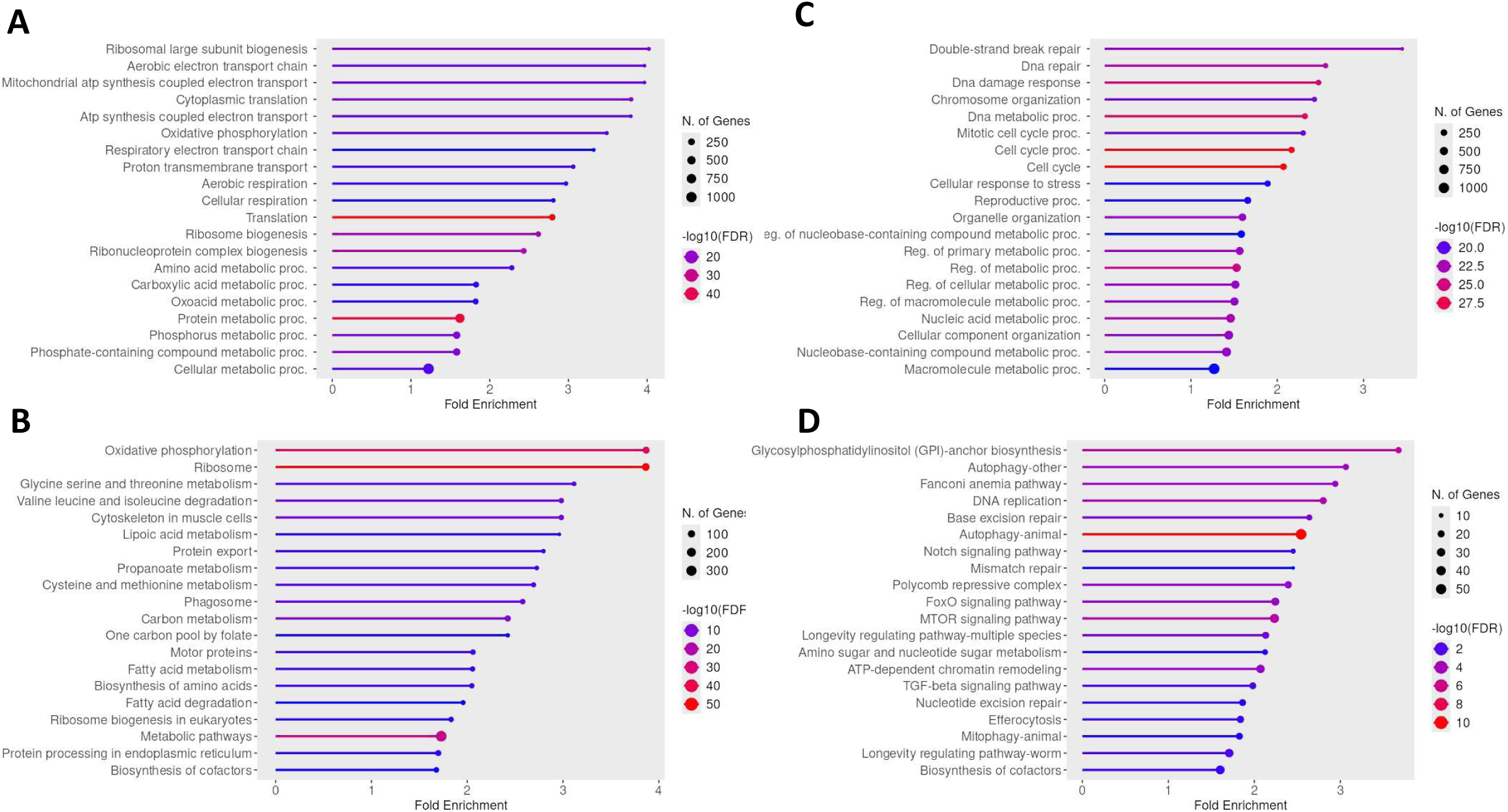
Enrichment analysis of differentially expressed genes in long-lived *eat-2* mutants. (**A**) Gene ontology (GO) term enrichment of upregulated genes. (**B**) Kegg pathway enrichment for upregulated genes. (**C**) GO term enrichment for downregulated genes. (**D**) Kegg pathway enrichment for downregulated genes. Enrichment analysis was performed using ShingGo 0.85.1 https://bioinformatics.sdstate.edu/go/.

**Figure S17.**
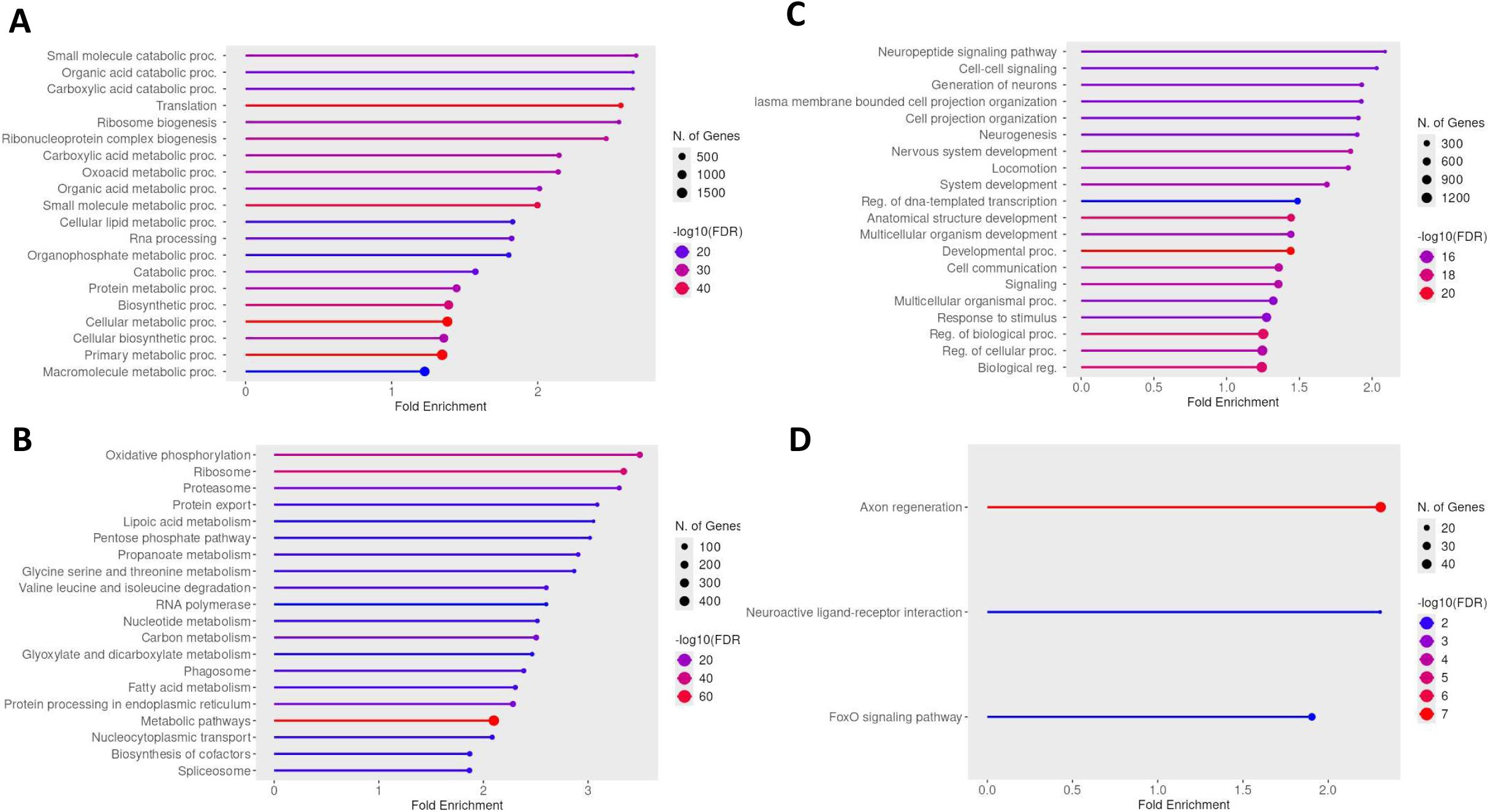
Enrichment analysis of differentially expressed genes in long-lived *osm-5* mutants. (**A**) Gene ontology (GO) term enrichment of upregulated genes. (**B**) Kegg pathway enrichment for upregulated genes. (**C**) GO term enrichment for downregulated genes. (**D**) Kegg pathway enrichment for downregulated genes. Enrichment analysis was performed using ShingGo 0.85.1 https://bioinformatics.sdstate.edu/go/.

**Figure S18.**
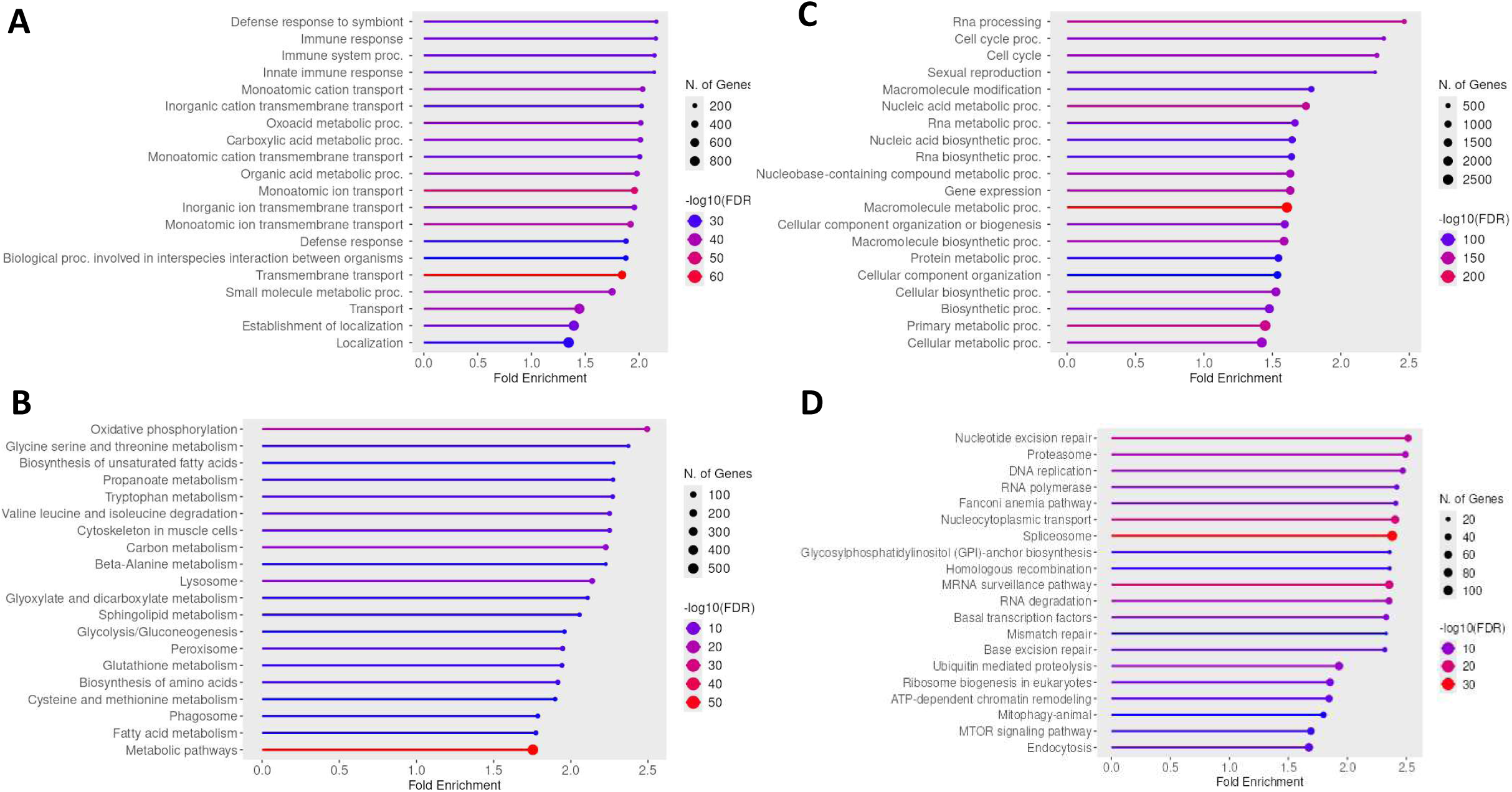
Enrichment analysis of differentially expressed genes in long-lived *ife-2* mutants. (**A**) Gene ontology (GO) term enrichment of upregulated genes. (**B**) Kegg pathway enrichment for upregulated genes. (**C**) GO term enrichment for downregulated genes. (**D**) Kegg pathway enrichment for downregulated genes. Enrichment analysis was performed using ShingGo 0.85.1 https://bioinformatics.sdstate.edu/go/.

**Figure S19.**
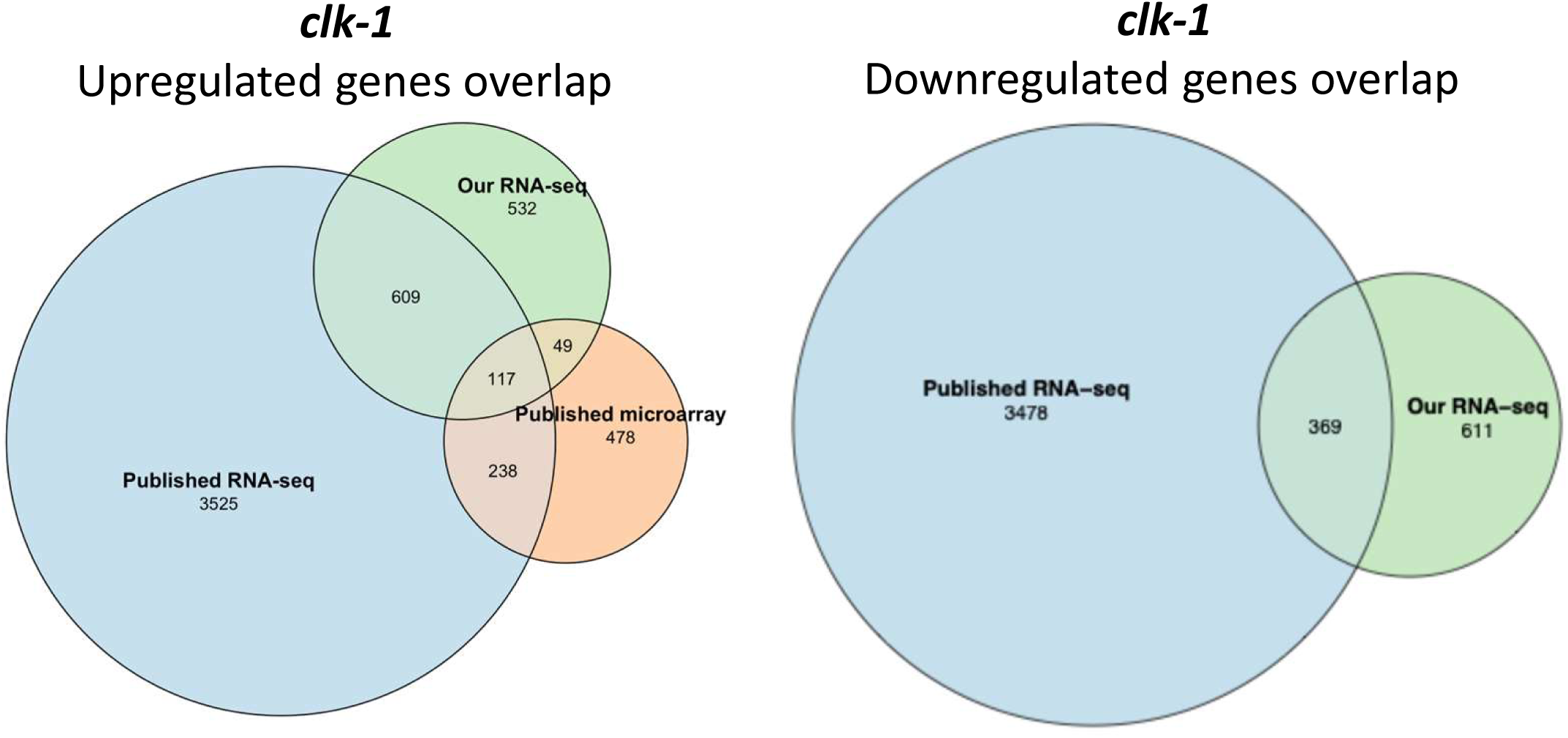
Comparison of gene expression changes in *clk-1* mutants in the current study to previous published gene expression results. The published microarray data is from Cristina et al. 2009 *PLoS Genetics* Microarray: https://doi.org/10.1371/journal.pgen.1000450. The published RNA-seq data is from Dutta et al. 2025 *PLoS Biology* (https://doi.org/10.1371/journal.pbio.3003504).

**Figure S20.**
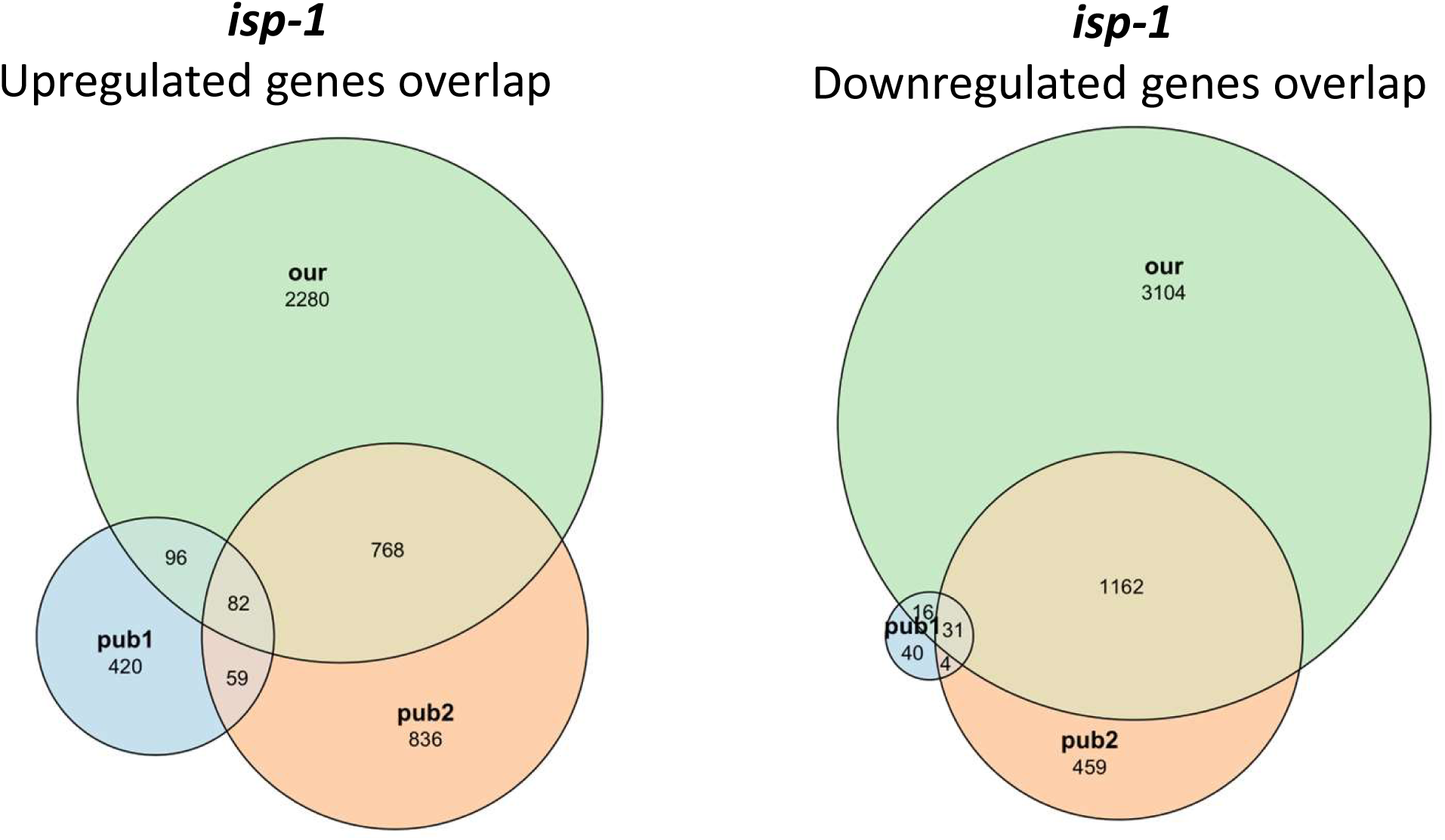
Comparison of gene expression changes in *isp-1* mutants in the current study to previous published gene expression results. The published microarray data is from *Cristina* et al. 2009 *PLoS Genetics (*https://doi.org/10.1371/journal.pgen.1000450) and Yee et al. 2014 *Cell* https://doi.org/10.1016/j.cell.2014.02.055

**Figure S21.**
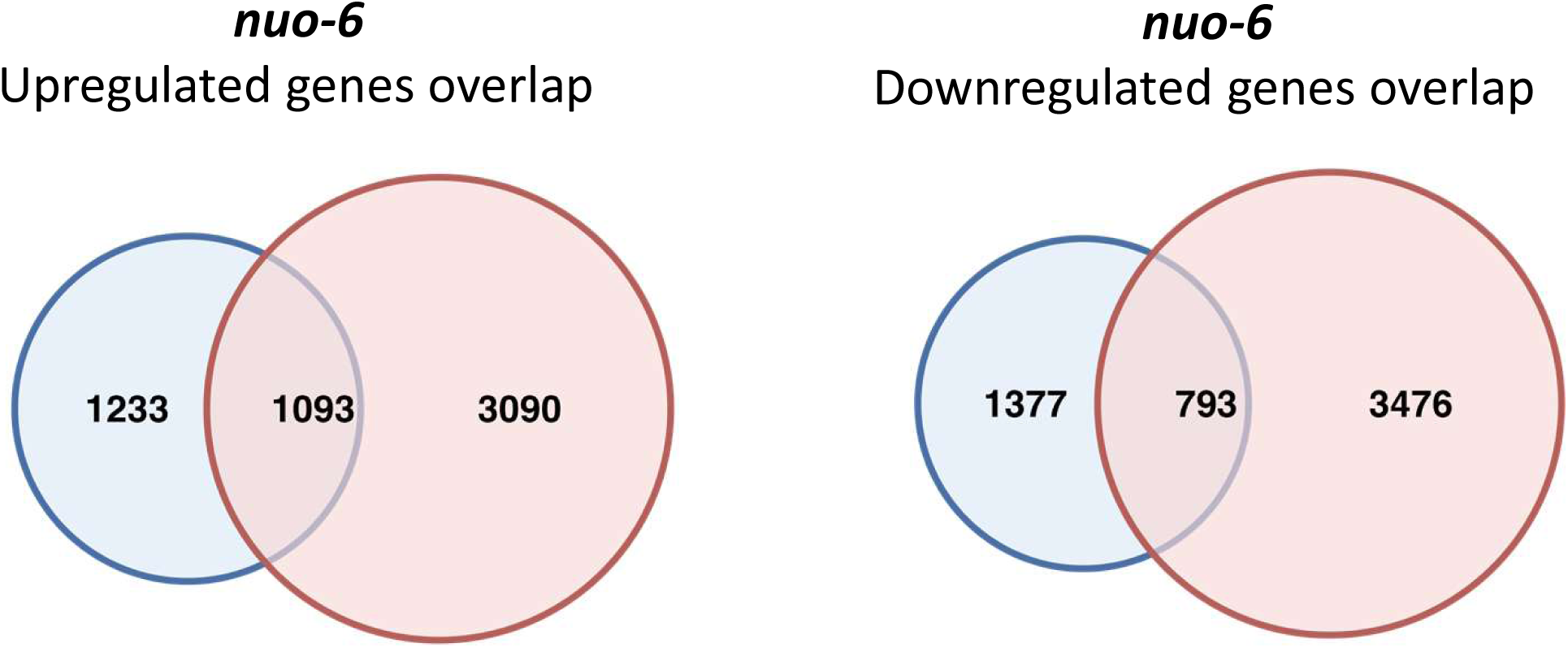
Comparison of gene expression changes in *nuo-6* mutants in the current study to previous published gene expression results. The published microarray data is from Yee et al. 2014 *Cell* (https://doi.org/10.1016/j.cell.2014.02.055).

**Figure S22.**
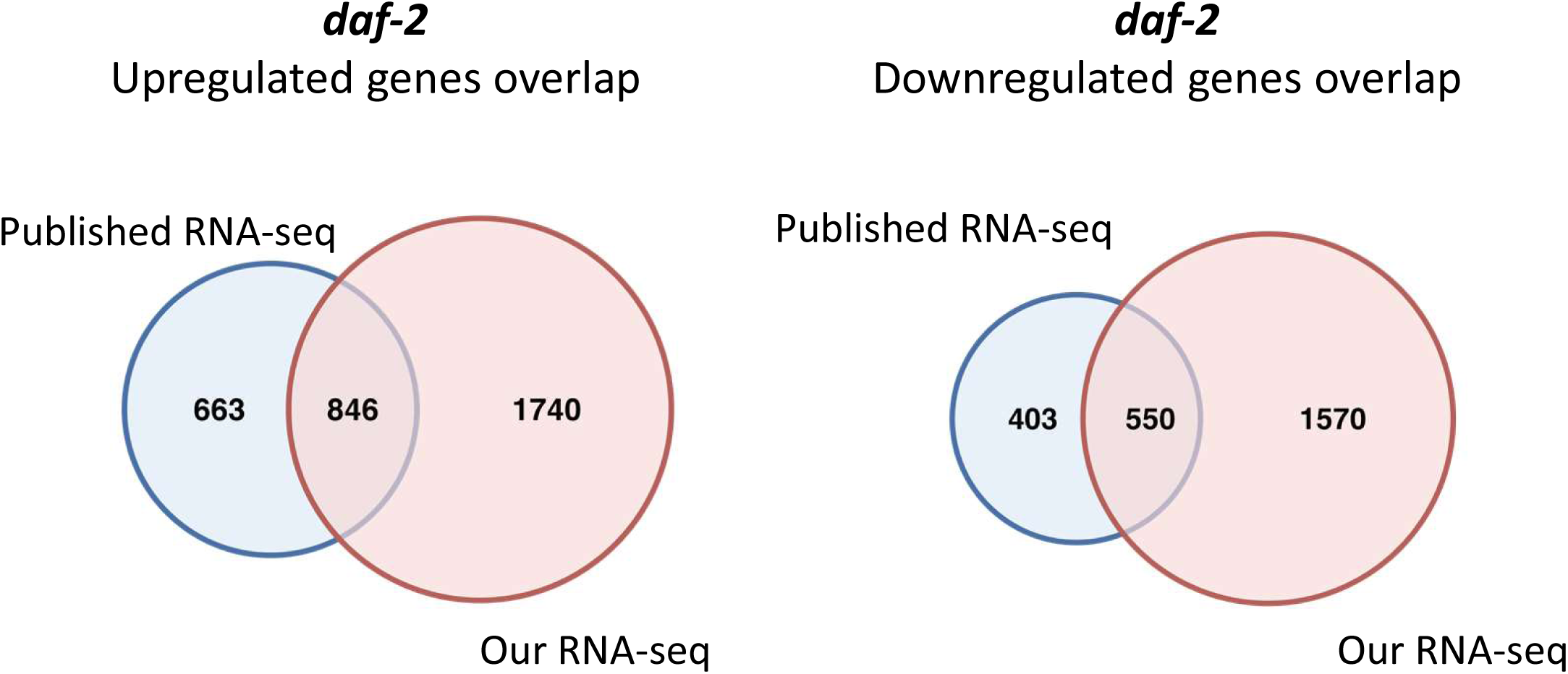
Comparison of gene expression changes in *daf-2* mutants in the current study to previous published gene expression results. The published RNA-seq data is from Zhang et al. 2022 *Nature Communications* (https://doi.org/10.1038/s41467-022-33850-4).

**Figure S23.**
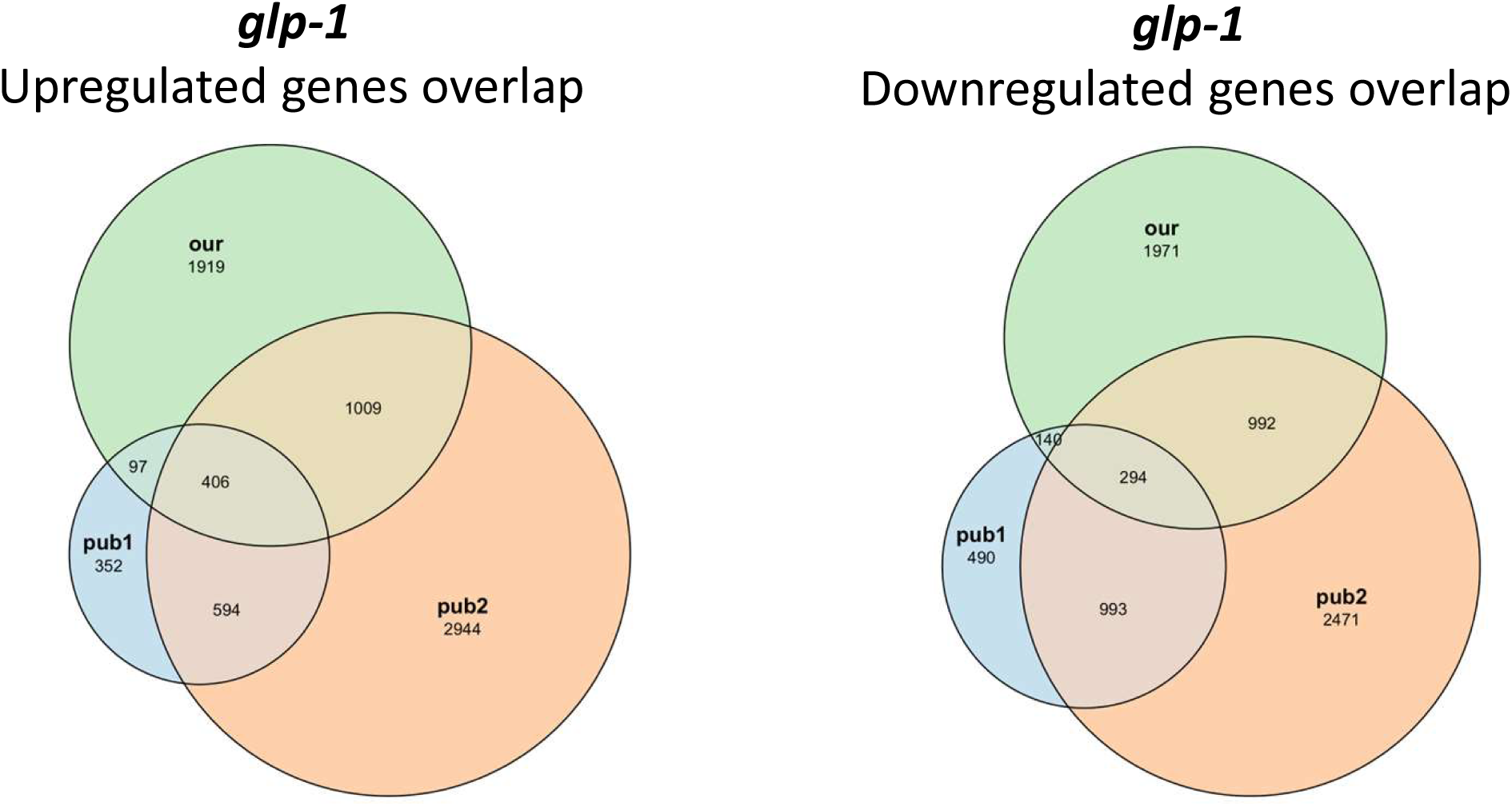
Comparison of gene expression changes in *glp-1* mutants in the current study to previous published gene expression results. The published RNA-seq data is from Chaturbedi et al., 2025 *Nature Communications* (https://doi.org/10.1038/s41467-025-64341-x) and Steinbaugh et al. 2015, *eLife* (https://doi.org/10.7554/eLife.07836).

**Figure S24.**
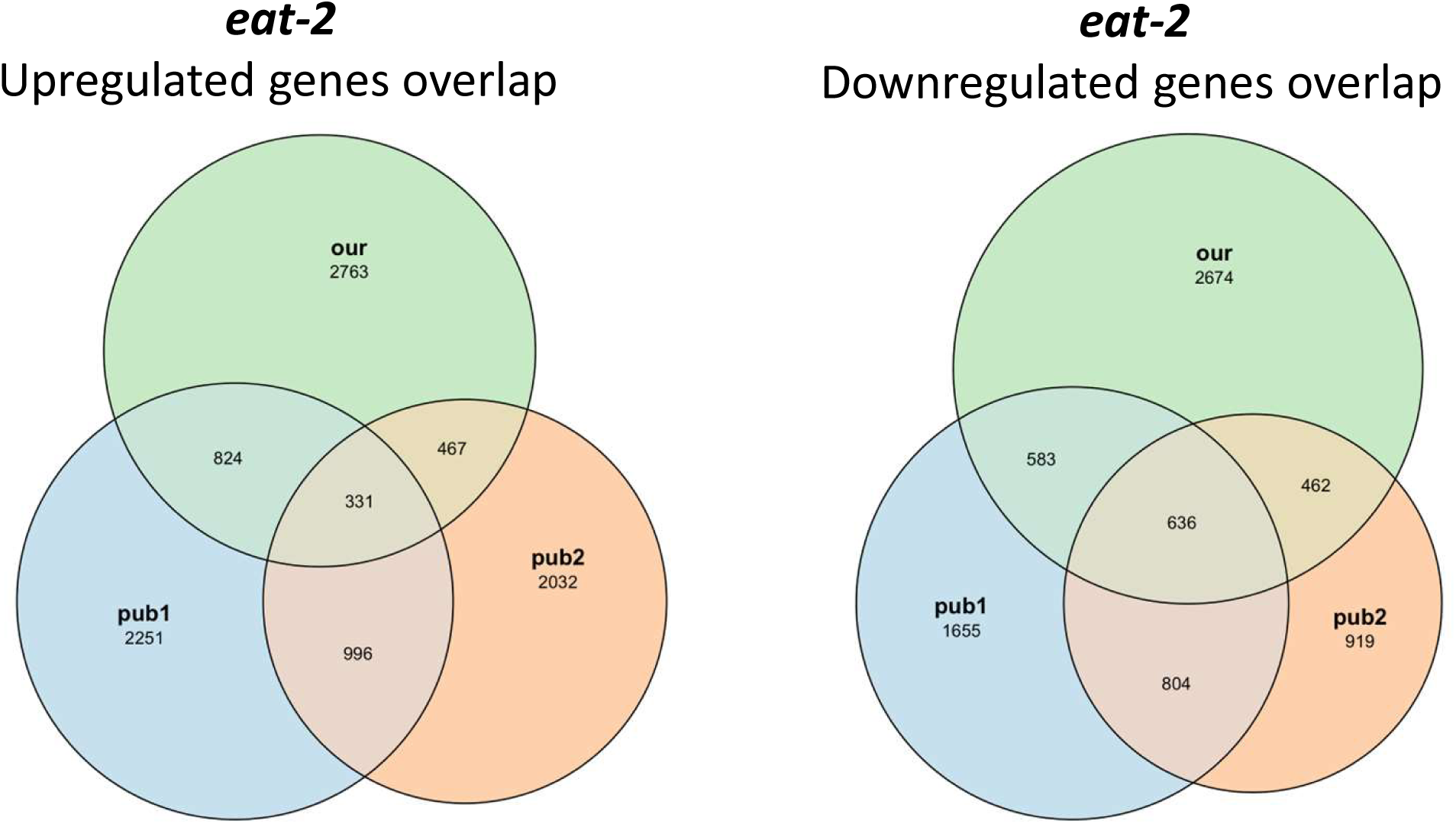
Comparison of gene expression changes in *eat-2* mutants in the current study to previous published gene expression results. The published RNA-seq data is from Dutta et al. 2025 *PLoS Biology* (https://doi.org/10.1371/journal.pbio.3003504) and Ng et al. 2020 *npj Aging (*https://doi.org/10.1038/s41514-020-0044-8).

**Figure S25.**
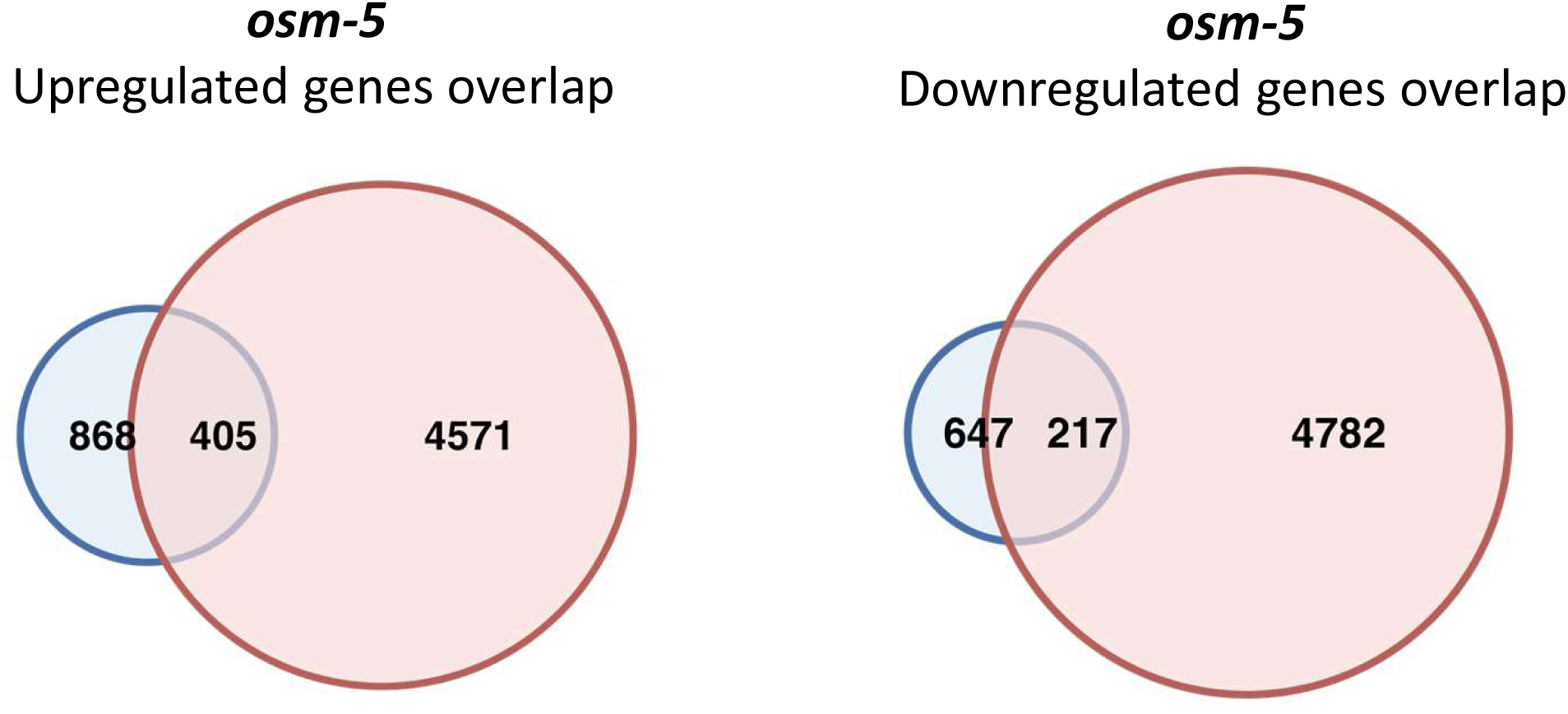
Comparison of gene expression changes in *osm-5* mutants in the current study to previous published gene expression results. The published RNA-seq data is from Li et al. 2024 *PNAS* (https://doi.org/10.1073/pnas.2321228121).

**Figure S26.**
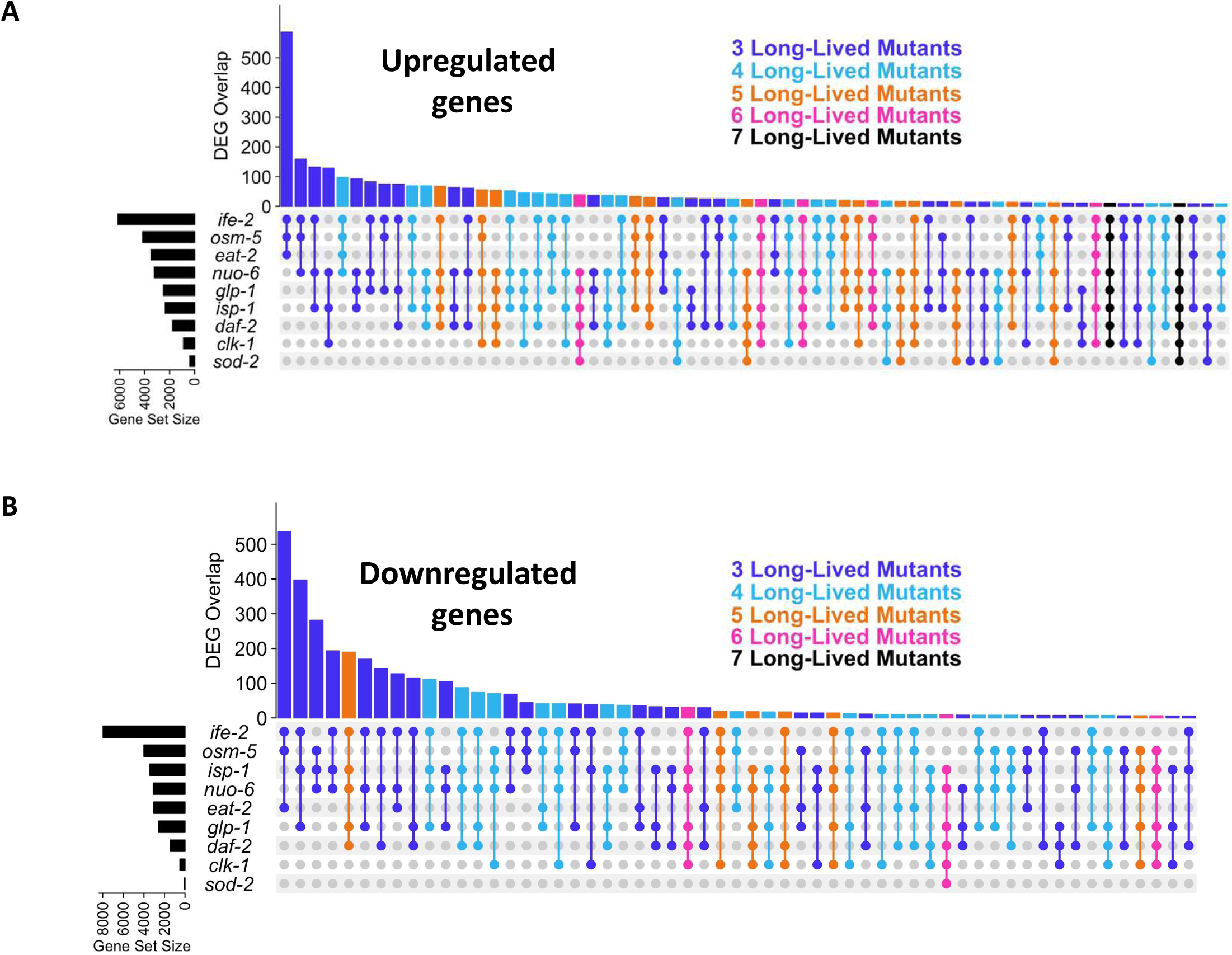
UpSetR plots showing degree of overlap between sets of long-lived mutants. (A) Significantly upregulated genes. (B) Significantly downregulated genes. Black bars on the left indicate the size of each gene set (number of differentially expressed genes in each long-lived mutant). Height of coloured bars indicates the number of genes in common between the strains indicated below. For example, the first column of the upregulated genes indicates the number of genes that are significantly upregulated in *ife-2, osm-5* and *eat-2* worms. The colours indicate the number of strains in each overlap: dark blue = 3, light blue = 4, orange = 5, pink = 6 and black = 7.

**Figure S27.**
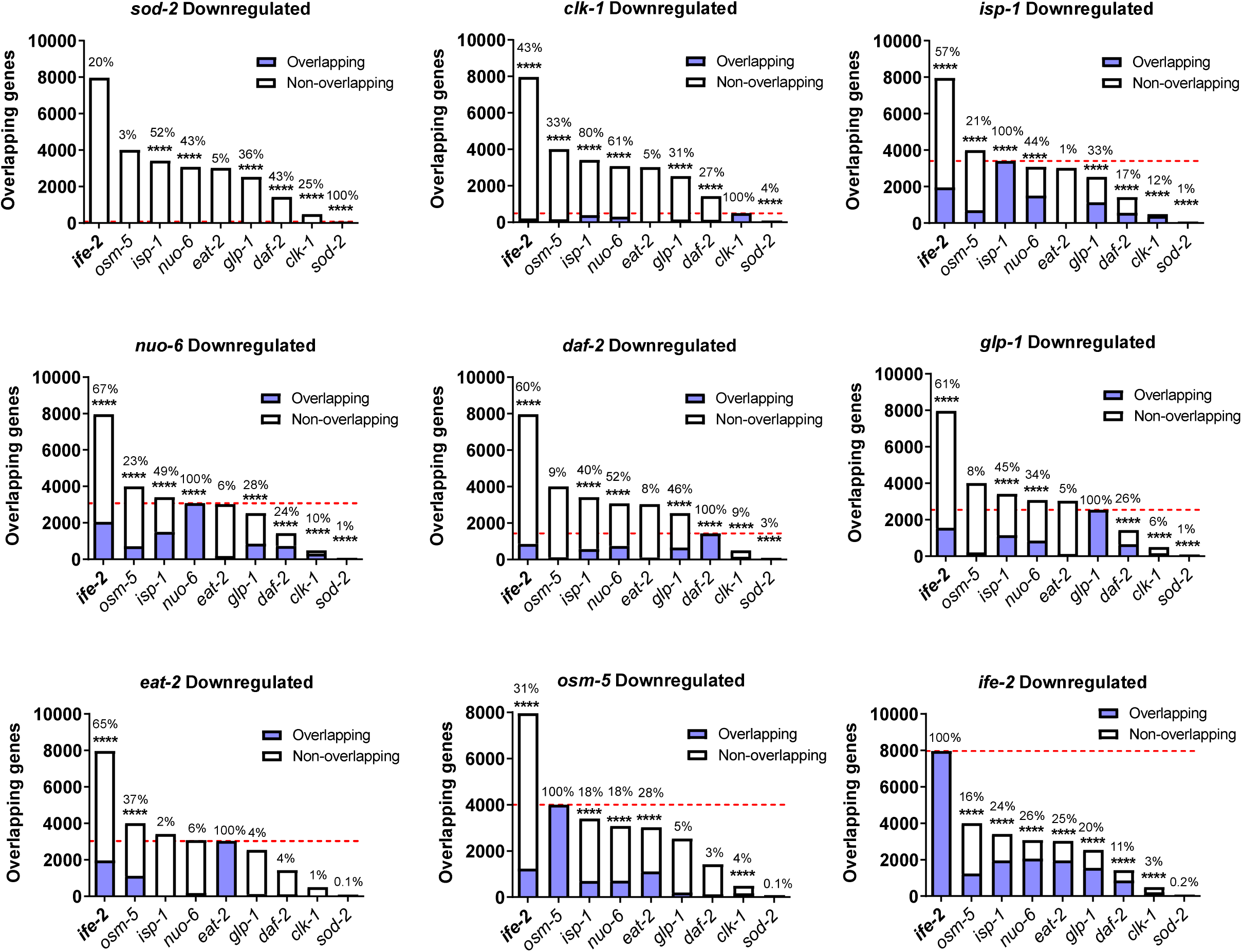
Degree of overlap of significantly downregulated genes between pairs of long-lived mutants is highly significant. Significantly downregulated genes (p<0.01) were compared between each pair of long-lived mutants. For most long-lived mutants, there is a highly significant degree of overlap between genes that are significantly downregulated in each long-lived mutant. For *eat-2* worms, there was only a significant overlap with *osm-5* and *ife-2* mutants. *ife-2* and *daf-2* mutants exhibited a significant degree of overlap with all of the other long-lived mutants. The percentage shown above each bar indicates the number of overlapping genes as a percentage of the total number of upregulated genes for the strain indicated in the graph title on top of each graph. Statistical significance was assessed using a Fisher’s exact test. ****p<0.0001.

**Figure S28.**
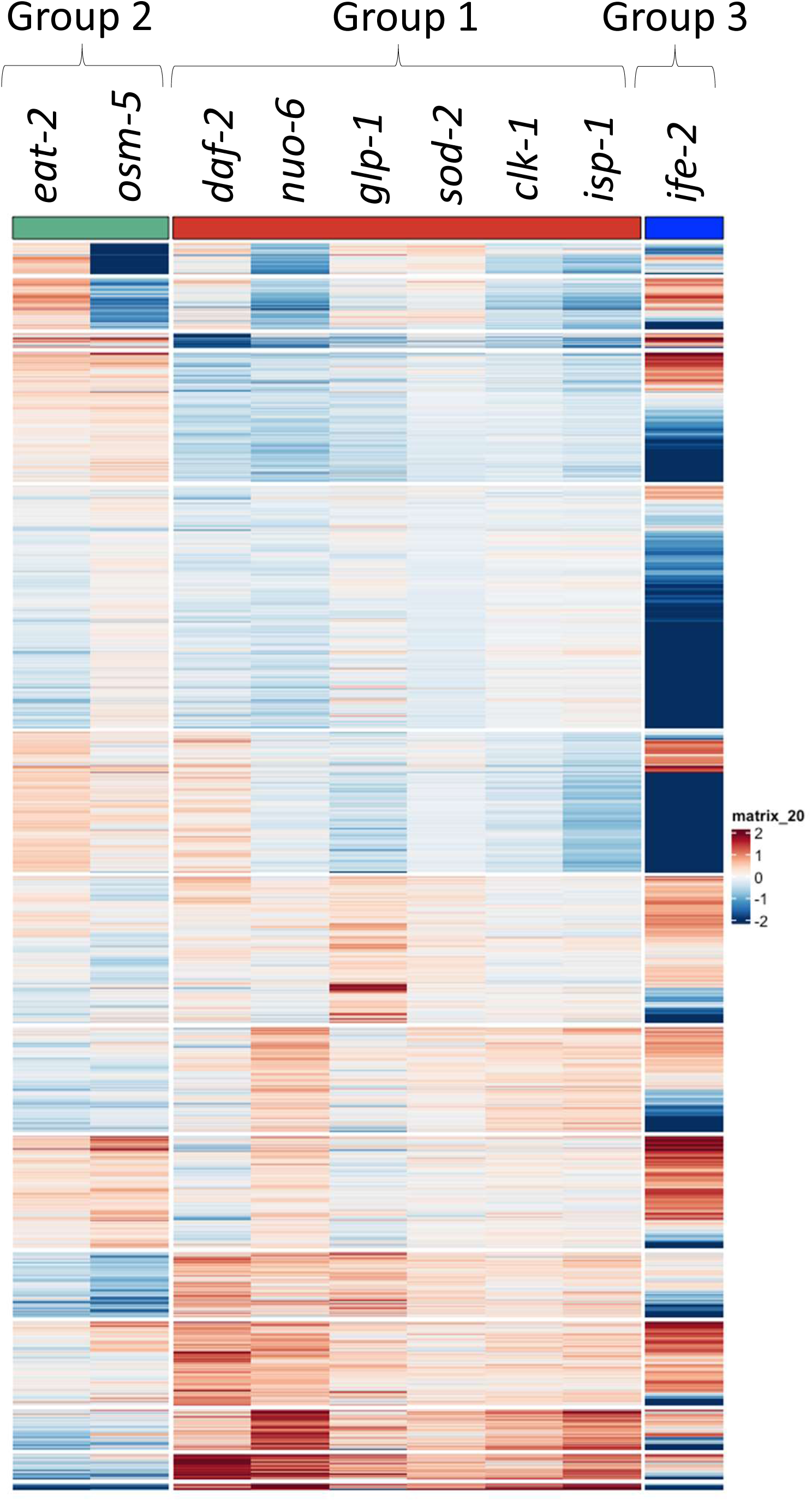
Heat map demonstrates that long-lived mutants cluster into three groups by gene expression. Group 1 contains *daf-2, nuo-6, glp-1, sod-2, clk-1* and *isp-1.* Group 2 contains *eat-2* and *osm-5.* Group 3 contains *ife-2*.

**Figure S29.**
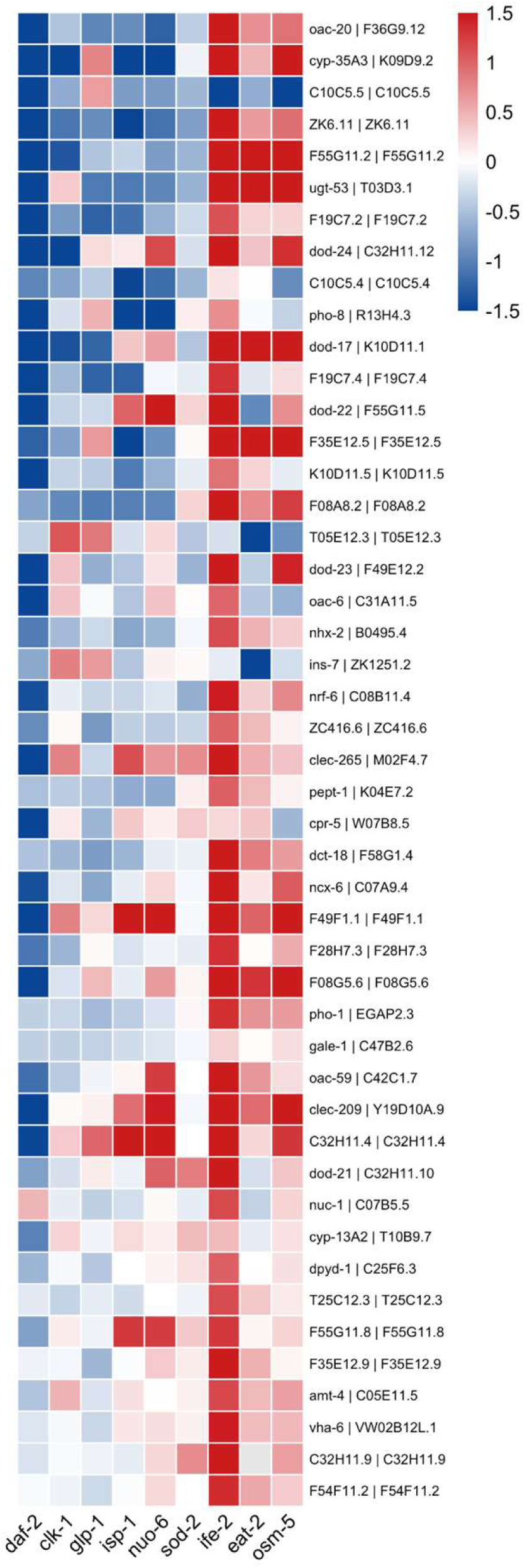
Group 1 longevity mutants exhibit downregulation of Class II DAF-16 target genes. A heat map of the top 50 high confidence Class II DAF-16 target genes (genes negatively regulated by DAF-16) from Tepper et al. Cell 2014 shows downregulation of genes that are negatively regulated by DAF-16 in group 1 mutants but not in group 2 or group 3 mutants.

**Figure S30.**
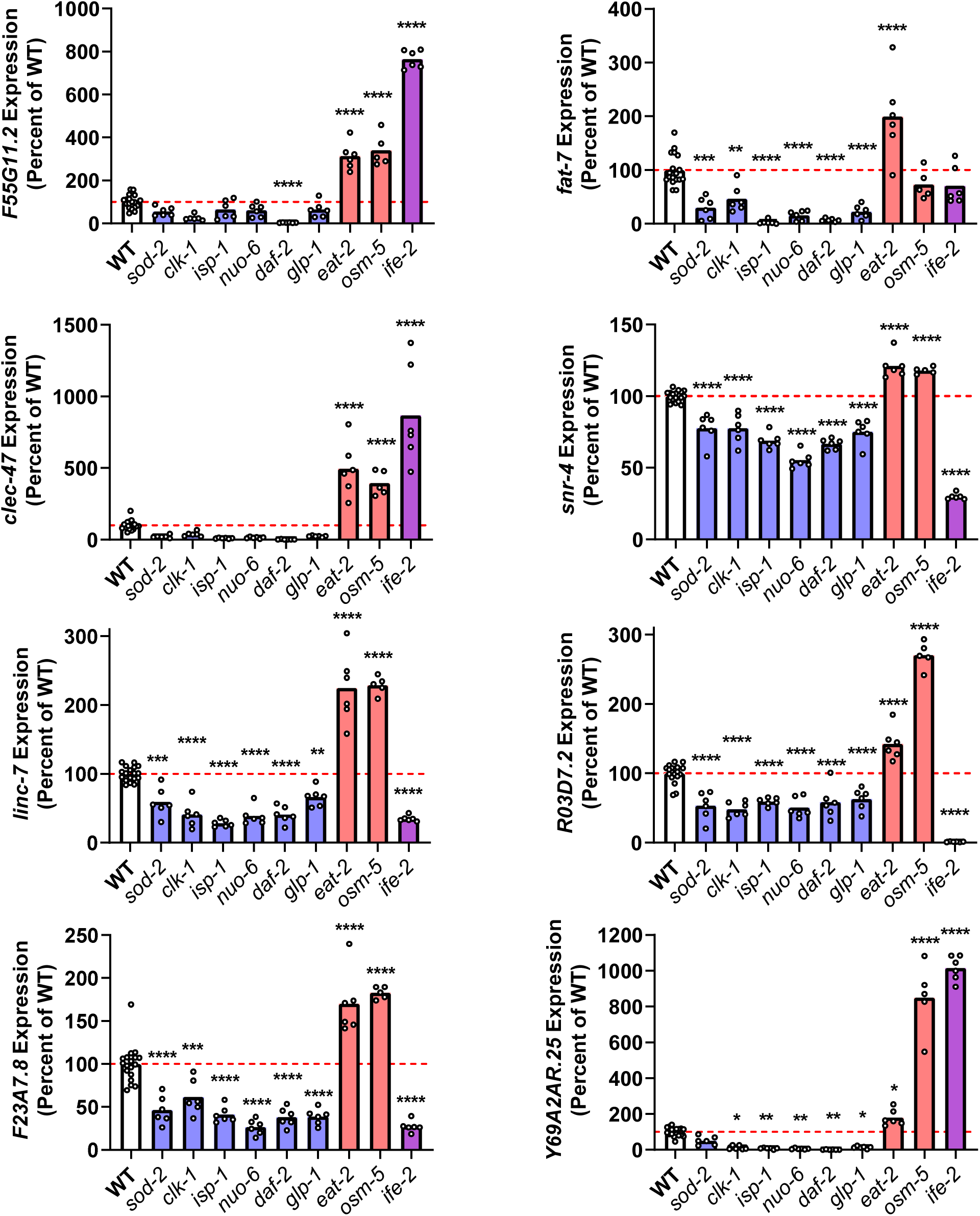
Examples of genes upregulated in group 2 longevity mutants. A number of genes were found to be significantly upregulated in group 2 longevity mutants *eat-2* and *osm-5* (red bars) but either unchanged or downregulated in group 1 longevity mutants (blue bars). These genes showed variable expression in the group 3 (purple bars) longevity mutant *ife-2.* Interestingly, among the genes that were specifically upregulated in group 2 longevity mutants were the PQM-1 target genes *F55G11.2* and *fat-7.* Statistical significance was assessed using a one-way ANOVA with Dunnet’s multiple comparisons test. **p<0.01, ***p<0.001, ****p<0.0001.

**Figure S31.**
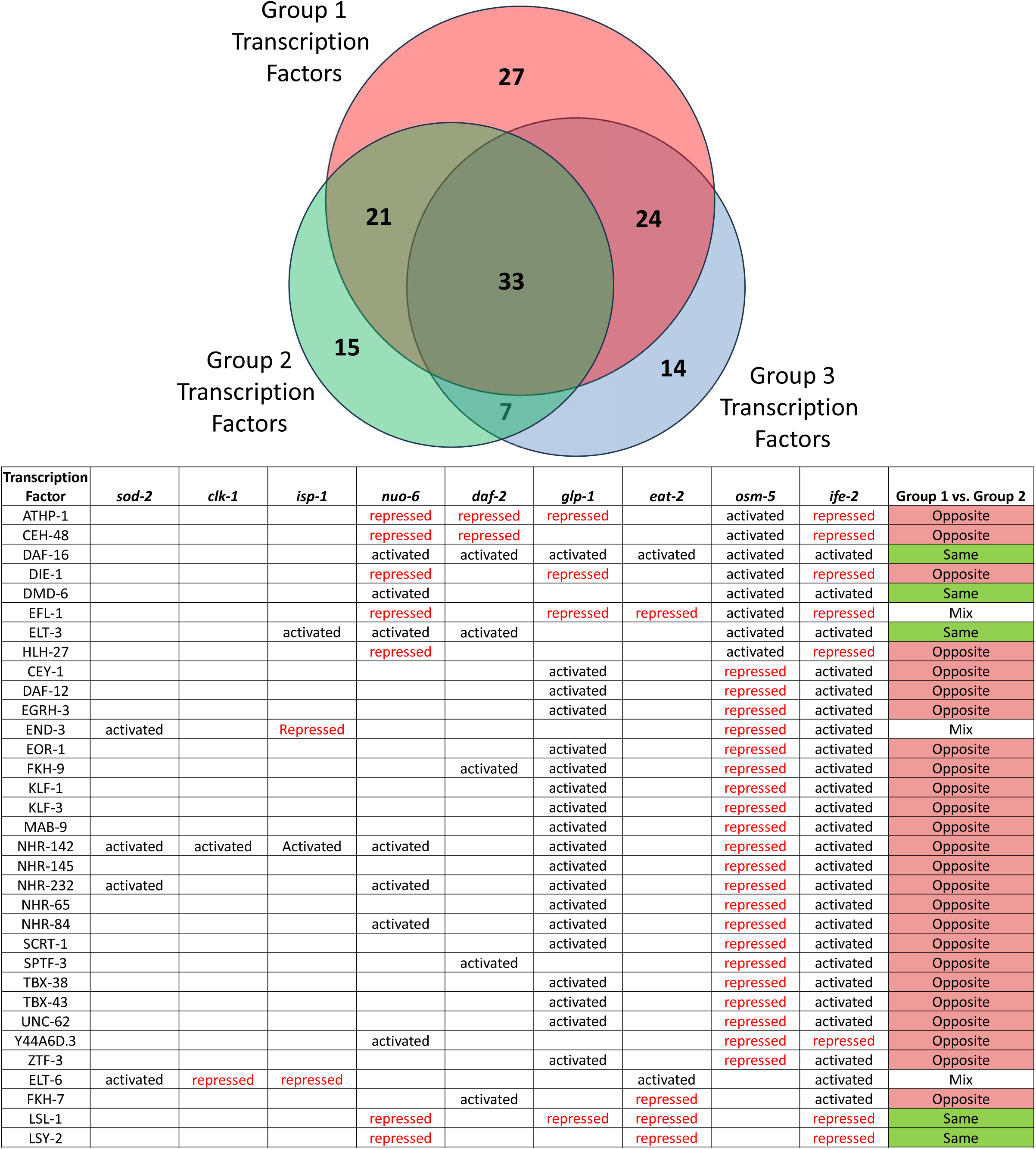
Activity of specific transcription factor are modulated in multiple long-lived mutants. The Venn diagram shows the overlap between transcription factor found to be modulated in group 1, group 2 and group 3 longevity mutants. There were 33 transcription factors that were found to be significantly modulated in all three longevity groups. Of those 33 transcription factors, 25 are modulated in opposite directions in group 1 and group 2 longevity mutants, while only 5 are modulated in the same direction. The direction of transcription factor modulation in *ife-2* worms is most similar to group 1 longevity mutants.

**Figure S32.**
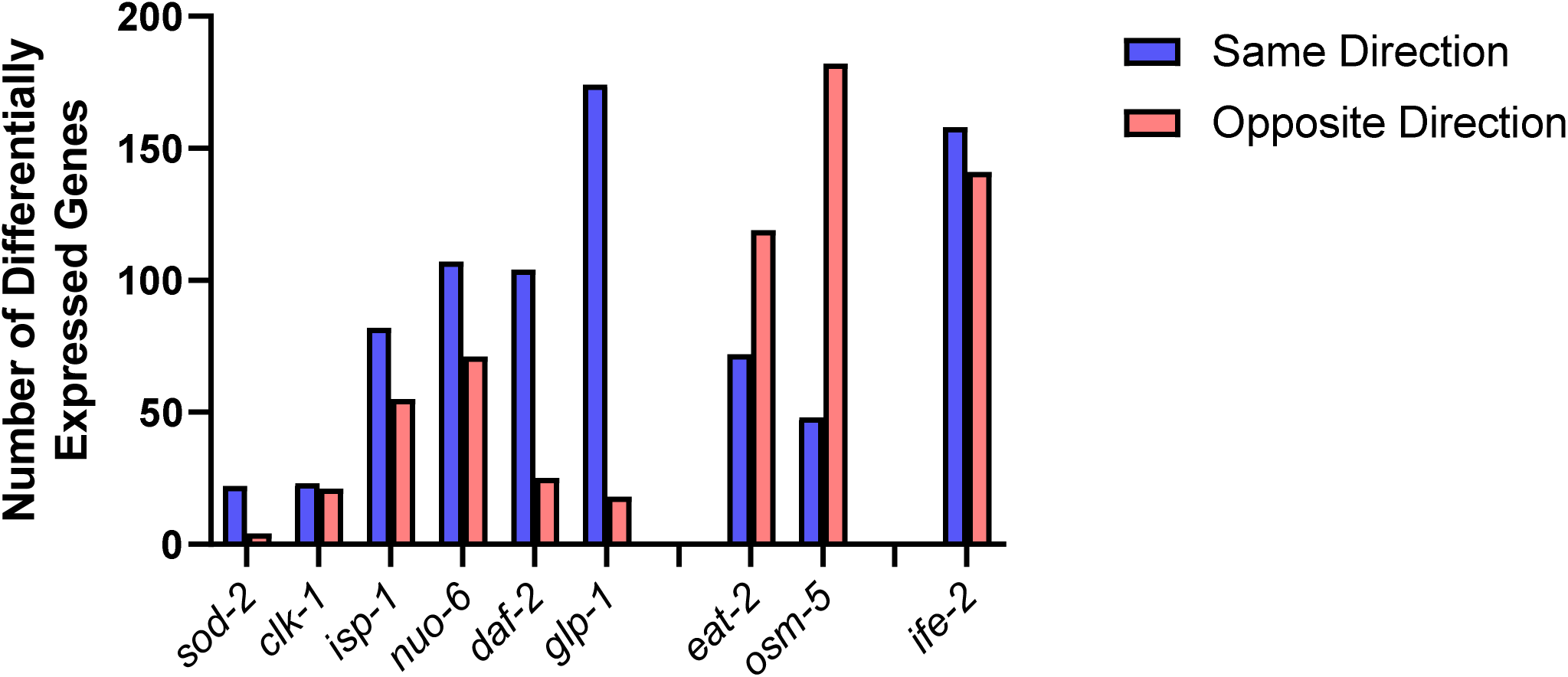
Comparison of gene expression changes in long-lived mutants to gene expression changes during aging. The gene expression changes in each of the nine long-lived mutants was compared to genes that are differentially expressed during aging. In each long-lived mutant, there were genes that were modulated in the same direction as changes that take place during aging (e.g. upregulated in long-lived mutant and upregulated during aging or downregulated in long-lived mutant and downregulated during aging) and genes that were modulated in opposite directions (upregulated in long-lived mutant and downregulated during aging or downregulated in long-lived mutant and upregulated during aging). Interestingly, in group 1 longevity mutants there were a greater number of genes modulated in the same direction as genes that are differentially expressed with age, while in group 2 longevity mutants there were a greater number of genes modulated in the opposite direction as genes that are differentially expressed with age. This was primarily due to genes that are downregulated during aging being upregulated in group 2 longevity mutants.

**Figure S33.**
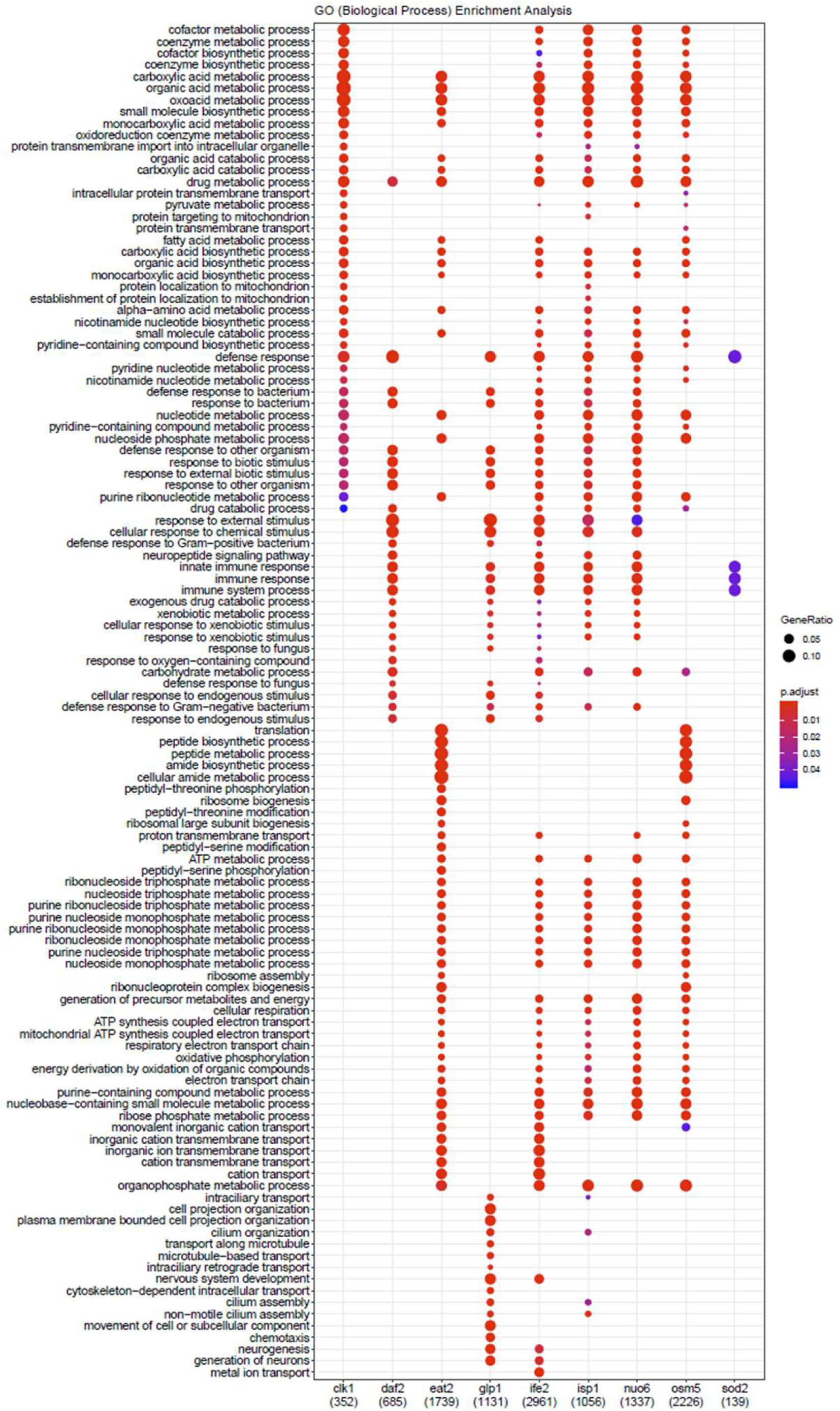
Genes upregulated in multiple long-lived mutants are involved in various metabolic processes, defense responses and innate immunity.

**Figure S34.**
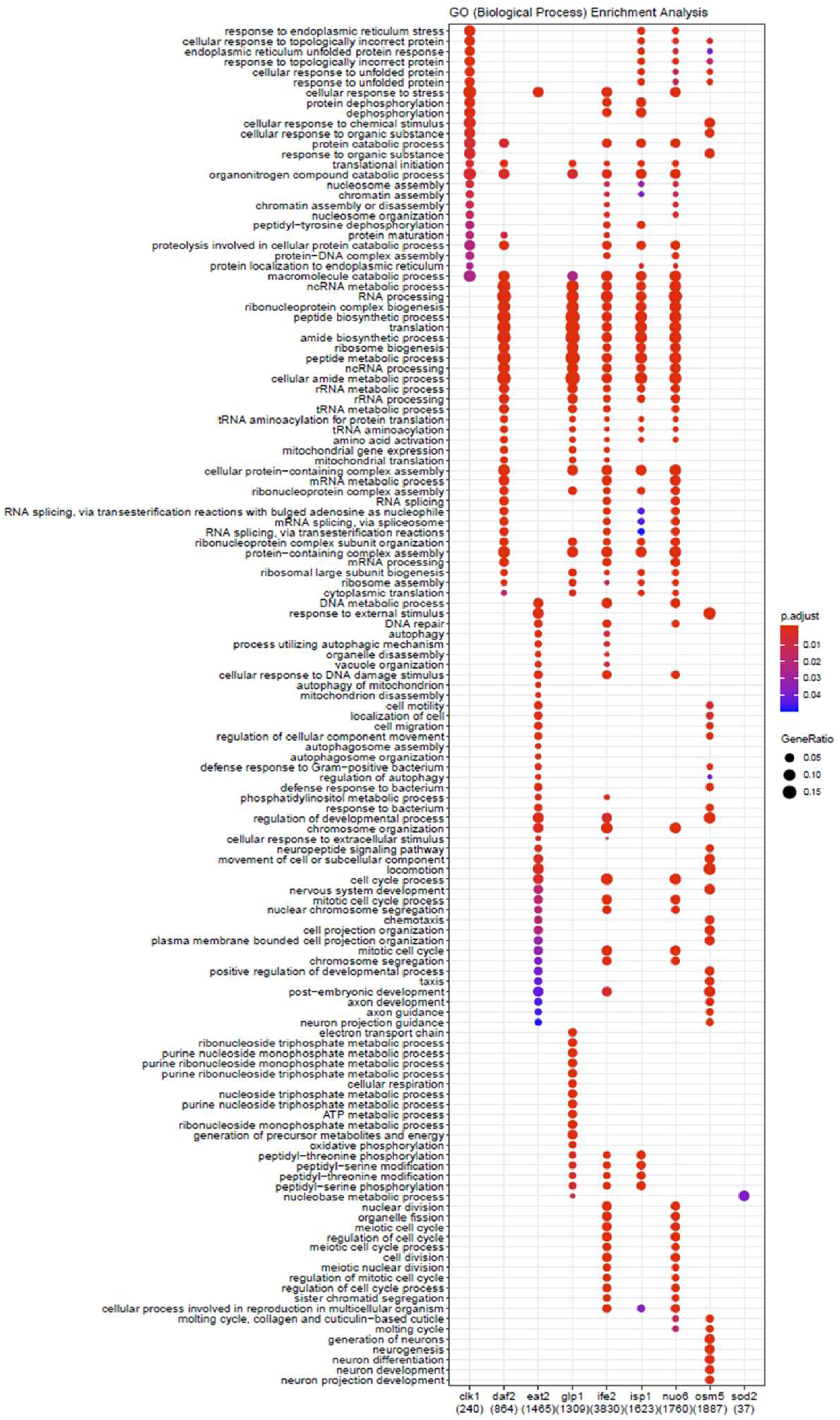
Genes downregulated in multiple long-lived mutants are involved in translation and various metabolic processes.

**Figure S35.**
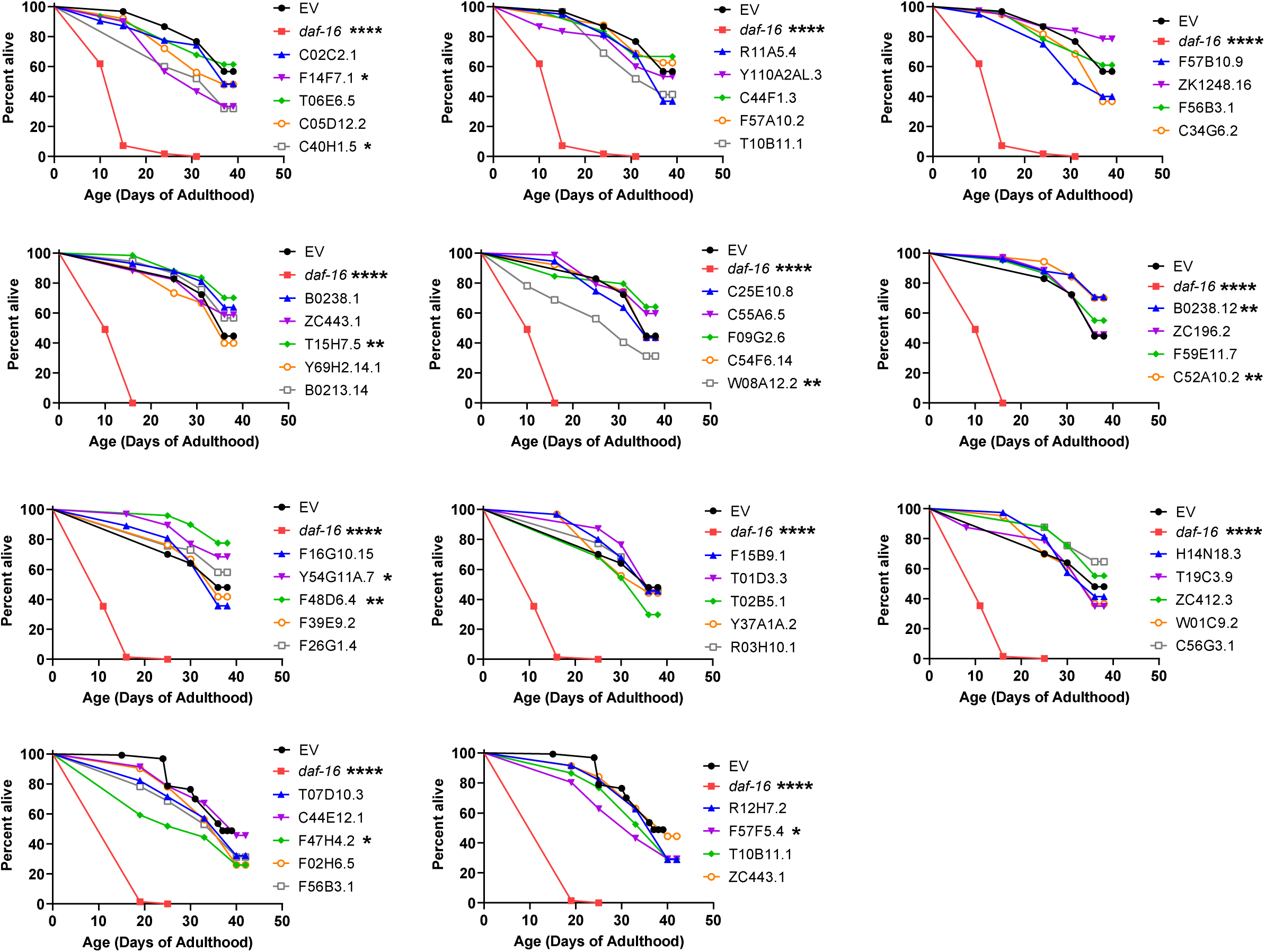
Survival plots for targeted RNAi screen in *daf-2* mutants. RNAi was initiated at the egg stage. *daf-16* RNAi was included as a positive control. Statistical significance was assessed using a log-rank test. *p<0.05, **p<0.01, ***p<0.001, ****p<0.0001.

**Figure S36.**
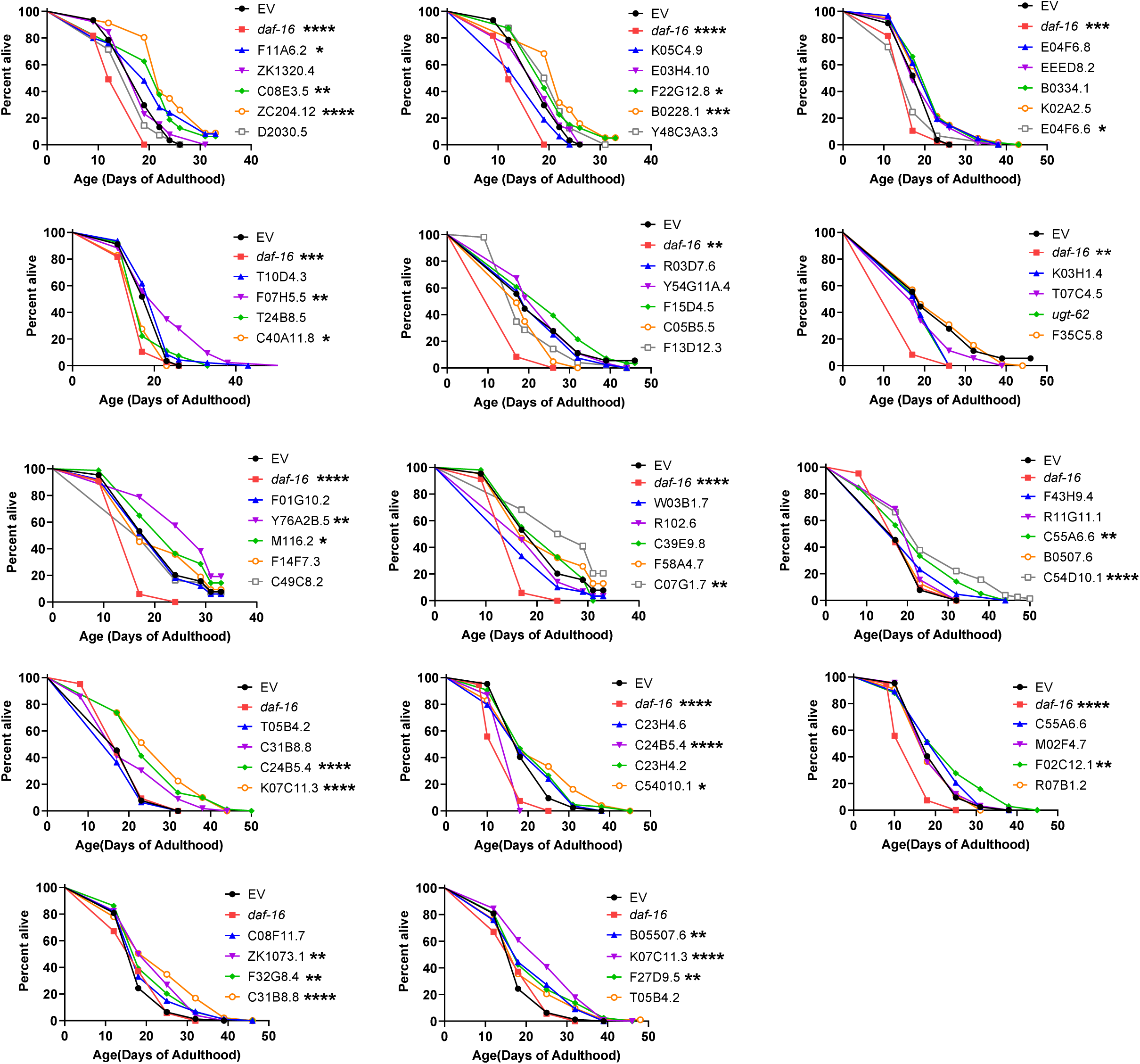
Survival plots for targeted RNAi screen in *nuo-6* mutants. RNAi was initiated at the egg stage. *daf-16* RNAi was included as a positive control. Statistical significance was assessed using a log-rank test. *p<0.05, **p<0.01, ***p<0.001, ****p<0.0001.

**Figure S37.**
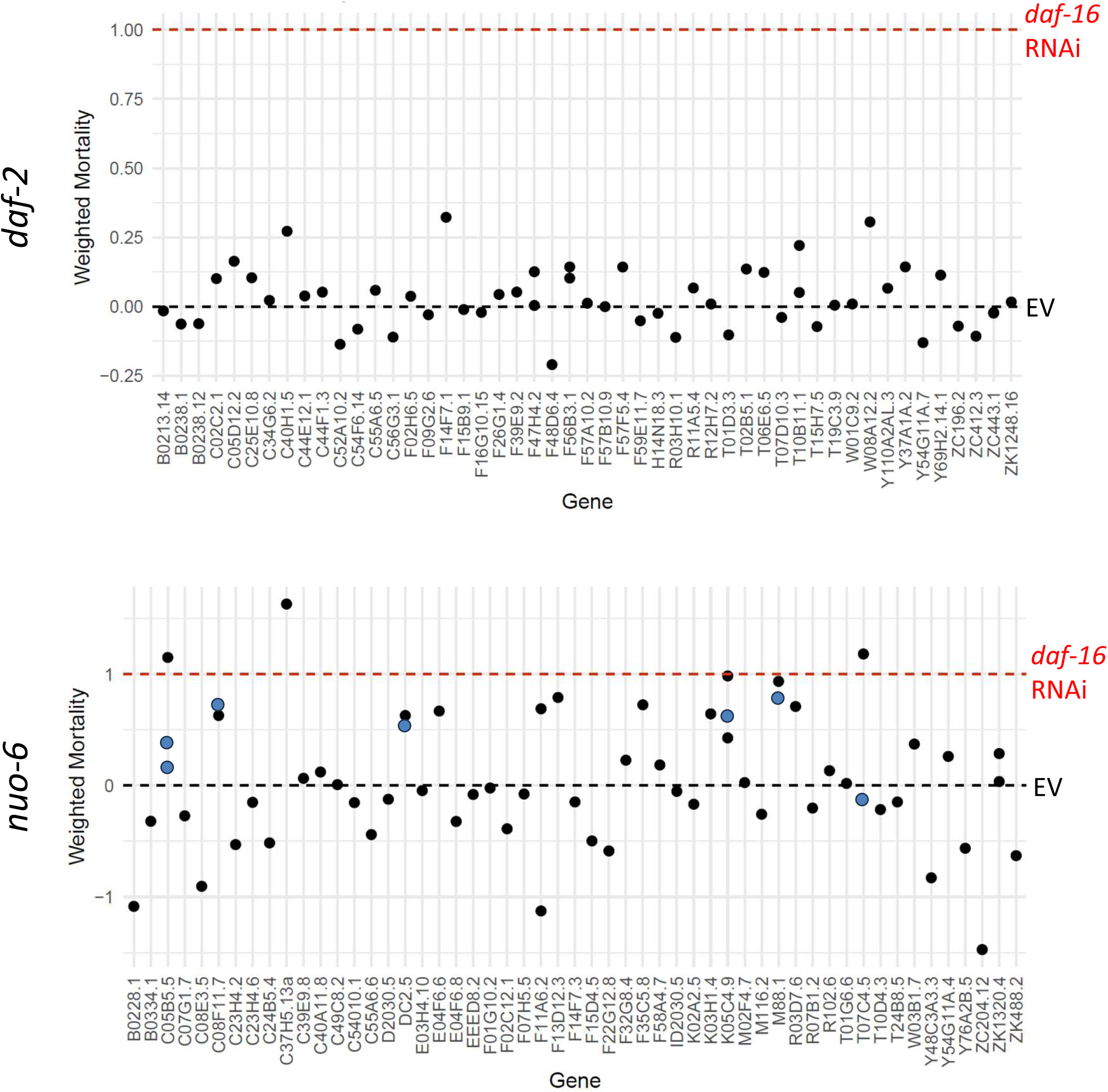
Summary of weighted mortality for targeted RNAi screen. RNAi was initiated at the egg stage. *daf-16* RNAi was included as a positive control. To compare the results of the RNAi lifespan screen across trials, weighted mortality was calculated as (mortality – mortality of EV)/(mortality of *daf-16* RNAi – mortality of EV). Black dots indicate data points from the initial screen. Blue dots indicate data from re-screening hits.

**Figure S38.**
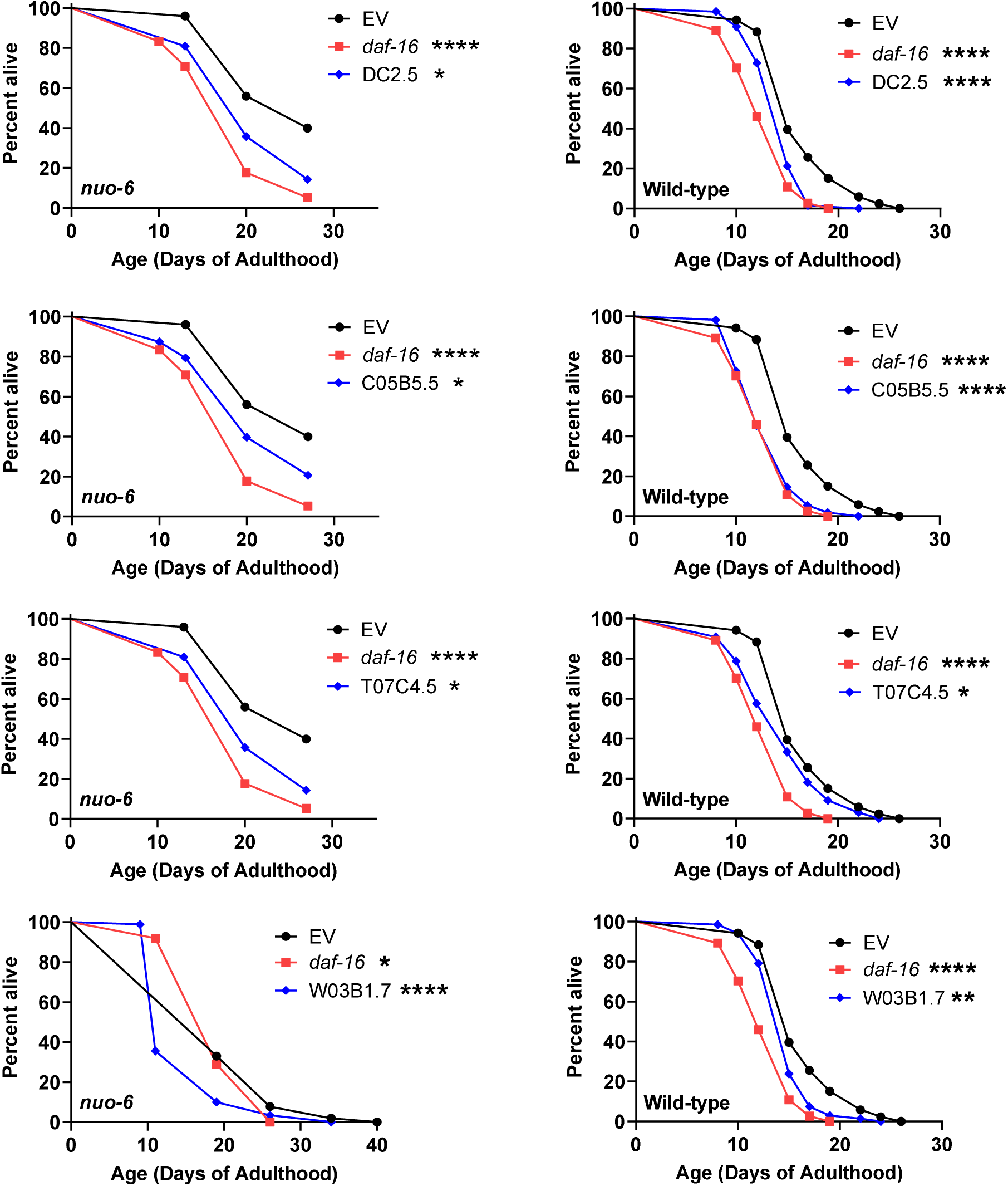
Repeat testing of RNAi clones that decreased *nuo-6* lifespan in initial screen. All four RNAi clones were found to decrease lifespan in both *nuo-6* and wild-type worms. Statistical significance was assessed using the log-rank test. *p<0.05, **p<0.01, ***p<0.001, ****p<0.0001.

## Notes

### Competing Interest Statement

The authors have declared no competing interest.

### Summary of Updates

A paragraph was added to the discussion that describes the role of DAF-16 in the longevity of long-lived mutants from the three different longevity groups.

